# Anesthesia unconsciousness recruits dynamic parabrachial activity within a distributed brain-wide ensemble

**DOI:** 10.1101/2025.03.28.646066

**Authors:** Kevin N. Schneider, Minke H.C. Nota, Hallie Lazaro, Daniel R. Rijsketic, Michelle Jin, Carlie Neiswanger, Asad Beck, Nick Ressler, Jason J.Q. Zhang, Amelia Li, Cole C. Shin, Ella Apley, Glorianna I. Gutierrez, Ainsley C. Barrow, Ian Campuzano, Alexandria D. Murry, Jovana Navarrete, Eric R. Szelenyi, Kentaro K. Ishii, Simon R.O. Nilsson, Garret D. Stuber, Christine A. Denny, Michael R. Bruchas, Boris D. Heifets, Horacio O. de la Iglesia, Sam A. Gloden, Mitra Heshmati

## Abstract

Neural circuits underlying unconsciousness remain poorly defined. We test the hypothesis that unconsciousness arises from specific, distributed circuits using general anesthesia in mice as a reproducible model. We identify a cortical-to-subcortical shift in neural activity during isoflurane anesthesia that is organized into nine discrete functional communities mapped at single-cell resolution. The lateral parabrachial nucleus (LPB) emerges as a central hub, exhibiting high interconnectivity, spontaneous firing under anesthesia, and preferential recruitment during reactivation of the brain-wide ensemble confirmed by single unit recordings. Chemogenetic reactivation of the captured brain-wide ensemble induces sedation, slow wave oscillations, hypothermia, and analgesia, which are components of anesthesia-induced unconsciousness. Reactivation of the LPB ensemble alone recapitulates a subset of these effects. Together, we define a global neural substrate for unconsciousness and recapitulate its dissociable autonomic, neurophysiologic, and behavioral effects using brain-wide ensemble manipulations. These results establish a neural circuit framework for anesthesia-induced unconsciousness in the mammalian brain.

**HIGHLIGHTS:**

- A distributed brain-wide ensemble comprises anesthesia unconsciousness
- Lateral parabrachial nucleus is a key, preferentially activated hub
- Brain-wide or parabrachial ensemble re-activation induces altered consciousness -

## INTRODUCTION

A prevailing hypothesis that general anesthesia produces unconsciousness by targeting a specific, controllable neural circuit substrate is yet to be confirmed. Many spatially segregated neural circuits are shown to regulate consciousness or recapitulate distinct components of an unconscious state^1,2^. General anesthesia likely encompasses neural circuit components of analgesia, sedation, immobility, and amnesia that together produce loss of consciousness^3^. Dissecting individual cell types and circuits contributing to these components has yielded valuable insights into the regulation of sleep^2,4–8^, arousal^9–16^, and anesthesia^17–25^, though with limited translatability for directly controlling consciousness.

The wet blanket explanation of anesthesia hypothesizes that anesthetic drugs work non-specifically to dampen excitatory signaling and enhance inhibitory signaling to generate unconsciousness^26^, or that anesthesia modulates mitochondrial metabolism in broad cell types^27^. Alternative hypotheses posit that specific neural circuits mediate anesthesia-induced unconsciousness^28^, such as specific cortical layer 5 decoupling^20,29^, or that anesthesia hijacks endogenous sleep and arousal-regulating pathways to produce unconsciousness^2,4^. Chemogenetic or optogenetic manipulation of cortical, thalamic, and hypothalamic nuclei, including the median preoptic nucleus (MEPO)^30^, as well as pharmacologic GABA_A_-R agonist infusions into the mesopontine tegmental anesthesia area (MPTA)^31^, have established multiple neural circuits important for loss-of-consciousness. A single, isolated neural correlate of unconsciousness has not been identified^32^. Here, we directly test the hypothesis that anesthesia-induced unconsciousness is subserved by globally distributed neurons in targetable circuits^33,34^. If this is true, a distributed, specific neural substrate of the anesthetized unconscious state exists that can be manipulated to modify conscious state transitions.

We apply a preclinical approach using intact-brain light-sheet fluorescent imaging^35–37^, activity-dependent targeted recombination in active populations (TRAP)^38,39^, chemogenetic activation^40^, high density single unit electrophysiology^41^, wireless mechano-acoustic recording of internal physiology^42^ and machine learning behavioral classification^43,44^ to manipulate and analyze cell-specific activity within the brain-wide isoflurane anesthesia unconsciousness network in mice.

Our approach better informs the neural framework of unconsciousness, as defined by general anesthesia, and investigates avenues for developing control of consciousness by targeting neural circuits.

While the parabrachial nucleus has an established role in arousal^14,45–48^, little is known about its contribution to generating unconsciousness. Our single-unit recordings across the brain indicate preferential recruitment of hindbrain activity in the brain-wide anesthesia ensemble that is driven by a lateral parabrachial (LPB) hub, also identified as a key community in the Fos activity map. We manipulate the LPB neural ensemble, revealing similar patterns of behavior as seen with brain-wide reactivation, including analgesia, freezing, hypothermia, and potentiation of the loss of righting reflex. Together, these studies establish a critical role for anesthesia-activated LPB ensembles in the unconscious state transition, as well as define the associated cellular resolution brain-wide network.

## RESULTS

### Single-cell resolution brain-wide activity map of isoflurane unconsciousness defines a cortical-to-subcortical shift in activity

To identify the neural circuitry of anesthesia-induced unconsciousness, we constructed an intact brain-wide map of activity following isoflurane (ISO) anesthesia using immediate early gene immunolabeling and light-sheet microscopy. Mice were anesthetized and maintained under 1.2-1.3% ISO for 180-min prior to fixation and intact brain processing^37^ (**Fig 1A**). Control mice were singly housed in the home cage. We analyzed whole-brain imaging datasets using two complementary pipelines, ClearMap^49^ and UNRAVEL^36^, which provide orthogonal views of Fos activity (**Fig 1B-C** and **Fig S1-S2**). ClearMap uses a cell-detection-based strategy to quantify Fos+ puncta after registration, producing both regional cell-density measures and voxelized statistical maps. In contrast, UNRAVEL prioritizes voxel-wise mapping of statistically robust differences in Fos immunolabeling intensity after background subtraction and z-scoring, then characterizes the anatomical composition and Fos+ cell densities of atlas-agnostic clusters of significant voxels (see **Methods** for a more detailed discussion). We followed these complementary analyses with network and community analysis of regional Fos maps (**Fig 2**)^50^.

**Figure 1.**
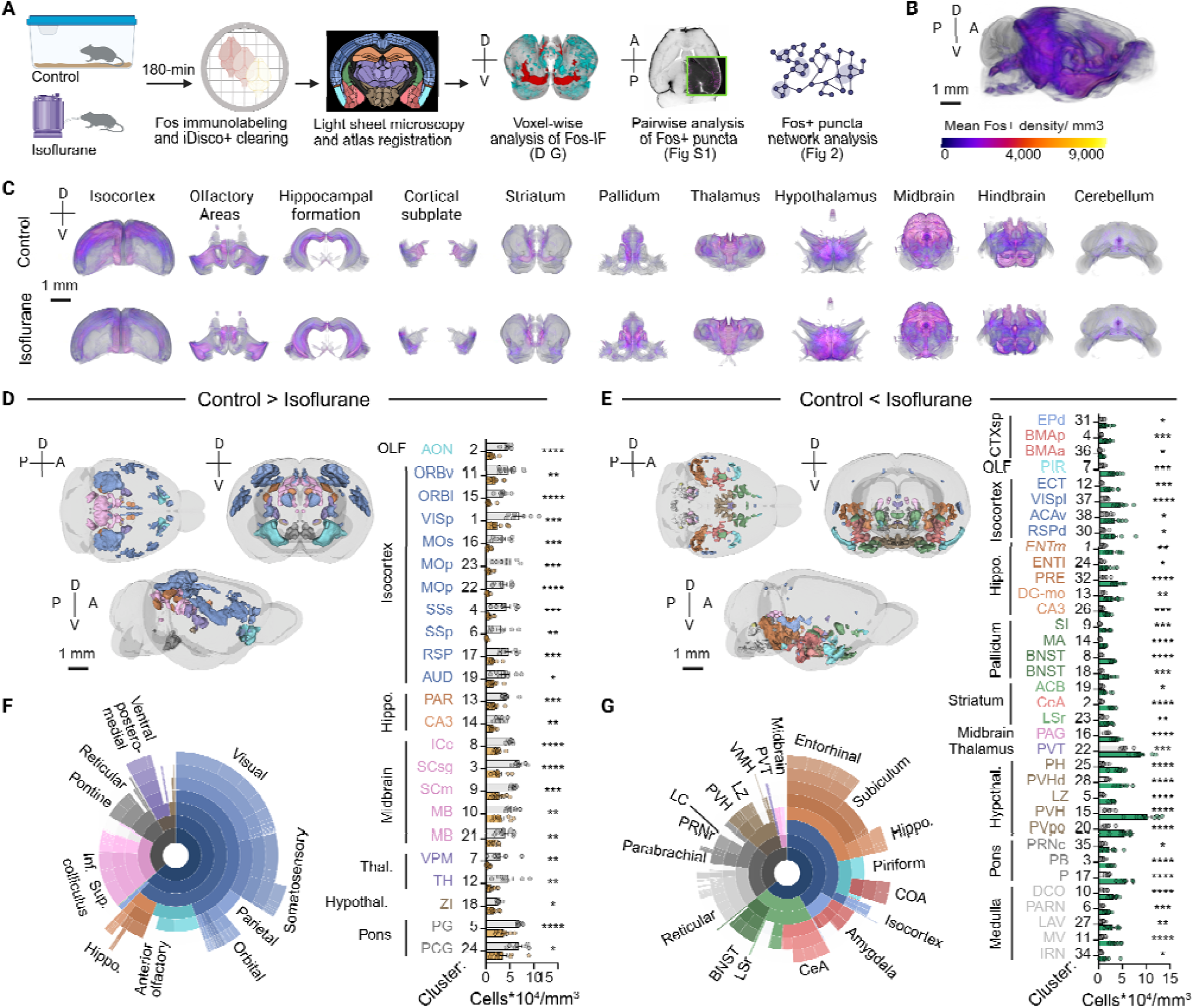
Single-cell-resolution map of isoflurane-activated brain-wide Fos expression. (A) Experimental overview: Three complementary pipelines were used to analyze whole brain Fos activation in home cage control and isoflurane-treated mice: voxel-wise immunofluorescence analysis with UNRAVEL (panels D-G), atlas-region-based Fos+ puncta quantification with ClearMap (Fig S1), and functional network analysis with SMARTTR (Fig 2). (B) Mean Fos+ density shown as a heatmap across the intact brain in all isoflurane samples (n = 9). (C) Mean Fos+ density heatmap across all control (n = 7, top row) and isoflurane samples (n = 9, bottom row), virtually sliced in the coronal plane at major anatomical subdivisions (see **Fig S1** for detailed ClearMap analysis and representative Fos images). UNRAVEL mapped regions where isoflurane either (D) suppressed Fos (q < 0.05) or (E) enhanced Fos (q < 0.005). FDR-corrected clusters differing in Fos+ cell density are shown in mirrored 3D brains and bar graphs (see **Video S1** and **Methods**). Data are shown as mean ± SEM, *p < 0.05, **p < 0.01, ***p < 0.001, ****p < 0.0001, unpaired two-tailed t-tests. (F-G) Sunburst plots summarize the anatomical composition of valid clusters by volume, with outer rings representing finer anatomical granularity in the Allen Mouse Brain Common Coordinate Framework (CCFv3) atlas hierarchy. Abbreviations are defined in **Table S1** with these exceptions: {’a’: ’anterior’, d’: ’dorsal’, ’l’: ’lateral’, ’v’: ’ventral’}. Raw Fos counts and z-scored ClearMap data are provided in **Table S1**. See Fig S2 and **Table S1** for details on each valid cluster and raw cell densities across multiple FDR thresholds. Scale bar, 1 mm.

**Figure 2.**
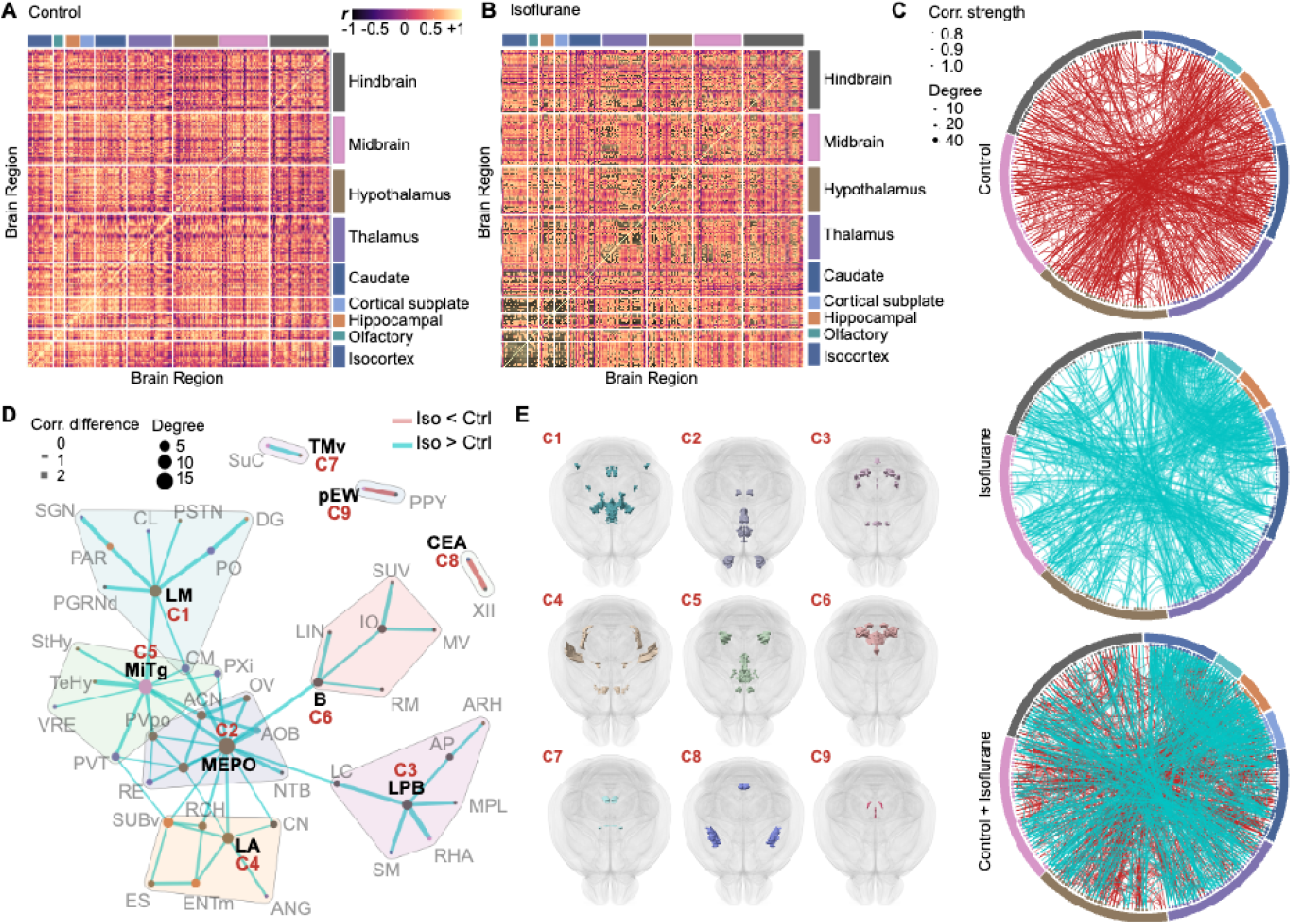
Network analysis reveals hubs that functionally differentiate an isoflurane unconscious state. Regional correlation heatmaps for (**A**) Control and (**B**) Isoflurane condition based on ClearMap atlas-registered mean Fos density data. Groupings of subregions by their larger parent regions are shown as sub-facets. (**C**) A combined network (bottom) showing the overlapping functional connectivity patterns of the individual control (top, red) and isoflurane (middle, blue) functional networks with matching network density (edge proportion = 3.15%) and the combined network below (see Fig S3 for proportional thresholding). (**D**) Functional difference network generated by plotting significant permuted regional correlation differences as edges (lines), and the regions they connect as nodes (circles). The uniquely shaded convex hull polygons denote regions classified as members of the same community. Key community nodes are abbreviated C1: LM = lateral mamillary nucleus; C2: MEPO = median preoptic nucleus; C3: LPB = lateral parabrachial nucleus, C4: LA = lateroanterior hypothalamic nucleus; C5: MiTg = microcellular tegmental nucleus; C6: B = Barrington nucleus; C7: TMv = ventral tuberomammillary nucleus; C8: CEA = central amygdalar nucleus; C9: pEW = pre-Edinger-Westphal nucleus. (**E**) Anatomical distribution of key community nodes identified in (**D**) and also shown in Video S2. See Fig S3 and Table S2 for all abbreviations and detailed statistics.

Using both approaches, we observe differential Fos patterns in the ISO state relative to control mice, with a shift from cortical to subcortical activation in the ISO-induced unconscious state (**Fig 1C**, **Fig S1B, Table S1**). When analyzed at a gross hierarchical level, the total mean Fos density is consistent across analyzed samples, significantly decreases with ISO in the isocortex and midbrain, and increases in the hypothalamic areas and cerebellum (**Fig S1G**; see **Fig S1C-D** for representative Fos images). At the granular level, ISO increases activity within subregions of the striatum (caudoputamen and central amygdala), pallidum (bed nucleus of the stria terminalis), thalamus (paraventricular nucleus), hypothalamus (paraventricular and ventromedial nuclei), and hindbrain (lateral parabrachial nucleus, locus coeruleus, and reticular nucleus) (**Fig S1H**). UNRAVEL confirmed these increases in activity (**Fig 1E-G**) and extended the findings to identify significant ISO-activated subcortical clusters that encompass multiple discrete brain subregions (see **Video S1** and **Fig S2**). For example, cluster 3 is predominantly in the LPB and neighboring pontine regions. Hotspots and coldspots from ISO-treatment are summarized in **Table S1**. Together, this shows that ISO activation occurs across several subcortical brain regions, with an overall cortical to subcortical shift in activity. These data establish a brain-wide neural framework for interpreting ISO-induced unconsciousness (**Fig 1**) and enable further network analysis (**Fig 2**).

### Network analysis of Fos maps reveals high interconnectivity of distributed communities

Since a given brain state is composed of dynamic interconnectivity between brain regions, applying network analysis approaches to study the structure of connectivity (i.e., network topology) can generate insights about the functional patterns that differentiate behavioral states^51–53^. We leveraged our brain-wide Fos mapping datasets to investigate how functional connections are changed by ISO-induced unconsciousness (**Fig 2**) using SMARTTR^50^, a software package supporting independently mapped IEG datasets for network analysis and visualization. We cross-correlated area-normalized Fos across all mapped subregions (**Fig 2A-B**) and found a higher number of strongly positive correlations in the isoflurane condition (**Fig S3**). Since Fos is an activity-dependent marker, positive correlations between brain regions represent either co-activation or co-suppression, while negative correlations represent opposing activation.

To better investigate which individual connections are most functionally altered by ISO, we subtracted the strength of each regional connection (i.e. correlation coefficient) in the control condition from its corresponding value in the ISO condition and compared each difference to a permuted null distribution (**Table S2**). Consistent with our cross-correlation analysis, there was a greater proportion of positively shifted connections in ISO (**Fig 2C**, **Fig S3**). We next constructed a network of the most significant functionally altered connections (p < 0.001), visualizing each correlation difference as an edge (lines), and the regions they connect as nodes (circles) (**Fig 2D**). This network revealed a naturally high interconnectivity amongst regions involved, visualized as a large, connected component (i.e., sub-network), rather than as collections of small, isolated networks or individual connections, as would be expected from chance (**Table S2**).

We also constructed individual networks (edge proportion 3.15%), to examine more general topological properties **(Fig 2C**, **Fig S3**). In the context of the brain, the property of small-worldness^54^ can be considered a balance between the capacity for functional specialization and integration. We confirmed that both the control and ISO networks exhibited small-worldness, defined as <7 > 1, although <7 was lower in ISO (3.61) compared to control (5.67) (**Fig S3D**). We examined global network properties across a range of alpha thresholds and proportional thresholds, finding the density of connections in the ISO network is consistently higher across a range of alpha thresholds, reflected in the higher trajectories of mean degree, clustering, and efficiency (**Fig S3B**). This effect is mitigated when controlling for edge density (**Fig S3C**). As a result, we found that global topological properties are highly influenced by greater functional interconnectivity in the isoflurane unconsciousness network.

Many complex real networks can be broken into subparts called communities, defined by nodes with high interconnectivity or probability of interconnectivity to other members within their community. We used an agglomerative community detection algorithm^55^ and discovered nine distinct communities within our functional difference network (**Fig 2D-E**). Metrics of nodal influence on a network, termed centrality, include degree (the number of direct connections to a node) and betweenness (how often a particular node lies on the shortest path between two other nodes). Examination of degree and betweenness (**Table S2**) revealed higher centrality of regions such as the MEPO, microcellular tegmental nucleus, lateral mammillary nucleus (LM), locus coeruleus, and LPB. Qualitatively, these metrics align well with the central, influential positioning of these regions within distinct, spatially distributed communities (See **Video S2**). Together, community detection identified nine distinct communities. The LM, MEPO and LPB hub communities were the most densely interconnected and covaried significantly with the ISO unconscious state (**Fig 2D**).

### Integrating behavioral analysis with internal dynamics during conscious state transitions to evaluate neural ensemble contributions

To investigate the contribution of this identified brain-wide network to ISO-induced conscious state transitions, we established an awake, behaving platform for concurrent assessment of dynamic internal physiology with automated behavioral classification during state transitions (**Fig 3**). We subdermally implanted mice with wireless, battery-free mechano-acoustic devices that detect heart rate, respiratory rate, physical activity, and body temperature, and that we previously validated for use with ISO^42^ (**Fig 3A**). Mice were then placed individually into a small modified open field chamber (15x15cm) that also permits ISO administration (**Fig 3A-B**). Behavioral state transitions were quantified with machine-learning-based supervised behavioral classification using pose estimation to capture rearing, grooming and immobility bouts in an automated, objective manner (**Fig 3C**)^43,44^. We established that the wireless mechano-acoustic devices capture a reproducible decrease in heart rate, respiratory rate, temperature and physical activity while mice underwent induction of ISO anesthesia (**Fig 3D, Video S3**) and their pose trajectories reflected reduced locomotor activity (**Fig 3E**). Using this behavioral platform, we went on to test the contribution of brain-wide ensembles to anesthesia-induced unconsciousness.

**Figure 3.**
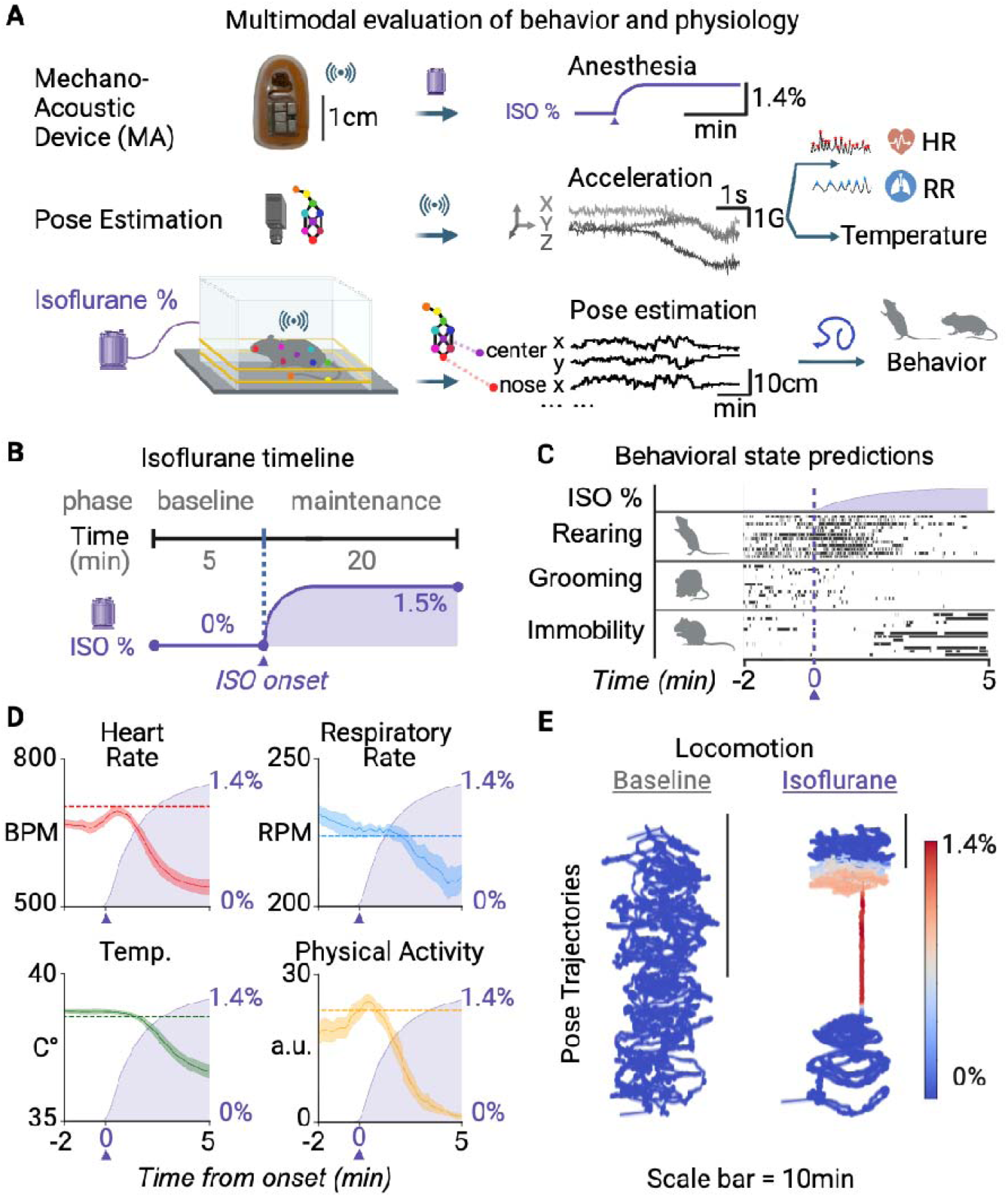
Integration of multimodal behavior and peripheral physiology during isoflurane state transitions. (**A**) Mechano-acoustic devices (MA devices) were implanted in mice to obtain internal physiological metrics (heart rate, HR; respiratory rate, RR; temperature & physical activity), integrated with pose estimation and isoflurane (ISO) gas concentration readout for a comprehensive assessment of internal state during isoflurane anesthesia. (**B**) We obtained physiological measurements as mice (n = 7) progressed through isoflurane induction. (**C**) Behavioral events detected by supervised classifiers using pose estimation. With increasing ISO concentration, mice gradually ceased grooming and rearing behaviors as they exhibited extended immobility. See Video S3 for representative integration of behavior and physiology. (**D**) Following induction (“ISO onset”), heart rate (“HR”; red), respiratory rate (“RR”; blue), temperature (“Temp.”; green) and physical activity (yellow) all decreased with increasing ISO concentration (%, violet), preceded by a brief rise only in HR and physical activity. Horizontal dashed lines show means of baseline trials (Means ± SEM; heart rate, 703.0 ± 2.26 bpm; respiratory rate, 223.8 ± 0.56 rpm; body temperature, 38.5 ± 0.01 °C; physical activity, 22.7 ± 0.38 a.u.). See Table S3 for detailed statistics. (**E**) 3-dimensional plot of pose trajectories reflecting locomotor changes with increasing isoflurane concentration. Color intensity represents ISO concentration from 0 to 1.4%. Scale bar is 10 minutes.

### Brain-wide activity-dependent capture of anesthesia unconsciousness

We hypothesized that if unconsciousness is regulated by discrete brain-wide-activated neurons, it should be recapitulated by activating the brain-wide neural ensemble. To capture brain-wide anesthesia-activated unconsciousness circuitry, Fos-TRAP2 (Fos2A-iCreER)^38^ mice received retroorbital injections of AAV-PHP.eB virus expressing Cre-dependent designer receptors engineered to be activated by designer drugs (DREADDs)^56^. After 3-4 weeks, mice underwent 180-min of ISO exposure (1.2-1.3%) and halfway through the exposure, at 90 min, received an intraperitoneal (i.p.) injection of 4-hydroxytamoxifen (4-OHT) to induce activity-dependent DREADD expression in brain-wide ISO-activated neurons (TRAP)^38,39^. On separate subsequent days, mice were implanted with either electrodes for electrocorticography (ECoG) or a wireless mechano-acoustic device^42^ to characterize neurophysiological and peripheral physiological effects, respectively, as well as behavioral changes after chemogenetic manipulation (**Fig 4A**).

**Figure 4.**
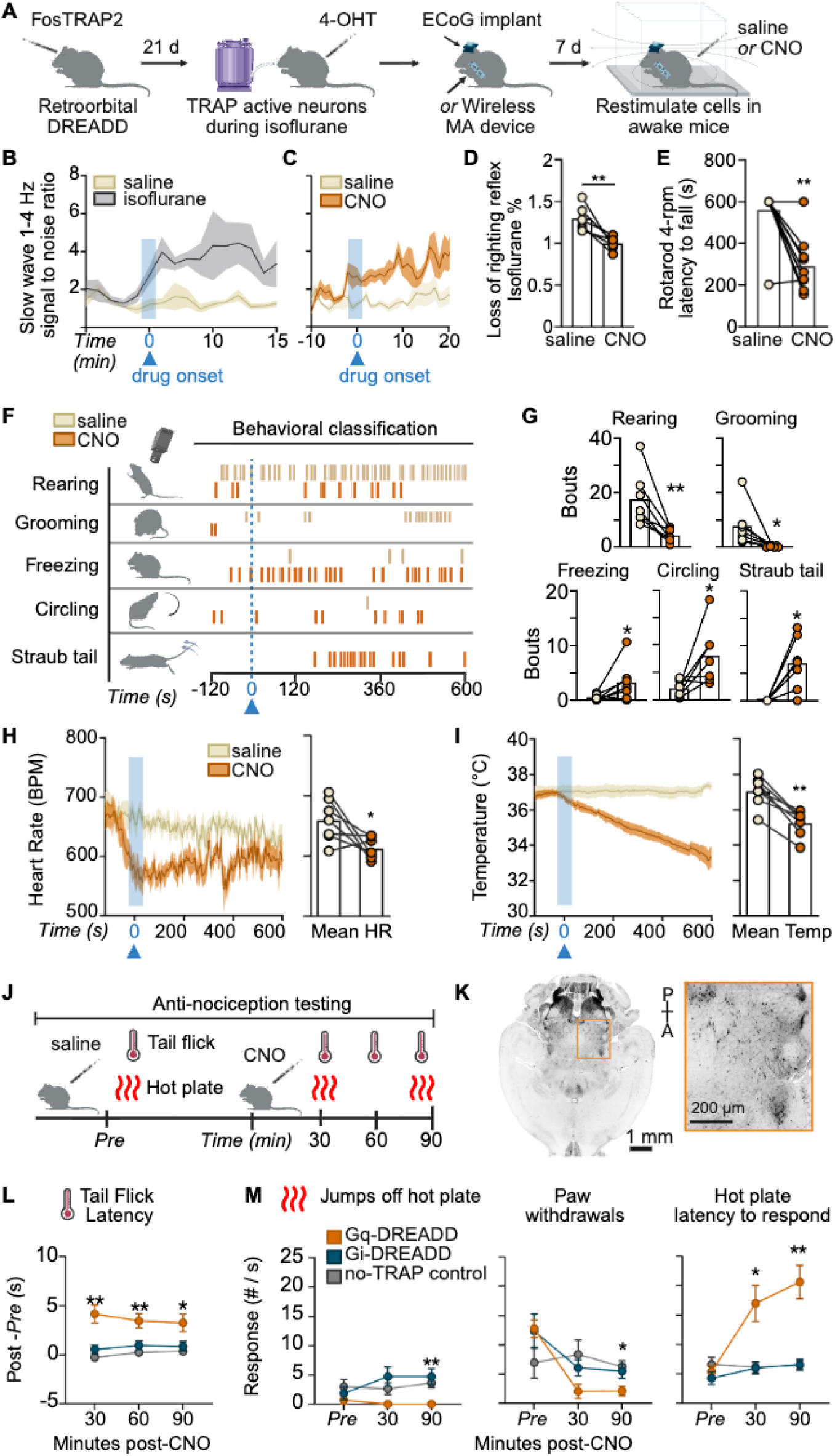
Brain-wide activity-dependent capture of anesthesia unconsciousness. (**A**) Experimental timeline: FosTRAP2 transgenic mice received retroorbital injection of AAV-PHP.eB Cre-dependent Gq-DREADD followed by activity-dependent capture (TRAP) during isoflurane exposure (1.2-1.3%) with 4-hydroxytamoxifen (4-OHT); (DREADD = designer receptor engineered to be activated by a designer drug). Mice were tested in the absence of anesthetic after either clozapine-n-oxide (CNO) or saline injection in a modified open field test with concurrent videorecording and peripheral physiology recording using wireless mechano-acoustic (MA) devices, or neural physiology with electrocorticography (ECoG). (**B**) 1-4 Hz delta slow wave activity is prominent after isoflurane induction and (**C**) after CNO injection as compared to saline in Gq-DREADD mice (n = 4, see Fig S4B for spectrograms). (**D**) Gq-DREADD expressing mice fall off a rotarod (4-rpm) faster after CNO injection (n = 10). (**E**) Gq-DREADD mice display loss of righting reflex at a significantly reduced isoflurane concentration after CNO injection, though this does not occur at baseline (see Fig S4C for dose-response and non-TRAP oil control). (**F**) A representative raster plot from all 6 (3 saline, 3 CNO) of one animal’s trials after analysis of classified behavior (also see Video S4). (**G**) Bouts of classified behavior for the average of all saline and CNO trials per animal (n = 7 mice, total of 18 CNO and 20 saline trials). (**H**) Left: Heart rate (beats per minute, BPM) decreases after CNO injection as compared to saline, shown time-aligned to drug onset, right: mean heart rate (HR) decreases after CNO injection in Gq-DREADD mice. (**I**) Left: Body temperature (Celsius) changes over duration of trial, time-aligned to drug onset, right: mean temperature decreases with CNO. See Fig S4 and Video S4 for Gi-DREADD data. (**J**) Experimental timeline for anti-nociception testing: FosTRAP2 mice received retroorbital injection of either Cre-dependent activating Gq-DREADD or inhibitory Gi-DREADD. They underwent isoflurane exposure, during which either 4-OHT or oil vehicle (if non-TRAP control) was administered. On test day, mice were tested for anti-nociception following saline injection (Pre) and 5 mg/kg CNO (Post). Warm water tail withdrawal (tail flick) latency was tested at 30-, 60-, and 90-minutes (min) post-CNO injection, and hotplate behavior was tested at 30-and 90-min post-CNO injection interleaved with tail flick tests. (**K**) Representative hindbrain light sheet image after intact brain iDisco+ clearing and mCherry-immunolabeling for DREADD expression (n = 7 mice). Scale bar is 1 mm and 200 µm in inset. See Fig 5 for brain analysis. (**L**) Gq-DREADD mice (n = 8, orange) show increased latency to tail flick compared to Gi-DREADD (n = 7, blue) or non-TRAP control mice (n = 8, gray) at every time point tested. (**M**) In the hot plate test, Gq-DREADD mice do not jump off the hotplate, have fewer paw withdrawals off the hot plate at 90-min, and show increased latency to first paw withdrawal response at 30 and 90-min post-CNO. 2-way repeated measures ANOVA with Tukey’s multiple comparisons: *p<0.05, **p<0.01. Paired t-tests: *p<0.05, **p<0.01. Error bars are the mean ± SEM. See Table S3 for detailed statistics.

After chemogenetic induction with the DREADD ligand clozapine-n-oxide (CNO, 5mg/kg i.p.), Gq-DREADD-expressing mice display increases in slow wave delta (1-4 Hz) oscillations as compared to their saline trials (**Fig 4B-C**, see **Fig S4** for spectrograms). We highlight a “drug onset” dashed line where the average onset of behavioral sedation occurs across CNO trials, approximately 5-10 minutes following injection. CNO induction promotes a significant increase in slow wave (1-4 Hz) signal to noise ratio similar to ISO induction (**Fig 4B-C**). Group-averaged spectrograms analyzed pre-and post-CNO exposure show this similarity between CNO and ISO induction, as well as a distinction from saline (**Fig S4B**).

Anesthesia-induced unconsciousness in rodents is commonly defined by loss of righting reflex (LORR)^57^, a postural reflex that allows rodents to regain the upright position. We find that CNO potentiates LORR with subanesthetic ISO administration, though LORR does not occur at baseline (**Fig 4E** and **Fig S4C**). A significant shift toward LORR occurs after chemogenetic stimulation of ISO-activated brain-wide circuitry in both CNO doses tested (5 and 10mg/kg) (**Fig S4A**). LORR potentiation does not occur in non-TRAP control conditions (**Fig S4C**) or in Gi-DREADD-inhibition of the brain-wide ensemble (**Fig S4F**). CNO (1 and 5 mg/kg) also produces dose-dependent reductions in movement velocity in activating Gq-DREADD-expressing mice (**Fig S4D**) but not inhibitory Gi-DREADD-expressing mice (**Fig S4G**). In behavioral tests, we continue to use 5mg/kg CNO, as this dose is validated to saturate DREADD receptors with minimal off-target effects^58,59^, does not modify behavior when administered to non-TRAP control and Gi-DREADD-expressing mice, and does not fully immobilize Gq-DREADD-expressing mice, allowing us to analyze behavioral repertoires of this altered state of consciousness.

We went on to examine behavioral effects after chemogenetic manipulation of the brain-wide ensemble in another cohort of mice. We assayed if coordinated movement, as measured in the rotarod test, is affected by chemogenetic stimulation of brain-wide anesthesia-activated circuitry. Mice underwent 2 days of training and were then placed on a slowly rotating rod (4 revolutions per minute, rpm) that poses minimal challenge to motor coordination under control conditions. After clozapine-n-oxide (CNO) injection (5 mg/kg), mice expressing activating Gq-DREADD fell off the rotarod when compared to their respective saline trials, indicating a loss of coordinated movement (**Fig 4D**), which is not observed in mice expressing brain-wide inhibitory Gi-DREADD (**Fig S4M**).

We observed that chemogenetic stimulation of ISO-activated brain-wide circuitry produces distinct behaviors, with differing temporal onset and distributions, when mice are tested and recorded in the modified open field chamber in the absence of anesthetic drug. An example raster plot of 6 cumulative saline and CNO trials collected from one Gq-DREADD-expressing mouse shows the representative distribution of 5 behaviors before and after drug onset (**Fig 4F**, also see **Video S4**). Using machine-guided supervised behavioral classification to quantify behavior relative to control saline injections, we find that CNO induction results in decreased rearing and grooming behavior, concomitant with an increase in stereotyped circling behavior, Straub tail, and freezing (**Fig 4G**). These effects were reproducible with each CNO trial across 4 separate cohorts of mice tested (see **Table S3** for detailed statistics).

Since altered consciousness may be associated with widespread physiological changes^60^, we analyzed peripheral physiological data collected from wireless mechano-acoustic devices that we previously validated for use under ISO. We aligned behavioral and physiological data to the average latency to the first bout of freezing observed across CNO induction trials (5 min after CNO injection, “drug onset”). Consistent with an altered state of consciousness, we observe significant decreases in body temperature and heart rate recorded after CNO induction (**Fig 4H-I**). CNO also increased mean respiratory rate in Gq-DREADD mice (**Fig S4E**).

In contrast, we did not see significant behavioral changes, besides decreased rearing, in brain-wide Gi-DREADD-expressing mice (**Fig S4H-I**). Mice that express Gi-DREADD never show the atypical behaviors, such as Straub tail, seen in the Gq-DREADD mice (**Fig S4H-I**). They also retain stable peripheral physiology after CNO induction (**Fig S4J-L**).

Since anesthesia-induced unconsciousness can encompass analgesia, we sought to identify the anti-nociceptive function of chemogenetic stimulation. Mice were tested for thermal anti-nociceptive responses in the warm water tail withdrawal (tail flick) test interleaved with hotplate testing after saline and at 30-min time points after CNO injection (**Fig 4J**). In the tail flick test, chemogenetic activation promoted significantly delayed tail withdrawal (**Fig 4L**). In hotplate testing, chemogenetic activation reduced jumping off the hot plate, decreased paw withdrawals, and increased latency to first response after CNO (**Fig 4M**). There was no effect of chemogenetic manipulation on thermal anti-nociceptive responding in non-TRAP control or TRAP Gi-DREADD mice (**Fig 4L-M**).

Following all behavioral tests, intact brains were cleared and immunolabeled for mCherry+ DREADD expression using iDisco+ and light-sheet microscopy to quantify brain-wide DREADD distribution (**Fig 4K** and **Fig 5**). Globally, we observe consistent brain-wide TRAP2 cell population density across gross cortical and subcortical divisions, besides significant differences in the hindbrain and cerebellum across samples analyzed (**Fig 5B**). To better understand brain region-specific differences, we used a voxel-based analysis where 3D and 2D TRAP2 density heatmaps exhibit clear patterns of region-specific TRAP2 enrichment (**Fig 5C-D**, **Table S3**). These expression patterns are not seen in non-TRAP controls that received oil vehicle instead of 4-OHT during TRAP (**Fig 5E-F**).

**Figure 5.**
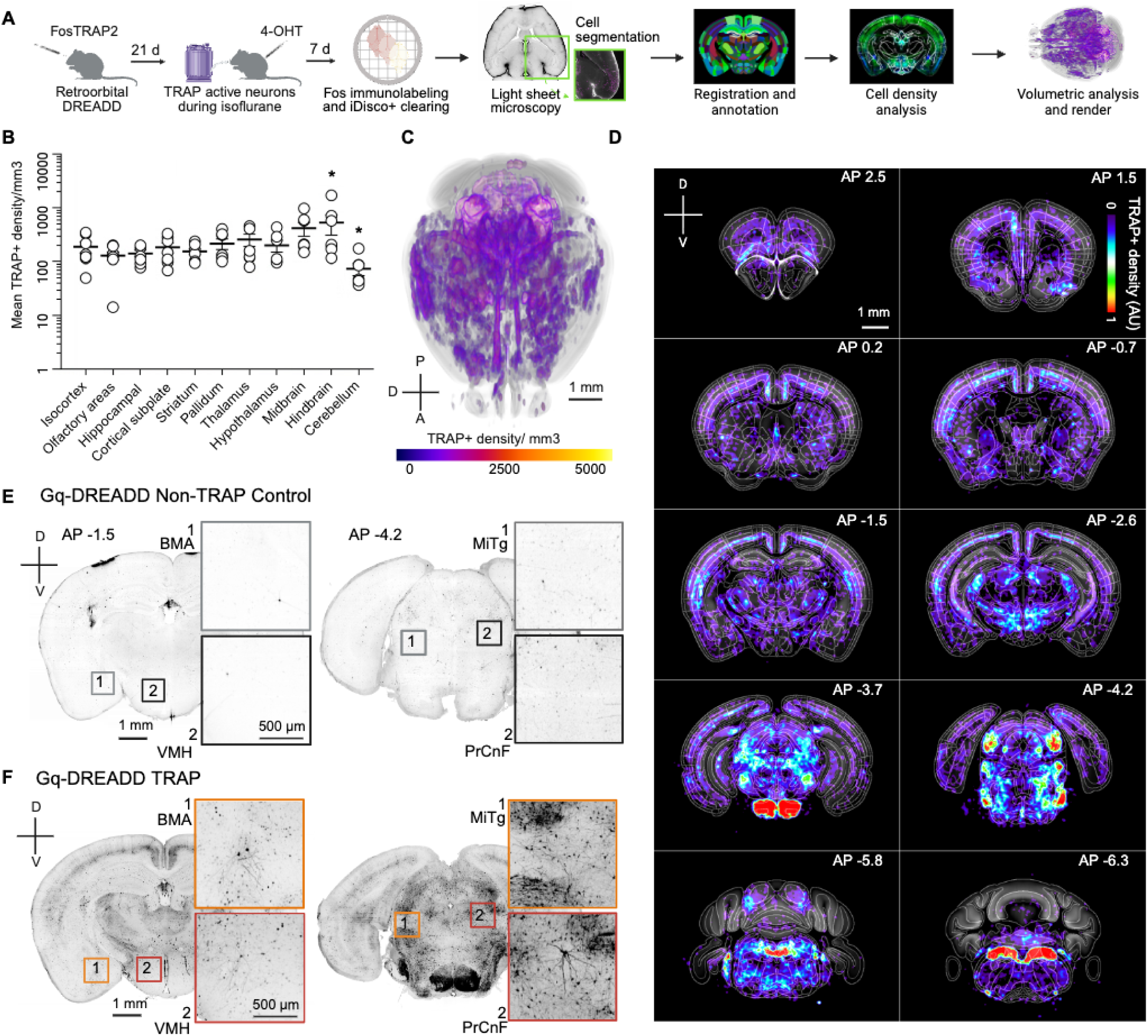
Spatial distribution of DREADD immunolabeling after brain-wide activity-dependent capture. (**A**) Experimental timeline: FosTRAP2 transgenic mice received retroorbital injection of an AAV-PHP.eB Cre-dependent Gq-DREADD followed by activity-dependent capture (TRAP) during isoflurane exposure with 4-hydroxytamoxifen (4-OHT). After experiments, intact brains were immunolabeled for mCherry-DREADD and cleared by iDisco+ (n = 7 mice). ClearMap was used to register brains, detect TRAP+ signal, generate cell counts and density heat maps. (**B**) Mean TRAP+ signal density calculated across larger brain subdivisions after atlas registration by ClearMap shows similar TRAP+ signal density across the brain, except for significant differences in the hindbrain and cerebellum (1-way ANOVA, *p<0.05). (**C**) 3-dimensional mean density heatmap of all samples (n=7). (**D**) Mean TRAP+ density across all samples shown as voxel-based 2D-projection heatmaps (calibration bar is arbitrary units). (**E**) Representative whole brain light sheet microscopy images after iDisco clearing and mCherry-DREADD immunolabeling digitally sliced to the 2D coronal plane from one control mouse expressing retroorbital AAV-PHP.eB Cre-dependent Gq-DREADD with oil vehicle administered during the isoflurane exposure (non-TRAP control, see **Methods**) and (**F**) one Gq-DREADD-expressing mouse that received 4-OHT during isoflurane exposure for activity-dependent capture of synthetic unconscious state. Abbreviations: BMA = basomedial amygdala, VMH = ventromedial hypothalamus, MiTG = microcellular tegmental nucleus, PrCnF = precuneiform area. Scale bar for full coronal slices is 1mm, inset scale bar is 500 µm. See **Table S3** for detailed counts and statistics.

By capturing the brain-wide anesthesia-activated neural ensemble, we describe the chemogenetic induction of an altered state of consciousness that mirrors patterns of behavioral, neural and physiological activity observed in natural sleep^61^, torpor^62,63^, and anesthesia^64^. This altered state is characterized by a significant increase in 1-4Hz slow wave oscillations, atypical freezing behavior, hypothermia, loss of coordinated movement, and thermal anti-nociception. Chemogenetic induction also potentiates LORR but does not generate LORR in the absence of anesthetic.

### Preferential single unit recruitment of isoflurane-responsive ensembles in the parabrachial nucleus of the hindbrain

Our previous experiments evaluated brain-wide cellular activation derived from static snapshots of Fos and DREADD expression maps. To examine the temporal dynamics of neurons engaged by ISO anesthesia, we used Neuropixels probes to record single-unit activity across multiple brain regions as mice underwent ISO exposure and chemogenetic reactivation of the captured brain-wide ensemble (**Fig 6**). Briefly, mice underwent surgery and were habituated to a custom head-fixed rig to facilitate acute Neuropixels recordings^41^ (**Fig 6A**, see **Methods** for details). In each session, we recorded activity starting with a 5-min awake baseline before ISO induction, after which mice were maintained unconscious under 1.3% ISO for 20-min (**Fig 6A**, “ISO”) before drug washout (25-min). After a second 5-min awake baseline, mice received CNO (5 mg/kg, i.p.) to re-active brain-wide neural ensembles (**Fig 6A**, “CNO”, 30 min). Dye-labeled probe trajectories were traced in light sheet images obtained from the intact, cleared brains of recorded mice (**Fig 6B**). A total of 2076 units from 8 mice were isolated and included for analysis (**Fig 6C**).

**Figure 6.**
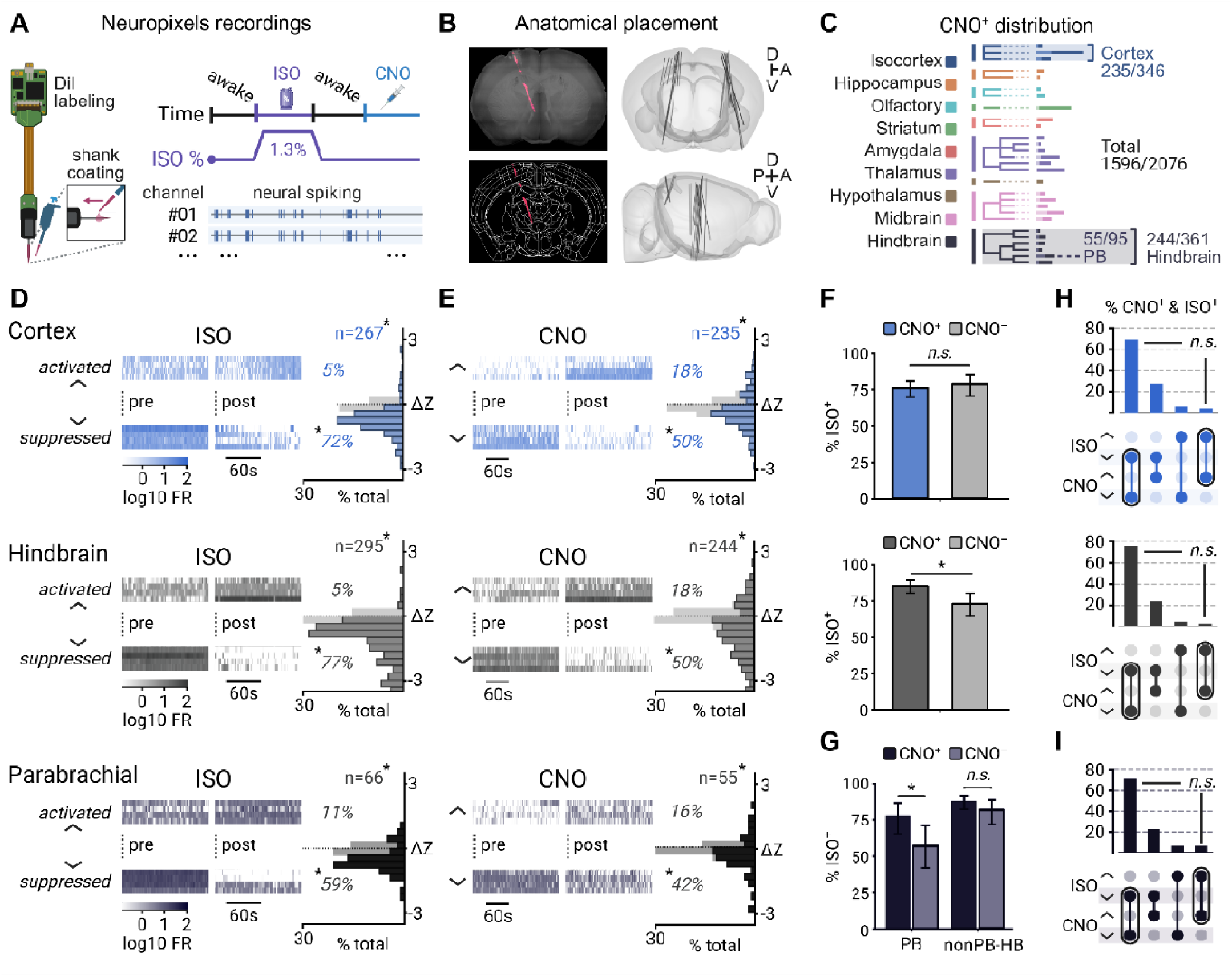
Brain-wide chemogenetic activation preferentially recruits isoflurane-activated neurons in the parabrachial nucleus. (**A**) Experimental schematic. Neuropixels probes (DiI-coated for histological tract reconstruction) were used to record single-unit activity from FosTRAP2 mice expressing Gq-DREADD, across a 5-min awake baseline and isoflurane induction (ISO, 1.3%) maintained for at least 20 min. This was followed by a 25-min wash and second 5-min awake baseline before delivering CNO (5 mg/kg) and monitoring chemogenetic activation of ISO-activated ensembles for 30 min. (**B**) Anatomical localization of recording probe tracts. Left: representative maximum intensity projection (top) across coronal depth (500gm) showing Dil probe tract (magenta), which are traced and aligned to anatomical regions in atlas space (bottom), to determine probe trajectories in the brain (right). (**C**) Anatomical distribution of recorded units by major region (color key) and the proportion responsive to CNO (darker bar color) per region; isocortex (Cortex, 235/346 CNO-responsive), hindbrain (244/361), and parabrachial nucleus (PB, 55/95) were targeted for analysis. Total CNO-responsive N = 1,596/2,076 total units obtained across 14 recordings, from 8 mice. (**D**) Left: representative firing-rate rasters (logD □ FR) for cortex (top), hindbrain (middle) and parabrachial nucleus (bottom) units significantly activated (top, “□”) or suppressed (bottom, “□”) by isoflurane, showing activity during “pre” (3 min leading up to onset) and “post” (3 to 6 min after onset) time periods. Right: distribution of single-unit response magnitudes as a change in their baseline-normalized logD □ firing rate (AZ, post -pre), as a % of total region population; gray bars denote neutral response distribution. (**E**) CNO-response panel as in D for cortex (top), hindbrain (middle) and parabrachial nucleus (bottom). Representative raster plots compare activity 5 min leading up to (“pre”) vs. 15-20 min after (“post”) CNO injection. ISO drove significant responsiveness above empirical null in all regions (CTX 267/346, 77.2%; HB 295/361, 81.7%; PB 66/95, 69.5%; q < 0.0001), with responsive units strongly biased toward suppression (of ISO-responders: CTX 248/267, 92.9%; HB 278/295, 94.2%; PB 56/66, 84.8%; q < 0.0001). Likewise, CNO engaged a significant proportion in all regions (CTX 235/346, 67.9%; HB 244/361, 67.6%; PB 55/95, 57.9%; all q < 0.0001), with a majority being similarly biased toward suppression among CNO-responders (CTX 173/235, 73.6%; HB 180/244, 73.8%; PB 40/55, 72.7%; all q < 0.0001). (**F-G**) Percentage of units that were ISO-responsive, computed separately within CNO-responsive (CNOD) and CNO-nonresponsive (CNOD) subgroups (mean ± Wilson 95% CI). (**F**) In cortex (top), ISO+ rates did not differ between CNOD and CNOD units (179/235, 76.2% vs. 88/111, 79.3%; OR = 0.84, q = 0.82), whereas in hindbrain (middle), CNO+ units overall were significantly enriched in ISO+ (209/244, 85.7% vs. 86/117, 73.5%; OR = 2.15, q = 0.007). (**G**) The enrichment within hindbrain was significant in LPB (43/55, 78.2% vs. 23/40, 57.5%; OR = 2.65, q < 0.05) but not in surrounding non-PB hindbrain (“non-PB-HB”). However, a direct PB vs. non-PB hindbrain comparison (CNO-responsiveness × region) was in the predicted direction but not significant (Firth-penalized likelihood-ratio test, OR = 1.60, p = 0.18). (**H-I**) Upset plots of joint ISO × CNO classification among double-responsive units, showing the percentage of each direction combination (activated / suppressed). Directional agreement (same sign response across ISO and CNO) did not exceed chance in any region (cortex 70.4%, q = 0.42; hindbrain 75.1%, q = 0.64; PB 74.4%, q = 0.32; nonPB-HB 75.3%, q = 1), with same-direction pairs being composed mainly of units suppressed by both. All p-values corrected using Benjamini–Hochberg FDR across 24 tests (6 per region), with significance reported as q values. See **Supplemental Fig S5** and **Table S4** for detailed statistics and data from additional regions. FR: firing rate; AZ: change in baseline-normalized firing rate; OR: odds ratio.

To determine whether CNO re-engaged the same units activated by ISO, we first classified each recorded unit by its responsiveness to ISO and CNO reactivation separately and evaluated their overlap within each region. We analyzed brain-wide responses across recorded regions and CNO-responsive proportions (**Fig S5, Table S4**). Across brain regions, CNO responses did not always maintain their original ISO-activated or suppressed response direction. Among units responsive to both ISO and CNO, a majority exhibited directional agreement. However, their proportion did not exceed chance in any region and same-direction pairs were overwhelmingly of joint suppression (**Table S4**). Thus, reactivation of the brain-wide ensemble re-engages units captured during ISO anesthesia and partially restores their anesthesia-state firing patterns.

At the population level, both major divisions, isocortex (n = 346) and hindbrain (n = 361), contained significant proportions of ISO-responsive and CNO-responsive units, exceeding shuffle null rates for each region (q < 0.001; **Fig 6D-E**, **Table S4** for detailed statistics). Responsive units were strongly biased toward suppression under ISO and CNO alike, consistent with broad inhibition that characterizes the anesthetized and chemogenetic re-activated states. Despite a shared suppressive signature, cortex and hindbrain diverged in whether CNO preferentially re-recruited the ISO-responsive ensemble (**Fig 6F, 6H**). In isocortex, CNO-responsive units were no more likely to be ISO-responsive than CNO-non-responsive units (OR = 0.84, q = 0.82), indicating that reactivation engaged cortical units independently of their past ISO responsiveness. In hindbrain, by contrast, ISO-responsiveness was significantly enriched among CNO-responsive units (OR = 2.15, q = 0.007), indicating preferential re-recruitment of the same units engaged during ISO anesthesia (**Fig 6F**). Together, this positions the hindbrain as preferentially re-activated relative to isocortex.

To determine whether preferential recruitment of the hindbrain was distributed across the region or localized to a specific node, we separated the LPB (n = 95), identified as a key hub in our whole-brain network analysis, from the surrounding non-PB hindbrain (nonPB-HB, n = 266). LPB showed significant preferential re-recruitment (**Fig 6G**), with CNO-responsive units enriched for ISO-responsiveness relative to CNO-non-responsive units (OR = 2.65, q = 0.036). The surrounding non-PB hindbrain did not reach significance (OR = 1.60, q = 0.17). Directly comparing the two subdivisions (CNO-responsiveness by region interaction) did not reach significance (Firth-penalized likelihood-ratio test, interaction OR = 1.60, p = 0.18), showing an effect in the predicted direction of LPB’s relatively greater re-recruitment (**Fig 6I** see also **Fig S5G**, **Table S4**). These results are consistent with the preferential single unit recruitment of a parabrachial ensemble that includes LPB as a key hub.

### Lateral parabrachial ensembles recapitulate altered consciousness when chemogenetically manipulated

Across our analyses, we discovered prominent activation of the LPB. LPB is significantly activated by ISO in Fos whole brain activity maps (**Fig 1**), emerges as a highly interconnected community (**Fig 2**) in network analysis of the Fos maps, exhibits neuronal firing under anesthesia and is preferentially recruited with chemogenetic stimulation after brain-wide activity-dependent capture (**Fig 6**). We hypothesized that if unconsciousness is regulated by discrete neural circuitry in the global network, unconsciousness should be recapitulated by activating highly significant, interconnected neural ensembles in the network, like the LPB. To capture ISO-activated LPB neural ensembles, we intracranially injected the same AAV-PHP.eB virus expressing Cre-dependent DREADD locally in the LPB of FosTRAP2 mice and subjected them to the same TRAP procedure under ISO, followed by behavioral testing with implanted wireless MA devices (**Fig 7A**). After behavioral tests, brains were processed with immunohistochemical labeling of mCherry-DREADDs observed predominantly at the LPB injection site, with some neurons also captured in pontine nuclei surrounding the injection site (**Fig 7B** and **Fig S6L-M**). Mice underwent alternating trials of saline and CNO with concurrent videorecording in the modified open field box. We quantified behavior and peripheral physiology using the same automated classification pipeline as with brain-wide ensemble chemogenetic manipulations to evaluate the contribution of the LPB ensemble to altered consciousness. A representative plot showing the behavioral classification from the average of one animal’s saline and CNO trials is shown in **Fig 7C** (also see **Video S5)**.

**Figure 7.**
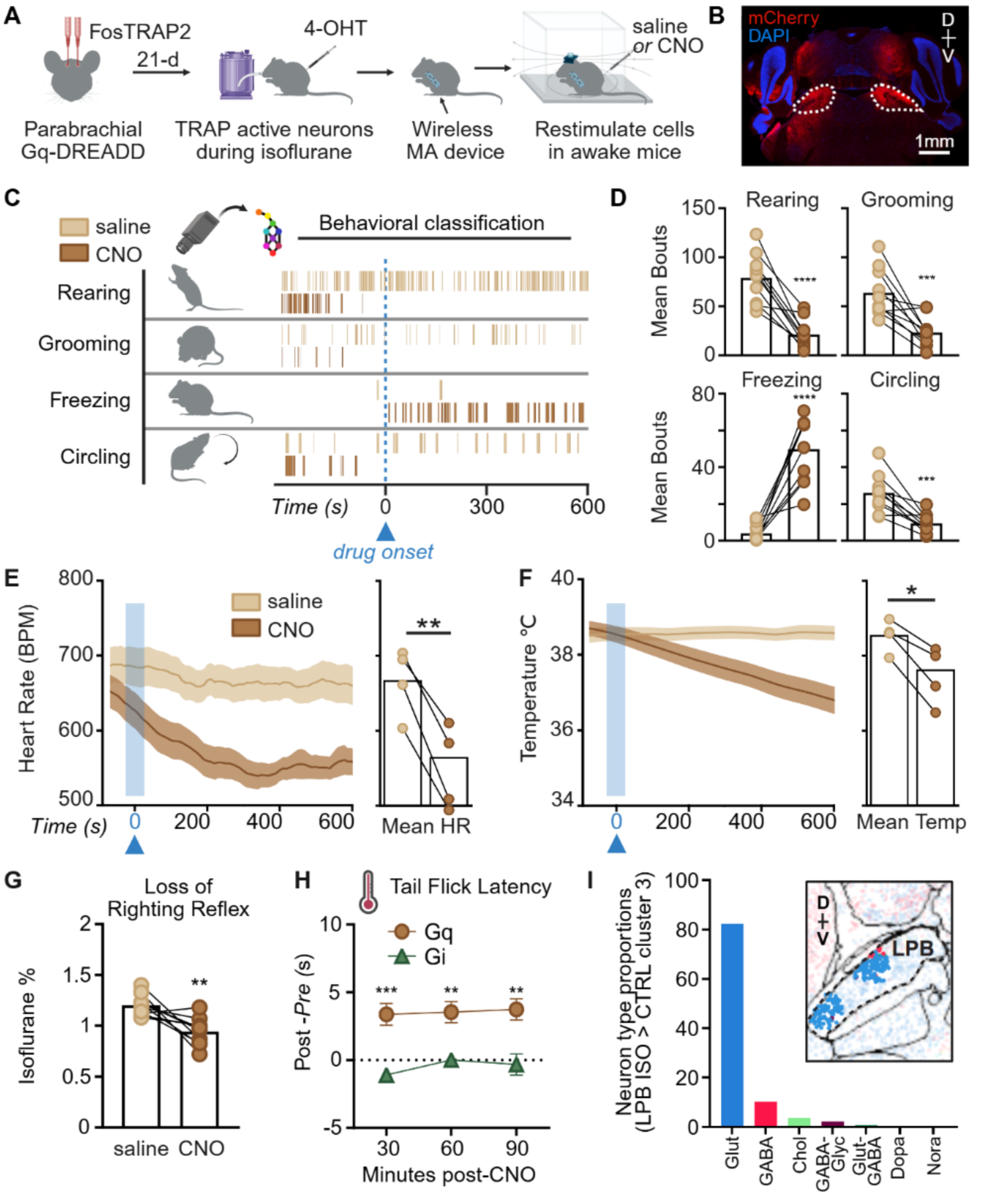
Chemogenetic activity-dependent capture of lateral parabrachial nucleus hub recapitulates components of altered consciousness. (**A**) Experimental timeline: FosTRAP2 transgenic mice received bilateral intra-parabrachial injection of AAV-PHP.eB Cre-dependent Gq-DREADD followed by activity-dependent capture (TRAP) during isoflurane exposure (1.2-1.3%) with 4-hydroxytamoxifen (4-OHT); (DREADD = designer receptor engineered to be activated by a designer drug). Mice were tested in the absence of anesthetic after either clozapine-n-oxide (CNO) or saline injection in a modified open field test with concurrent video-recording and peripheral physiology recording using wireless mechano-acoustic (MA) devices. (**B**) Representative image of mCherry-DREADD immunolabeling after viral expression in the LPB. Scale bar is 1 mm; also see **Fig S6**. (**C**) A representative raster plot from one animal’s classified behavior from the average of CNO and saline trials (also see **Video S5**). (**D**) Bouts of classified behavior for the average of all saline and CNO trials per animal (n = 10 mice, total of 29 CNO and 47 saline trials). (**E**) Left: Heart rate (beats per minute, BPM) decreases after CNO injection as compared to saline, shown time-aligned to approximated drug onset, right: mean heart rate (HR) decreases after CNO injection in Gq-DREADD mice (n = 4). (**F**) Left: Body temperature (Celsius) changes over duration of trial, time-aligned to drug onset, right: mean temperature decreases with CNO (n = 4). (**G**) Gq-DREADD mice display loss of righting reflex at a reduced isoflurane concentration after CNO injection (n = 10), though this does not occur at baseline. (**H**) Gq-DREADD mice (n = 10, brown) show increased latency to tail flick compared to baseline at every time point tested, while Gi-DREADD mice (n = 7, green) do not. (**I**) LPB cell-types activated by isoflurane analyzed using Fos maps overlaid with Allen Brain Cell Atlas predominantly consist of glutamatergic cells, as well as GABAergic and cholinergic subpopulations; see **Fig S7** for detailed analysis. 2-way repeated measures ANOVA with Sidak’s multiple comparisons: **p<0.01, ***p<0.001. Paired t-tests: *p<0.05, **p<0.01, ***p<0.001, ****p<0.0001. Error bars represent mean ± SEM. See **Table S5** for detailed statistics. See **Fig S6** for all Gi-DREADD data.

Similar to brain-wide ensemble manipulations, we observed a significant reduction in rearing and grooming behaviors after CNO induction, as well as prominently reduced locomotor movement (**Fig 7D**, **Fig S6**). We applied the same heuristic approach to quantify reduced movement bouts, finding that LPB ensemble stimulation induced significant freezing (**Video S5**). LPB ensemble re-activation did not induce circling bouts that were observed after brain-wide ensemble stimulation (**Fig 4G**), rather circling behaviors significantly decreased (**Fig 7D**). In peripheral physiological recordings, similar patterns of decreased heart rate and decreased temperature were induced (**Fig 7E-F**) with no significant change in respiratory rate after chemogenetic stimulation (**Fig S6H**). These behavioral and physiological effects were not observed in mice expressing inhibitory Gi-DREADDs in the LPB anesthesia-activated ensemble (**Fig S6E-G, S6I-K**). Overall, these data support the causal role of an LPB hub in generating unconsciousness.

CNO again potentiated loss of consciousness as defined by LORR, with LORR occurring at significantly lower ISO concentrations compared to the animal’s saline trials in Gq-DREADD mice (**Fig 7G**). However, LORR was not generated by stimulating the LPB neural ensemble in the absence of ISO. No significant potentiation of LORR by ISO was observed in Gi-DREADD mice (**Fig S6C**). After CNO injection (5 mg/kg, i.p.), neither Gq-DREADD nor Gi-DREADD mice showed a change in latency to fall off the rotarod when compared to their respective saline trials, indicating no changes in coordinated movement (**Fig S6A, D**). Given the established contribution of LPB circuitry to pain regulation, we also assessed the effects of chemogenetic induction on analgesia using the tail flick test. We observed significantly increased latency to tail flick in Gq-DREADD-expressing, but not Gi-DREADD mice, indicating that LPB ensemble stimulation, but not inhibition, promotes thermal anti-nociception (**Fig 7H**). Taken together, these data show that stimulation of the ISO-activated LPB neural ensemble is sufficient to recapitulate dissociable components of altered consciousness observed after brain-wide ensemble manipulation, establishing a causal contribution of LPB neural activation within the broader anesthesia network.

The LPB is a heterogeneous brain region composed predominantly of glutamatergic neurons with diverse transcriptional profiles, as well as smaller GABAergic subpopulations^65^. To determine if the ISO-activated LPB ensemble was dominated by a specific marker-defined neuronal subpopulation, we integrated ISO Fos maps (**Fig 1D-G**) with Allen Brain Cell Atlas MERFISH spatial transcriptomic data and quantified imputed neurotransmitter, LPB clade, and neuropeptide marker expression within the LPB portion of cluster 3 (**Fig 7I**, **Fig S7**, and **Table S5**). Fos-cluster neurons mapped onto diverse LPB populations, including *Atoh1*-related and *Lmx1*-related glutamatergic clades and sparse GABAergic populations (**Fig 7I** and **Fig S7**). Neuropeptides and cell-type-specific markers showed diverse expression patterns across LPB cluster neurons (**Table S5**). For example, *Pdyn* and *Cck* were enriched in the *Meis2*+ Atoh1-related clade, which comprised 58.4% of LPB cluster neurons, whereas *Calca*, *Nts*, and *Tac1* were enriched in the *Lmx1b*+ *Lmx1*-related clade, which comprised 27.0% of LPB cluster neurons (**Fig S7**). Together, these data indicate that no single marker-defined LPB cell type exclusively comprises the ISO-activated neural ensemble. Nearly all mapped glutamatergic LPB projection targets^65^ also showed increased Fos after ISO exposure, consistent with broader downstream network recruitment (**Fig S7**). In conclusion, the LPB ensemble is heterogenous and not defined by a single cell-type or subregion that underlies unconsciousness.

## DISCUSSION

In this series of experiments, we characterize the brain-wide ISO anesthesia unconsciousness network at cellular resolution. We test the hypothesis that neural circuit substrates of unconsciousness exist that can be selectively manipulated using activity-dependent chemogenetic manipulation to modify consciousness. We conclude that altered consciousness is recapitulated by manipulating either the brain-wide neural ensemble or an LPB hub, establishing a distributed neural circuit framework for anesthesia unconsciousness.

Recent circuit dissection studies in rodents have focused on stimulating or inhibiting isolated brain nuclei to uncover the contributions of individual brain regions to unconsciousness. Many studies have focused on hypothalamic nuclei that we also identify as significant, such as the preoptic nucleus. While these studies provide insights into region-specific contributions, they have also produced paradoxical findings. For example, targeted stimulation of MEPO contributes to arousal from anesthesia despite it also having high activity during sleep and anesthesia maintenance^30^. Selective stimulation of either GABA or glutamate subpopulations in the preoptic nuclei is insufficient for generating unconsciousness^66^. One potential explanation for these differences is that cell populations necessary for unconsciousness are functionally rather than genetically distinct, as we found in LPB, and activity-dependent manipulations are necessary to disentangle their contributions.

In addition to LPB, our whole brain connectome also identifies the MEPO as a key community node contributing to the unconscious state. The preoptic nucleus of the hypothalamus and the LPB may both contribute to hypothermia and coordinate autonomic responses^62,67–70^. The central amygdala is another community node that was found previously to modulate pain responding under anesthesia^18^. Since significant data exists to support causal contributions of hypothalamic^15,16,30,66,71,72^ and amygdala nuclei^18,73,74^ to anesthesia-induced unconsciousness, we focused our region-targeted experiments on LPB.

Previous studies have shown that stimulation of the parabrachial nucleus accelerates arousal from an unconscious state^45–47^. However, these prior studies were performed without activity-dependent stimulation and may recruit populations in the region that promote arousal. Spatially aligning our Fos maps with the Allen Brain Cell Atlas shows that the ISO-activated LPB cluster encompasses diverse LPB cell-types, including glutamatergic cells expressing calcitonin gene-related peptide (CGRP), as well as GABAergic populations^65^. These LPB cell-types have distinct, bidirectional effects on freezing behavior and pain responding, where glutamatergic CGRP-positive cells increase freezing^75^ and induce nociplastic pain^76^, while GABA cells alleviate pain when stimulated^77^. Our behavioral findings indicate that recruitment of both LPB cell populations likely occurs after activity-dependent capture, with GABA stimulation promoting anti-nociception and glutamatergic stimulation promoting freezing behavior, as previously reported in the literature. This divergence reflects the complexity of behaviorally defining heterogeneous neural interconnectivity and underlines the importance of harnessing integrated, activity-dependent and systems-level investigations for delineating neural mechanisms of unconsciousness.

Taken in context with the known effects of general anesthesia on cortical layer 5 pyramidal cells^7,20,29,78^, the distributed, LPB-centered network described here may be upstream of cortical cellular-level effects and reflect different levels of a shared process. Whether reactivating the anesthesia unconsciousness ensemble, or the LPB hub specifically, alters cortical coupling is an open and testable question to link these circuit and cellular mechanisms.

Analysis of single unit firing patterns indicates that a recruitment of ISO-activated neurons occurs with chemogenetic stimulation, particularly in the hindbrain. Many neurons exhibit similar suppressed firing with CNO induction, but ISO-activated neural firing patterns are different. This suggests either ISO recruits additional neuronal or non-neuronal mechanisms to generate unconsciousness, or the ensemble captured during ISO maintenance diverges from neurons necessary for induction of unconsciousness^79^.

Surprisingly, we find that neural ensemble activation alone does not generate LORR, reflecting a limitation of activity-dependent chemogenetic stimulation, or suggesting non-neuronal or non-specific, wet-blanket mechanisms are necessary for this loss-of-consciousness behavior^27,57,80–82^. Activating brain-wide neural ensembles induces an altered consciousness that is reflected by behavioral sedation and increased slow wave oscillations but not captured by LORR as a behavioral proxy. The slow wave state produced is comparable to a non-REM sleep pattern^2^, further supporting a convergence of anesthesia and sleep neural circuitry. Similarly, LPB ensemble activation potentiates LORR, though it does not generate it without ISO administration. The LORR behavioral definition is not applicable to altered states of consciousness like sleep and psychedelic states^83,84^ suggesting chemogenetic re-activation of anesthesia neural circuitry more closely resembles other altered states of consciousness.

Thermal anti-nociception accompanying the chemogenetic state transition may be confounded by and reflect the altered state of consciousness. We previously showed that anti-nociception can be recruited at lower CNO doses than those necessary for immobility after brain-wide activity-dependent manipulations using a different transgenic approach^85^, mimicking drugs that have analgesic effects at sub-anesthetic doses. Thus, brain-wide activity-dependent capture also provides a platform for dissociating neural circuit components of analgesia and unconsciousness circuitry, if such dissociations exist. Developing a deeper understanding of unconsciousness neural circuitry could inform therapeutic approaches, such as focused ultrasound^86^ or brain stimulation approaches^87^, to modify pain, sleep, coma and disorders of consciousness.

### Limitations of the study

While the TRAP2 approach is highly specific (>95% of activity-labeled cells are Fos+), it is only moderately efficient (∼65% of Fos+ cells are activity-labeled)^58^. Similarly, following retro-orbital injection at viral titers like our own (2-3x10^11^), PHP.eB neuron infection efficiency is high in the cortex (∼70% of neurons are infected), but moderate in other brain areas like the striatum (∼50% of neurons)^41^. Though these transgenic and viral efficiencies are sufficient for inducing significant behavioral changes, they introduce technical limitations for analyzing endogenous brain-wide mechanisms, so we also use endogenous Fos mapping. Our behavioral readouts are also limited by the temporal expression of Fos for activity-dependent capture, and the efficiency of CNO or DCZ to induce neural depolarization via DREADD receptor activation. We previously used a TRAP2 chemogenetic approach that uses transgenic DREADD expression independent of viral efficiency and report less robust behavioral results of brain-wide activation^85^. To account for temporal limitations of Fos as a readout of anesthesia activation, we also use single unit neurophysiology.

## RESOURCE AVAILABILITY

### Lead contact

**\*** Requests for further information and resources should be directed to and will be fulfilled by the lead contact, Mitra Heshmati.

### Materials availability

This study did not generate new unique reagents.

### Data and code availability

- All original code for behavioral analysis is available at https://github.com/sgoldenlab/simba
- All original code for UNRAVEL analysis is available at https://zenodo.org/records/13655988.
- All code for SMARTTR analysis is available at https://mjin1812.github.io/SMARTTR
- Datasets are included in compiled supplemental tables.
- Any additional information required to reanalyze the data reported in this paper is available from the lead contact upon request.

## ACKNOWLEDGMENTS

This project was funded by the National Institute of General Medical Sciences R35GM146751 (to MH), a Foundation for Anesthesia Education and Research Mentored Research Training Grant (to MH), and a National Institute of Health Small Business Innovation Research Grant R44MH140096 (to SAG and MH). We also thank funding from the National Institute of Mental Health F31MH133295 (to JN), R01MH130591 (to BDH), the National Institute on Drug Abuse R37DA033396 (to MRB), P30DA048736 (to MRB), P50DA042012 (to BDH) and R01DA059374 (to SAG), the National Institute on Aging F30AG084312 (to MJ), the Washington Research Foundation (to ES and SAG), the University of Washington Anesthesiology & Perioperative Medicine Research Training grant T32GM086270 (to KNS). We thank Yizhe Zhang, Erica Sanchez, Andy Pelos, Rachael Hansen, Lukas MacMillen and Leah Hulvershorn for their technical support. We thank Dr. Richard Palmiter and Dr. Sekun Park for feedback on lateral parabrachial investigations and sharing Fos-2a-Cre transgenic mice. We thank Dr. Cameron Good for supporting the mechano-acoustic device at NeuroLux, Inc. We also thank Dr. Nicholas Steinmetz and Dr. Zhiwen Ye for feedback on the high-density silicon probe neurophysiology studies.

## AUTHOR CONTRIBUTIONS

MH and SAG conceptualized and directed all aspects of the study. MH, SAG, KNS, MHCN, DRR and MJ wrote the first draft of the manuscript. SAG, MH and MRB designed whole brain studies. KNS, MHCN, HL, DRR, JN, ADM, ACB, IC, MH and SAG collected, processed and/or imaged brain tissue. DRR, ADM, ES, IC, KI, GS, BDH, SAG and MH performed whole brain analysis. MJ and CAD performed network analysis. JN and ES prepared viruses and performed retroorbital virus injections. MHCN, HL, CN, ACB and MH performed activity-dependent chemogenetic behavioral experiments. MHCN, HL, NR and AL collected and analyzed mechano-acoustic recordings. SRON, HL, MHCN, CCS, NR, EA, JZ, and AL analyzed behavioral video recordings. KNS performed and analyzed high density silicon probe experiments. AB, GG and HOdlI performed electrocorticography recordings and analysis. All authors edited and approved the manuscript.

## DECLARATION OF GENERATIVE AI AND AI-ASSISTED TECHNOLOGIES

During the preparation of this work, the authors used AI-assisted technologies [Claude and ChatGPT] for proofreading and grammatical editing. After using these tools, the authors reviewed and edited the content and take full responsibility for the content of the publication.

## SUPPLEMENTAL TABLES

**Supplemental Table S1:** Brain-wide Fos detailed statistics and supplemental data (Fig 1)

**Supplemental Table S2:** Network analysis permutation results, unique regions analyzed, and detailed network community node centrality statistics (Fig 2)

**Supplemental Table S3:** Brain-wide TRAP Supplemental Data: Mechano-acoustic internal state physiology during isoflurane (Fig 3), extended behavior and electrocorticography results (Fig 4), behavioral classifier statistics (Fig 4), Brain-wide TRAP ClearMap data (Fig 5)

**Supplemental Table S4:** Neuropixels high density single unit recordings detailed statistics and anatomical distributions (Fig 6)

**Supplemental Table S5**: Parabrachial TRAP detailed behavior, Parabrachial MERFISH data (Fig 7)

## SUPPLEMENTAL VIDEOS

**Video S1**. Anesthesia-activated Fos clusters and their spatial relationship to Neuropixels recordings. Rotating 3D brain models show UNRAVEL clusters representing increases or decreases in Fos+ cell density after isoflurane. Subsequent views overlay Neuropixels units and recording channels, including units within the LPB Fos cluster.

**Video S2**. Anatomical distribution of community network analysis video. Rotating 3D brains depict key community nodes from functional network analysis.

**Video S3**. Isoflurane induction with dynamic mechanoacoustic physiology recording and automated behavioral classification (see **Fig 3**).

**Video S4**. Brain-wide isoflurane-activated ensemble chemogenetic induction with mechanoacoustic physiology recording and automated behavioral classification. Representative video clips at 4X speed from one Gq-DREADD mouse and one Gi-DREADD mouse’s saline and CNO trials after brain-wide TRAP under isoflurane (see **Fig 4**).

**Video S5**. Lateral parabrachial ensemble chemogenetic induction with dynamic mechanoacoustic physiology recording and automated behavioral classification. Representative video clips at 4X speed from one Gq-DREADD mouse and one Gi-DREADD mouse’s saline and CNO trials after LPB TRAP (see **Fig 7**).

## STAR*METHODS

### KEY RESOURCES TABLE

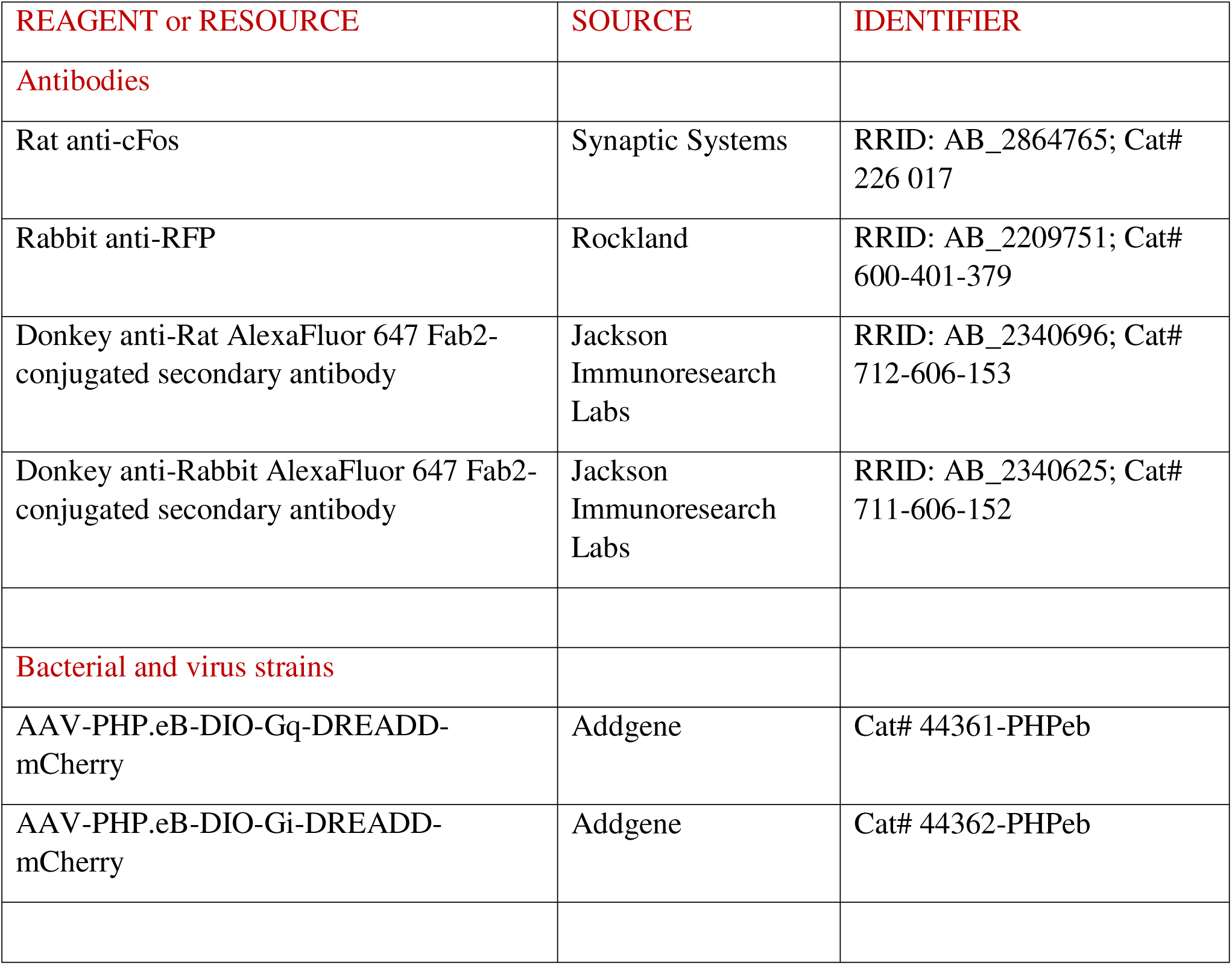
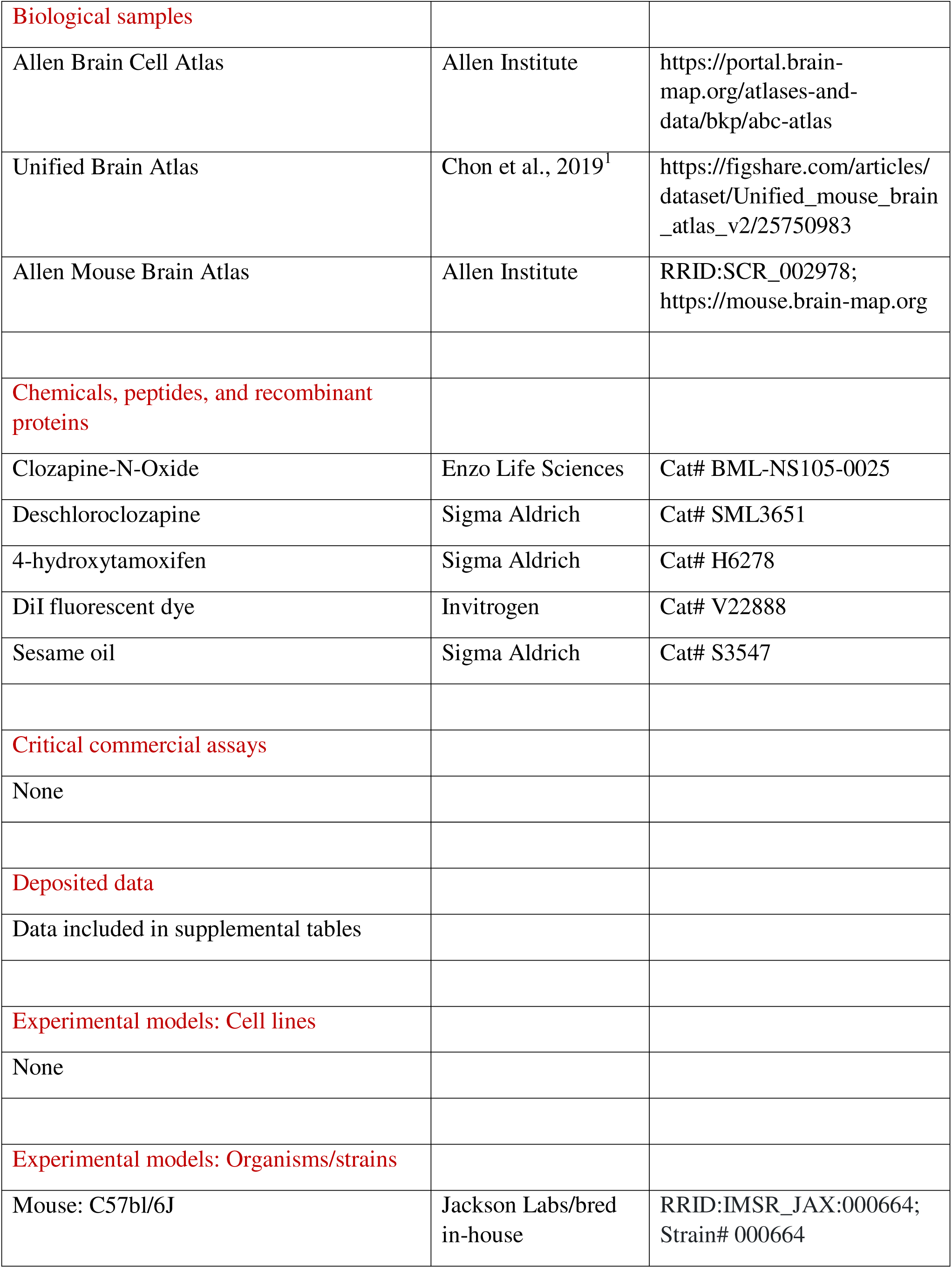
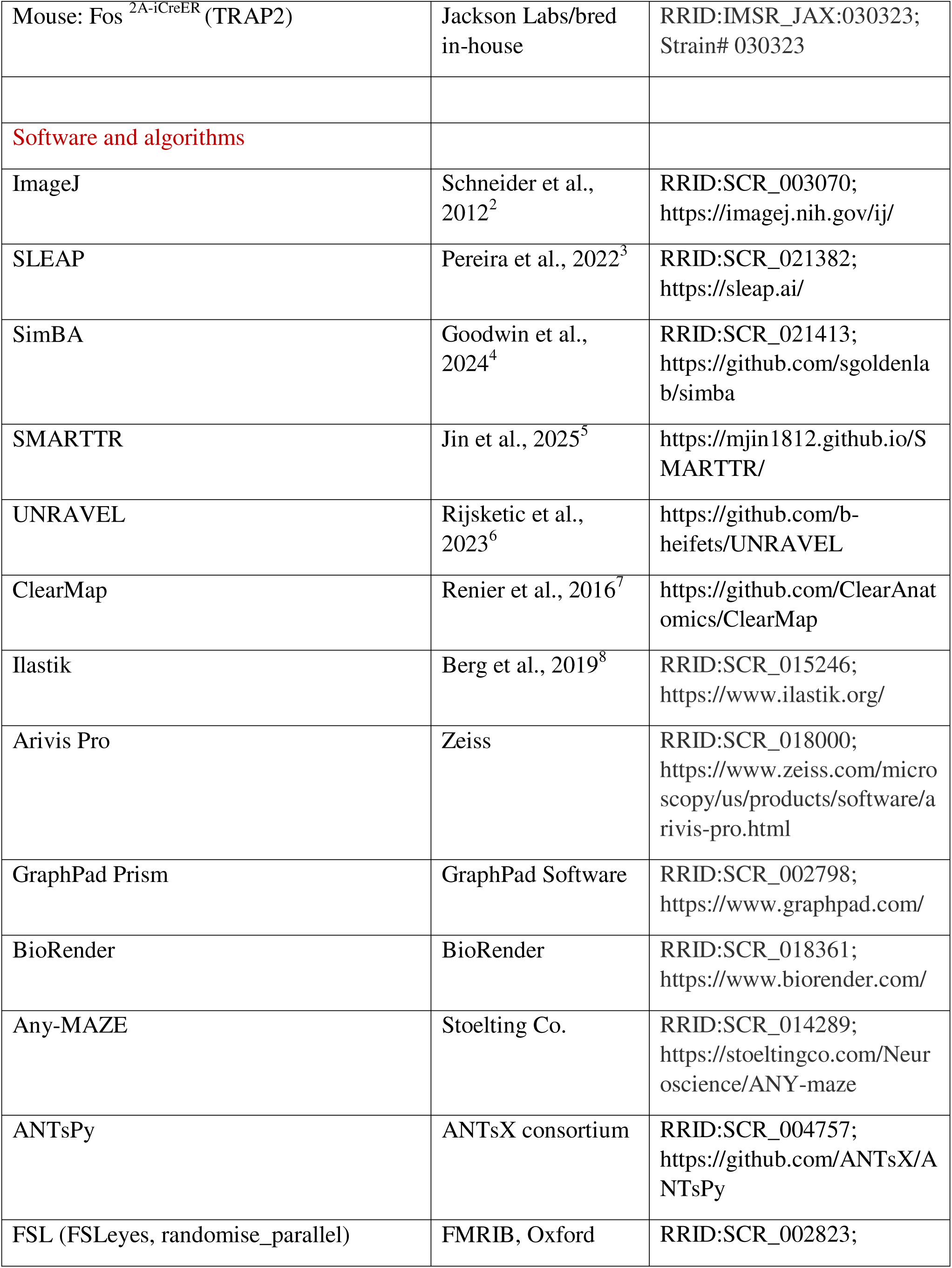
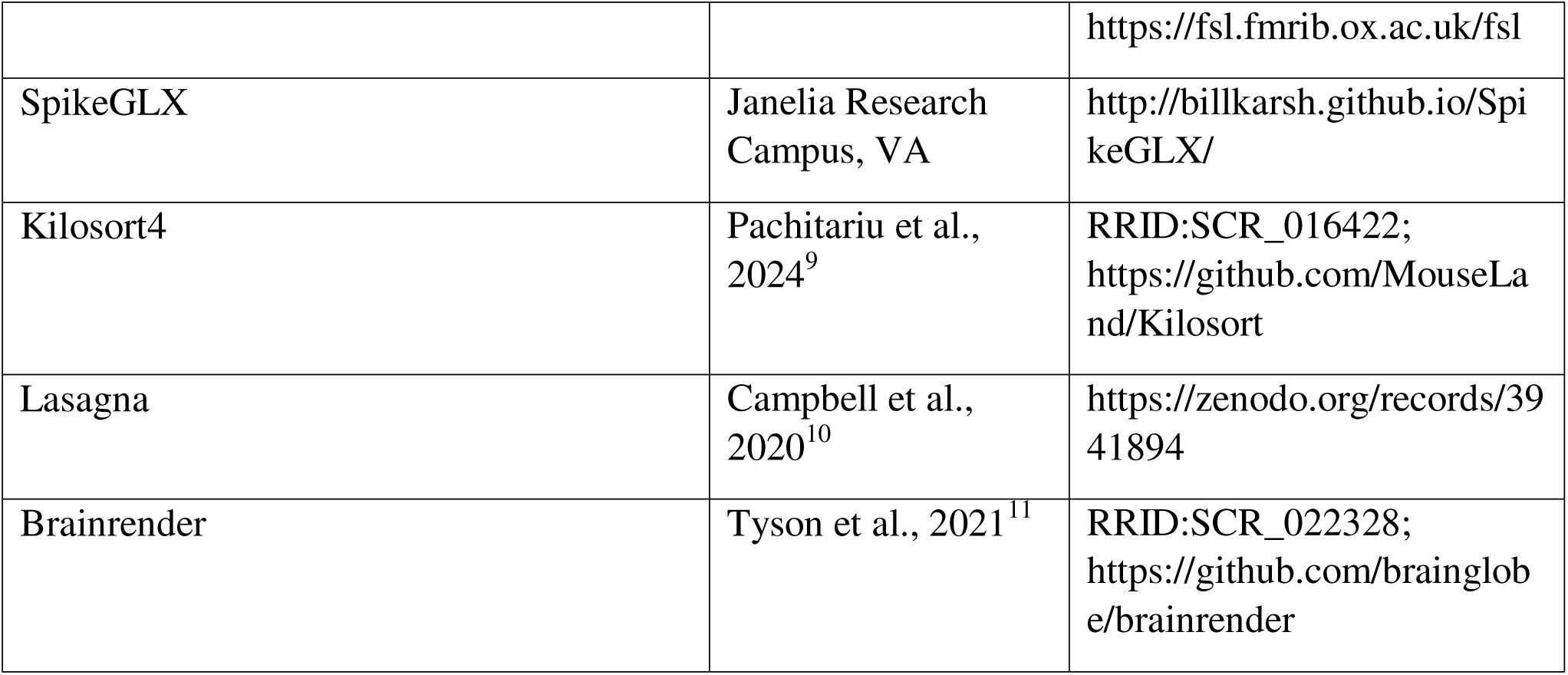

#### METHOD DETAILS

##### Mice

All experiments were approved by the National Institutes of Health and the University of Washington Institutional Animal Care and Use Committee. 3-to 6-month-old male and female C57bl/6J or homozygous Fos ^2A-iCreER^ (FosTRAP2) transgenic mice were used in all studies. Animals were group-housed in a 12-hour reverse light/dark cycle facility with *ad libitum* access to food and water. Animals used in electrocorticography or Neuropixels studies were singly housed during the recording experiments.

##### Intact brain tissue collection after isoflurane exposure

Mice were singly housed in the home cage for 3 days prior to tissue collection for intact whole brain Fos immunolabeling. Habituation to the testing room was achieved by placing mice in a temperature controlled red-light room for at least 2-3 hours a day for 3 days prior to tissue collection. On the test day, mice in control condition remained singly housed in their home cage inside the testing room where isoflurane was delivered. Mice in the isoflurane condition were individually induced with 2% isoflurane and after loss of toe pinch reflex was confirmed, each mouse was maintained under general anesthesia for 180 minutes with 1.2%-1.3% isoflurane in oxygen delivered via nose cone. The concentration of isoflurane delivered at each nose cone was confirmed to be within range using a gas analyzer prior to placing the mouse in the nose cone. Mice were actively warmed on a heating pad and continuously monitored by the experimenter. After 180 minutes, all mice were more deeply anesthetized with isoflurane and transcardially perfused with phosphate-buffered saline (PBS) followed by 10% formalin. Brains were dissected and post-fixed in formalin for 24 hours at 4°C prior to intact whole brain clearing and immunolabeling.

##### Intact whole brain clearing and immunolabeling

A previously published, modified version of the iDisco+ protocol was used to immunolabel and clear intact brain samples^7,12^. The following antibodies were used: Rat anti-cFos (1:2000; Synaptic Systems) for Fos immunolabeling or Rabbit anti-RFP (1:2000; Rockland) for mCherry DREADD immunolabeling, followed by Donkey anti-Rat or Donkey anti-Rabbit AlexaFluor 647 Fab2-conjugated secondary antibody (1:500; Jackson ImmunoResearch).

##### Intact whole brain imaging

Cleared, intact whole brains were imaged with a uniform axial resolution light sheet microscope (SmartSPIM, LifeCanvas Technologies) using either the 1.6x (5 µm voxels) or 3.6x (1.8 µm voxel) objectives for Fos samples, and 3.6x for TRAP samples. Samples were mounted horizontally using a custom sample holder immersed in dibenzyl ether (DBE). Images were acquired in two channels (488 nm: autofluorescence; 639 nm: mCherry/AlexaFluor-647) and stitched using the SmartSPIM acquisition software. TRAP+ images were down-sampled (0.5x) prior to image processing with isotropic 3.6 µm voxels. 3-dimensional images were visualized using Arivis Pro software (Zeiss).

##### Retroorbital virus injection

FosTRAP2 mice were briefly anesthetized with isoflurane (5-10-min) and administered adeno-associated virus (AAV) by retroorbital injection to one eye. AAV-PHP.eB-hSyn-DIO-hM3D(Gq)-mCherry (44361-PHPeB, Addgene) was used for activating DREADD and AAV-PHP.eB-hSyn-DIO-hM4D(Gi)-mCherry (44362-PHPeB, Addgene) was used for inhibitory DREADD. Virus was diluted in total 100µL sterile saline to ∼ 2-3x10^11^ GC/mL based on prior studies^13^.

##### Lateral parabrachial nucleus viral injection

FosTRAP2 mice were anesthetized with isoflurane (1.3%-3%) and head-fixed for survival surgery. They also received single injections of lidocaine-bupivacaine and ketoprofen for peri-operative analgesia. Stereotactic coordinates were standardized relative to Bregma and Lambda distance, and a Hamilton syringe was used to inject 0.4ml of virus at a rate of 0.2ml/min. AAV-PHP.eB-hSyn-DIO-hM3D(Gq)-mCherry (44361-PHPeB, Addgene) or AAV-PHP.eB-hSyn-DIO-hM4D(Gi)-mCherry (44362-PHPeB, Addgene) was injected bilaterally into the PBN (AP: -4.90mm, ML: +/-1.35mm, DV: -3.40mm). Mice recovered from surgery for at least two weeks prior to TRAP, followed by at least 1-week post-TRAP recovery prior to behavioral testing.

##### Activity-dependent labeling (TRAP)

3-4 weeks following retroorbital or intracranial virus injection, FosTRAP2 mice were induced with 2% isoflurane and then maintained unresponsive at 1.2-1.3% for a total time of 180 min. Mice were actively warmed on a heating pad and continuously monitored by an experimenter. A single 50 mg/kg intraperitoneal (i.p.) injection of 4-hydroxytamoxifen (4-OHT, Sigma Aldrich), was administered at the 90-min halfway point during the isoflurane exposure to induce *Cre*-recombination and activity-dependent labeling^14^. Non-TRAP control mice also received retroorbital virus injections and isoflurane exposure but then sesame oil vehicle (Sigma Aldrich) in the equivalent volume of 4-OHT (∼0.2mL) was given i.p. at the 90-min halfway point.

##### Clozapine-n-oxide administration

Clozapine-n-oxide (CNO, Enzo Life Sciences) was solubilized in water and stored as 5mg/mL aliquots. For testing, the aliquot was diluted in sterile saline to an end concentration of 0.5 mg/mL for 5 mg/kg behavioral studies. Mice received i.p. injections according to their body weight ranging from 0.2-0.3 mL for a 20-30g mouse. The equivalent volume i.p. sterile saline was delivered for saline trials.

##### Deschloroclozapine administration

Deschloroclozapine (DCZ, Millipore Sigma) was solubilized in DMSO and stored as 2mg/mL aliquots. For testing, the aliquot was diluted in sterile saline to an end concentration of 0.01mg/mL for 0.1mg/kg behavioral studies. Mice received i.p. injections of DCZ according to their body weight.

##### ClearMap brain-wide TRAP+ analysis

We processed 7 intact formalin-fixed brain samples from FosTRAP2 transgenic mice infected with AAV-PHP.eB-DIO-Gq-DREADD-mCherry virus (see *Retroorbital virus injection* and *Activity-dependent labeling* procedures) using a modified version of the ClearMap pipeline as previously published^12,15^. We trained a pixel classifier in Ilastik^8^ to segment and quantify single cells based on somatic signal classification. This was achieved by selecting 21 image tiles (200 x 200 x 220 gm) from three separate samples, cropped from dorsal (2), central (3), and ventral (2) positions of the brain. To improve segmentation accuracy, an additional three tiles were used to correct for artifacts in ventricular and boundary regions. The fully trained classifier was then applied to all samples, generating segmented cell counts per brain region and voxelized heatmaps. We computed TRAP+ cell counts within two different levels of hierarchy within the Unified Brain Atlas^1^: 1) 11 major anatomical divisions (**Fig 5**) and 914 minor subregions within those high-level anatomical divisions that no longer split into further daughter regions. We provide the raw TRAP+ cell counts, cell density, and statistical results for all analyses in **Table S3**.

##### ClearMap brain-wide Fos analysis

We processed 16 intact brain samples (n = 7 controls and n = 9 isoflurane) using a modified version of the ClearMap pipeline as previously published^7,12^. We computed Fos-positive activated cell counts within two different levels of hierarchy within the Unified Brain Atlas ^1^: 1) 11 major anatomical divisions and 914 minor subregions within those high-level anatomical divisions that no longer split into further daughter regions (**Fig 1A-C**; **Fig S1**). To ensure accuracy within the hierarchical relationships, we ensured there were no overlapping spatial footprints and no double counting of any regions, following each subregion branch until the end or “stop-level.” We transformed the raw Fos cell counts to z-scores and normalized the data to account for regional volume differences. We used the home cage control group (control) to normalize the z-scores for each region of interest using the formula z = (x-g)/o, where x represents the Fos count of the treatment group, g represents the mean Fos count of the control group, and o represents the SD of the control group. We then converted the z-score into -log10 for visualization. For the high-level analysis, we used an unpaired t-test, *p < 0.05 (**Fig S1G**). For the stop-level analysis, we used a t-test followed by the false discovery rate (FDR) multiple comparison correction with an alpha significance level of 0.01 (q < 0.01), two-tailed (**Fig S1H**). We provide the raw Fos counts, z-scored data and statistical results for all analyses in **Table S1**.

##### UNRAVEL Fos analysis

We also analyzed whole brain Fos immunolabeling using UNRAVEL^6^ (**Fig 1D-G**, **Fig S2**, **Fig S7**; **Video S1**) (UN-biased high-Resolution Analysis and Validation of Ensembles using Light sheet images), with refactored Python code (documentation: b-heifets.github.io/UNRAVEL/index.html). Autofluorescence images were resampled to 50 gm resolution, and tissue was masked using Ilastik (v1.4.0; pixel classification; all features) for N4 bias field correction (ANTsPy; 0.4.2). The resampled images were padded (15% voxels on all sides) and smoothed with a Gaussian filter (sigma = 0.4) before being registered with an average template from iDisco and light-sheet microscopy (LSFM), which was aligned with the Allen Mouse Brain Common Coordinate Framework (CCFv3)^16^. Registration was performed using ANTsPy: ants.affine_initializer(fixed_image=autofluorescence_image, moving_image=template, search_factor=1, radian_fraction=1, local_search_iterations=500), followed by ants.registration(fixed=autofluorescence_image, moving=template_aligned_with_tissue, grad_step=0.1, syn_metric=’CC’, syn_sampling=2, reg_iterations=(100, 70, 50, 20)). Registration accuracy was visually confirmed using FSLeyes (v0.30.1; FMRIB), with the atlas warped to each fixed registration input image. Fos immunofluorescence was enhanced through 3x3x3 spatial averaging and rolling ball background subtraction (pixel radius = 4). The resulting images were warped to 25 gm atlas space using transforms from registration. Each hemisphere was z-scored using a warped tissue mask and a hemispheric mask ((image -mean intensity in the brain)/standard deviation of intensity in the brain). The images were smoothed with a 100 gm kernel and mirrored to average the left and right sides. A unilateral atlas mask, excluding ventricles, fiber tracts, and undefined regions, spatially restricted the voxel-wise analyses.

Voxel-wise comparisons between the isoflurane and control groups (t-test design) were performed using non-parametric permutation testing (18,000 permutations) with the general linear model (randomise_parallel; FMRIB; v6.0.2). P-value maps were corrected for multiple comparisons using the false discovery rate (FDR) method, applied across a series of q-values to identify clusters of significant voxels. Data in **Fig 1D-G** are from the most stringent q-value thresholds (q < 0.005, p < 0.000056 for Control < Isoflurane; q < 0.05, p < 0.0012 for Control > Isoflurane). Clusters smaller than 100 voxels were excluded. Clusters were warped back to full-resolution tissue space for cell density measurements (Fos+ cells per cluster volume). Fos+ cells were segmented using Ilastik. 3D counting was performed using connected components-3d (v3.16.0; connectivity = 6). Clusters were considered valid if unpaired t-tests confirmed that differences in Fos labeling reflected changes in Fos+ cell densities. Information on valid clusters, including regional composition and Fos+ cell densities across FDR q-values, is provided in **Table S2**. Representative images are provided in **Fig S2**. Statistical maps and other related data are available upon request. Scripts from UNRAVEL (github.com/b-heifets/UNRAVEL) are available at zenodo.org/records/13655988.

##### UNRAVEL LPB cell types and marker expression analysis

To characterize the cellular composition of the isoflurane-activated lateral parabrachial nucleus (LPB) ensemble, the Control < Isoflurane UNRAVEL cluster 3 mask from the whole-brain Fos analysis was transformed into Allen Brain Cell Atlas (ABCA) MERFISH-CCF space using UNRAVEL. The LPB portion of cluster 3 was defined by intersecting the transformed cluster mask with the PB label in MERFISH-CCF space. Unified Mouse Brain Atlas v2 labels were also transformed into MERFISH-CCF space to visualize the relationship between cluster 3 and LPB subregions, including external LPB (LPBE) and dorsal LPB (LPBD). Neurons located within the LPB portion of cluster 3 were retained for downstream analysis and summarized using ABCA neuronal taxonomy annotations.

Cell-type composition was summarized across the ABCA hierarchy, including neurotransmitter class, class, subclass, supertype, and cluster. For visualization in **Fig S7**, sunburst plots were collapsed to three ABCA levels for readability, whereas **Table S5** provides summaries across all five ABCA annotation levels. In sunburst plots, wedge size represents the percentage of LPB cluster 3 neurons assigned to each ABCA category. For marker-expression plots, wedge color represents mean imputed expression, shown as logD(CPM + 1).

Using imputed MERFISH data from Yao et al. (2023)^17^ for each retained neuron, marker gene expression was quantified for all present cell types. Marker selection was guided by the LPB molecular and projection framework described by Pauli et al. (2022)^18^. For each marker gene and ABCA annotation level, we calculated mean imputed expression and the percentage of marker-positive neurons. Marker-positive neurons were defined as cells with imputed expression >3 logD(CPM + 1). *Atohl* -related glutamatergic neurons were operationally defined as *Slc17a6*+/*Meis2*+/*Lmx1b*-, *Lmx1*-related glutamatergic neurons as *Slc17a6*+/*Meis2*-/*Lmx1b*+, inhibitory neurons as GABAergic neurons, and remaining neurons as other.

For the **Table S5** overview, each marker was summarized by identifying, at each ABCA annotation level, the most prevalent category within LPB cluster 3 among categories with mean imputed expression >3 logD(CPM + 1). Percent values in the ‘Marker overview’ tab indicate the percentage of LPB cluster 3 neurons assigned to the listed ABCA category. Categories with no mean expression above threshold at a given ABCA level were denoted as not detected (n.d.). Proportions of all present cells can be found in the ‘LPB cluster 3 neurons’ tab. Look up tables in the ‘LUT’s’ tab can apply sunburst plot coloring in Flourish. Other tabs characterize marker gene expression across cell types.

Major LPB projection pathways and target regions were adapted from Pauli et al. (2022)^18^. To relate LPB projection targets to isoflurane-induced Fos activity, mapped target regions were annotated using the Control < Isoflurane UNRAVEL analysis, noting Fos upregulation in LPB target regions at FDR q < 0.005 and 0.005 ≤ q < 0.05.

##### Correlational, Permutation, and Network Analysis

All associated network analyses and visualizations were conducted with the SMARTTR^5^ package, igraph, tidygraph package, and custom written functions (**Fig 2**). Pearson correlations between normalized Fos^+^ activity per region were calculated and asymptotic p-values for each correlation were determined using a one-sample t-test. Significantly correlated regions are visualized in heatmaps after FDR (Benjamini-Hochberg) correction using a significance threshold of q < 0.05. A permutation analysis was conducted to identify the functional connections that differed most between experimental groups. Correlation differences were calculated by subtracting correlations in the Control group from respective correlations in the Isoflurane group (r_Isoflurane_ – r_Control_ ). Each correlation difference was used as a test statistic against a null distribution produced by shuffling the labels of Control and Isoflurane mice 10,000 times, with the correlation difference recomputed for each shuffle. Each test statistic was compared to its individual null distributions to determine the p-value. Significance correlation differences for all analyses were thresholded using an alpha value of 0.001 and visualized as a constructed network. Edge (line) thickness in the permuted network represents the magnitude of correlation difference, with regions represented as nodes. An agglomerative fast greedy modularity optimization algorithm^19^ was applied using tidygraph for community detection.

Thresholding is normally applied to assess the topological features of the most salient connections, although there is little consensus on the best approach, (i.e., thresholding by connection strength or connection proportion) or choice of threshold^20^. Thus, we constructed networks for Control and Isoflurane across a range of alpha thresholds (FDR uncorrected) from 0.0001 to 1, and across a range of edge proportion thresholds, from 0.001 to 0.50.

In both the permuted and individual functional networks, several topology metrics per node, such as degree, clustering coefficient, efficiency, and betweenness centrality^21^ are automatically calculated using the create_networks() or create_permutation_network() functions in SMARTTR. Summary statistics are calculated using the summarise_networks() function, which averages nodal metrics to calculate mean degree, mean efficiency, mean clustering coefficient, and mean betweenness for a given network. To assess for “small-world” properties^22^ of empirical Control and Isoflurane networks (edge proportion of 3.15%), we compared the clustering coefficient (*C*) and characteristic path length (*L*) with the corresponding values (*C^rand^, L^rand^*) averaged from 100 random null networks. Null networks are generated by applying the degree sequence-preserving maslov-sneppen^23^ rewiring algorithm with the rewire_network() function, and averaged metrics were calculated with summarize_null_networks(). Number of edge rewirings per random network are equivalent to number of edges x 100. The small-world index (s), a ratio summarizing the propensity for high clustering while maintaining efficiency in a network, is defined here as [Equation]. Values above 1 are considered indicative of small-world properties. All analysis code is available on (https://mjin1812.github.io/SMARTTR). See also **Tables S3-S5** for detailed statistics.

##### Loss of righting reflex

Latency to loss of righting reflex was assessed in a chamber equipped for isoflurane delivery and sampling of isoflurane concentration with a gas analyzer (**Fig 4**, **Fig 7**, **Fig S4** and **Fig S6**). Mice were individually injected 5mg/kg CNO, 10mg/kg CNO, 0.1mg/kg DCZ, or saline with 10 mg/kg CNO or saline in alternating trials at least one day apart and placed in the chamber 15 minutes after injection. Isoflurane was administered stepwise from 0.5 to 2% on the vaporizer in 0.5 increments and loss of righting reflex was assessed at each stepwise increment with the exact sampled concentration recorded.

##### Warm water tail withdrawal test

Mice were tested for the latency to remove their tail from a warm water bath (52.5°C). Mice are restrained within a paper towel, and 1/3 to ½ of the tail is immersed, and the time to remove the tail (tail flick latency) is recorded^24–27^. This is done at baseline after saline, and then in 30-min intervals after CNO injection. See **Table S3** and **Table S5** for detailed statistics.

##### Hot plate test

Mice are placed on a 53°C plate for 30 seconds, and behaviors are video recorded^28^. This occurs at baseline and after CNO at 2-time intervals (30-min and 90-min post-CNO) interleaved between tail withdrawal tests. Latency to jump, groom paws, and number of occurrences for these behaviors are quantified from videos by two independent researchers. See **Table S3** for statistics.

##### Rotarod test

Coordination post CNO injection was tested for each mouse on a rotarod^29^ after modified open field testing. Mice were habituated for two days, 10-min per day, at 4 rotations per minute (rpm) before testing. On test day, mice received a saline i.p. injection in the morning and were placed on the rotarod at 4 rpm. Latency to fall was recorded, unless animals stayed on for the duration of the experiment (10 min). Mice were recorded again in the afternoon with a 5mg/kg i.p. CNO injection. See **Table S3** and **Table S5** for statistics.

##### Annotation

Behavior was annotated in 20-min video clips obtained from the modified open field test. For fitting the Straub tail classifier, we used a sampled annotated dataset where the behavior was present in 24244 frames (808s) and behavior was absent in 627312 frames (20910s). For fitting the grooming classifier, we used a sampled annotated dataset where the behavior was present in 47005 frames (1566s) and behavior absent in 94010 frames (3133s). For fitting the rearing classifier, we used a sampled annotated dataset where the behavior was present in 31596 frames (1053s) and behavior absent in 315960 frames (10530s).

##### Featurization

We featurized video and pose-estimation data using image, movement, geometry, circular-and frequentist statistical methods (see below). For acceptable runtimes, we used GPU accelerated and multiprocessing methods available through the SimBA API^4^ . Features were computed in metric space after calculating the pixel-to-metric conversion factors in the SimBA graphical interface. Classifications were smoothened to remove any detected bouts less than 200ms in length. Computations are primarily dependent on numba^30^ and shapely^31^ libraries and the featurization classes are available on the SimBA GitHub repository.

##### Grooming and rearing behavioral classifiers

We used the same feature set to create classifiers for rearing and grooming. This feature set included movement and acceleration measures of individual body-parts as well as paired body-part movement Spearman correlations in sliding time windows (0.25-2s). We also computed distribution descriptive statistics (mean, variance, sum, z-scores, skew, median absolute deviations, mean absolute change, root mean square) of the animal hull geometry and animal hull sub-geometries (animal hull width, length, and area, as well as head area, posterior hull area, anterior hull area, left lateral hull area, right lateral hull area) in sliding windows of 0.25-2s. We computed Spearman correlations between the hull sub-geometry areas in the same sized sliding time-windows. Using circular statistics, we calculated descriptive statistics of animal directions including animal circular range, animal circular standard deviation, and directional vector lengths in sliding windows of 0.25-2s.

##### Straub tail classifier

For accurate Straub tail segmentation, we first performed video background subtraction and centered and egocentrically aligned the mice in the videos. Next, we defined a buffered (1.75cm in each direction) polygonal geometry using the mouse tail key-points (tail-base, tail center, tail-tip) and sliced the tail geometry from each frame. Using the tail geometry coordinates, we computed the tail geometry mean area, standard deviation, mean absolute deviation in sliding time-windows of 1-2s. For each frame, we compared the tail geometry to the tail geometry 0.5-2s seconds prior using Hausdorff distances. For the sliced images of the tails, we compared the current tail image versus the tail image 0.5-2s prior using mean squared error of the raw RGB pixel values. We also computed descriptive statistics (average, variance, sum, mean absolute change) of the tail tip and tail center movements, and the tail length in sliding windows of 1-2s.

##### Feature importance

Model Gini feature importance scores showed the following key feature determinants for each behavior: (1) *Grooming* was primarily influenced by features describing the distribution of the animal’s head area, body length, and snout movement within sliding temporal windows. (2) *Rearing* was primarily driven by features capturing the distribution of the distance between the animal’s ears, head area, and body width in sliding temporal windows. (3) *Straub tail* was mainly determined by features describing the distribution of the tail-tip movements, tail length and area, overall body area and right-side hull area, as well as correlations between the movements of the tail tip and tail base in sliding temporal windows. See **Table S2** for detailed feature importance.

##### Performance

Out-of-sample frame wise performance showed strong performance for grooming (precision: 0.92, recall: 0.99, f1: 0.96), rearing, (precision: 0.98, recall: 0.96, f1: 0.97) and Straub tail (precision: 0.99, recall: 0.96, f1: 0.98). See **Table S2** for model metrics.

##### Heuristic models

Freezing and circling were detected using heuristic rules as previously described^32,33^. Circling was scored as present when the directional circular range of the animal was above 320 degrees, and the animal movement (computed from the animal centroid key-point) was above 6cm in the preceding 10s time-window. Freezing was scored as present in frames where the velocity (computed from the mean movement of the nape, snout, and tail-base body-parts) fell below 5 mm/s in the preceding 3s. Although the nape body-part was not pose-estimated, this body-part was approximated to be halfway between the two ears.

##### Wireless mechano-acoustic device implant

Mice were subdermally implanted with wireless mechano-acoustic devices (MA device: nVital, NeuroLux, Inc.) on the ventral surface while under isoflurane anesthesia for a duration of less than 30 min. They also received single subcutaneous injections of lidocaine-bupivacaine and ketoprofen for peri-operative analgesia. They were then group housed following surgery. The devices contain an accelerometer adhered to the animal’s sternum with GLUture, mechanically coupled to cardiac, respiratory, and behavioral motion, as well as a temperature sensor. Mechano-acoustic and temperature signals stream via Bluetooth at a rate of 800Hz and 1Hz, respectively. Wireless power transfer by resonant magnetic inductive coupling allows the devices to wirelessly charge and transmit data. See Ouyang et al. (2024)^34^ for surgical procedure and device details.

##### MA physiological data processing & analysis

For heart rate, first heart sounds were found by subtracting the behavioral motion component of the z acceleration signal, as estimated by a Savitzky-Golay filter (8th order, 21-point window), from the raw z acceleration. The lowpass filtered (15Hz cutoff) Shannon energy envelope of the heart sound signal revealed individual heart beats. Average beat-to-beat intervals per second yielded heart beats per minute. The respiratory signal was found by down-sampling z-axis acceleration to 50Hz, and band-pass filtering (2 -5 Hz cutoff) revealed individual respirations. Average beat-to-beat intervals per second yielded respirations per minute. Physical activity was calculated by finding the average variance of each acceleration axis (normalized to system noise variance) in 1 second rolling windows and taking the square root to quantify acceleration variability in arbitrary units. bpm: beats-per-minute; rpm: respirations-per-minute; a.u.: arbitrary units. See Ouyang et al. (2024)^34^ for further details. Statistical comparisons were performed on raw (unsmoothed) physiological metrics averaged across a predefined 5–15 min post-injection window. Normality of paired differences per subject was assessed using the Shapiro-Wilk test. Differences between chemogenetic modulation conditions (CNO vs saline) were tested using one-tailed t-tests. Data processing was performed using custom Python scripts (scipy.stats), and group means and paired t-tests were conducted in GraphPad Prism (see **Table S3** for detailed statistical outputs).

##### Modified open field test and MA device data collection for isoflurane induction (**Fig 3**)

A 15cm x 15cm smaller, modified open field box was adapted with an isoflurane in oxygen delivery inlet, gas analyzer sampling line, and scavenge. The sides of the box were wrapped in 14G copper wire attached to tuner and power distribution control (PDC) boxes to charge the implanted devices and allow data streaming (NeuroLux, Inc). 7 animals (2 trials each) were recorded under isoflurane exposure. Trials consisted of 5 minutes of recording before isoflurane onset, 20 minutes following loss of reflex to a loud clap, and 20 minutes after isoflurane delivery ended.

##### Modified open field test and MA device data collection for chemogenetic studies (**Fig 4** and **Fig 7**)

For 3 days before the experiment, mice were habituated to saline intraperitoneal (i.p.) injections and explored a 15cm x 15cm modified open field box for 10 min. The sides of the box were wrapped in 14G copper wire attached to tuner and power distribution control (PDC) boxes (NeuroLux, Inc). Implanted wireless mechano-acoustic (MA) devices (NeuroLux, Inc) were also tested during this time along with power input from the PDC box (7 watts). During each experiment day, the animal explored the enclosure for 2-min before an i.p. injection of either saline, 1 mg/kg clozapine-n oxide (CNO, Enzo Life Sciences), or 5mg/kg CNO. Recordings were taken with a web-camera positioned directly above the animal for 20-min. The average latency to freezing in all CNO trials was used to time-align “drug onset” in saline trials. Each mouse underwent 3-6 saline trials and 2-3 CNO trials with no more than one trial per day. For brain-wide TRAP experiments, a total of 18 CNO trials and 20 saline trials was analyzed from n = 7 Gq+ mice (3 females 4 males) in **Fig 4**, and a total of 17 CNO trials and 17 saline trials was analyzed from n = 6 Gi+ mice (2 females, 4 males) in **Fig S4**. For LPB TRAP experiments, a total of 29 CNO trials and 47 saline trials was analyzed from n = 10 Gq+ mice (2 females, 8 males) in **Fig 7**, and a total of 21 CNO trials and 31 saline trials was analyzed from n = 7 Gi+ mice (2 females, 5 males) in **Fig S6**. Behaviors for a sub-selection of animals were manually scored to identify commonly observed behaviors, as well as determine criteria for behavioral classifiers. Video recordings collected during the experiment were used for pose estimation and supervised behavioral analysis (see *Modified open field test behavioral analysis)* and analyzed for changes in maximum velocity using ANY-maze video tracking software (**Fig S4** and **Fig S6**). Data collected from the MA devices were processed with custom Python scripts. GraphPad Prism was used to analyze group means and paired t-tests, and to analyze data with 2-way ANOVA and Tukey multiple comparison post-tests. See **Table S3** and **Table S5** for statistics.

##### Perfusion for tissue processing

After all studies, mice were deeply anesthetized with isoflurane and underwent transcardial perfusion with phosphate-buffered saline (PBS) followed by 10% formalin and dissection of the brain for further analysis (see *Intact brain clearing and immunolabeling)*.

##### Modified open field test behavioral analysis

We created pose estimation models to track 9 body-part key-points using SLEAP^3^ (snout, left ear, right ear, left lateral, right lateral, centroid, tail base, tail center, and tail-tip). Missing pose-estimation data was interpolated using forward fill and the data was smoothed using a Savitsky-Golay filter across a 500ms sliding window. Grooming, rearing, and Straub tail were detected using random forest classifiers^4^. Freezing and circling were detected using heuristic rules as previously described^32,33^.

*Electrocorticography surgical procedure & neural signal data recording:*

Electrocorticography (ECoG) was recorded using a tethered implant as previously described^29^. Briefly, using aseptic stereotaxic surgical techniques under isoflurane anesthesia, a midline incision was made above the skull, and ECoG-recording electrodes (Pinnacle Technology, Lawrence, KS; No. 8209: 0.10-in.) were screwed through cranial holes at the following coordinates:

- *Channel 1 (left frontal cortex)*: 1.5 mm lateral and 2 mm anterior to bregma
- *Channel 2 (rightparietal cortex)*: 1.5 mm lateral and 2 mm posterior to bregma
- *Ground (visual cortex)*: 1.5 mm lateral and 4 mm posterior to bregma
- *Reference (cerebellum)*: 1.5 mm lateral and 6 mm posterior to bregma

Mice were singly housed in standard cages under a 12:12 light-dark (LD) cycle following surgery. After a 14-day recovery period, they were moved to recording cages, fitted with a tether and preamplifier (100x signal gain) connected to the Pinnacle Technology recording system, and given at least 1 day to acclimate before recording.

Animals were injected with saline or CNO (5mg/kg) at the same time of day on consecutive days (days 1 and 2, respectively), with pre-experimental (day 0) and post-experimental (day 3) days serving as baseline comparisons. To assess neural dynamics during isoflurane, mice were placed in a specialized chamber and induced with 3% isoflurane until fully immobile (∼10 minutes). Isoflurane was then discontinued, and the mice were allowed to fully wake (∼20–30 minutes). In all recording days, electromyography (EMG) signals were obtained by inserting a pair of silver wires into the neck muscles. All screws and wires were connected to a common 6-pin connector compatible with the Pinnacle recording device using silver wire. Signals were continuously recorded at 400 Hz with low-pass filters at 100 Hz. All recordings were saved and exported for processing using the European Data Format (EDF). Simultaneously, animal activity was continuously recorded using webcams.

For each animal, one hour of data (time-matched across days) was extracted for each condition (pre, saline, CNO, post), along with the entirety of the isoflurane induction condition, and saved to individual .edf files. Spectrograms were generated by low-pass filtering each file at 12 Hz and log-transforming the signal data. Additionally, slow wave signal-to-noise ratio (SNR) was calculated using minute-long bins by dividing mean power in the delta frequency band (1–4 Hz, primarily capturing delta waves associated with sleep and anesthesia^35^) by mean power in surrounding frequencies (0.5–1 Hz + 4–7 Hz). All analyses were conducted using the left frontal cortex electrode (i.e., channel 1).

To quantify the change in delta (1-4 Hz) over the course of drug onset (i.e., 1 minute prior to and 15 minutes following drug onset), we ran a linear mixed model predicting delta activity ratio using the interaction term *time × condition* (using the ‘Pre’ condition as reference and controlling for the random effect of individual animal). The results show that the only significant differences in delta activity were the increases seen over time in the CNO and ISO conditions, with no other significant direct or interaction effects (see **Table S3**).

##### Head-fixed implantation and habituation

For high density silicon probe (Neuropixels) recordings, mice were implanted with a custom-made titanium headplate (H.E. Parmer Company, TN) for head-fixation of the animal, and 3D-printed “well”^36^ providing safe access for probe insertion and retention of saline solution on the skull surface during recordings. Following recovery (∼2 weeks), mice were habituated to head-fixation for at least 3 days, starting with 15-to 20-min periods and increasing daily by 20 min until they exhibited minimal distress with fixation for 1 h^37^. For head-fixed recordings, mice were placed in a 3D-printed mouse holder^38^ adapted for isoflurane delivery via a 3D-printed stereotax anesthetic nose cone. During recordings, isoflurane gas concentration was measured with a gas analyzer (BC Biomedical, MO).

##### High-density in vivo electrophysiology

At least 1 day prior to each recording, craniotomies were made targeting regions of interest, planned beforehand using a specialized tool^39^ to take into consideration optimal probe orientations and angles of entry (5-15°), and finally sealed with Kwik-Cast (World Precision Instruments, FL). When not sealed, craniotomies were kept moist with continuous applications of saline into the “well.” On recording days, mice were first head-fixed and allowed to acclimate before probe insertion. Neuropixels probes were mounted on a micromanipulator system (Sensapex, SC) and lowered via the planned trajectory to 100 µm beyond the target depth, then retracted after a 10-min delay and allowed to rest for an additional 10 min before starting the baseline recording. All recordings were acquired using SpikeGLX (Janelia Research Campus, VA). Recordings (**Fig 6A**) began with a 5-min awake baseline period, followed by isoflurane exposure (ISO, 1.3% target concentration on the vaporizer) maintained for at least 20 min. During induction, mice were kept warm via external heating, and loss-of-reflex (LOR) was verified via intermittent foot pinches. Following the 20-min ISO phase, the vaporizer was turned off to initiate a 25-min washout period, followed by a second 5-min awake baseline recording. Mice were subsequently injected with CNO (5 mg/kg, i.p.) to induce activation of isoflurane-tagged Gq^+^ neural ensembles, and recorded for an additional 30 min post-injection to monitor onset response activity. All recordings were acquired using SpikeGLX (Janelia Research Campus, VA).

##### Histological alignment of recording tracts

To determine the placement of isolated units in the brain, sorted units were aligned to the probe tracts after intact brain imaging. After recordings, intact formalin-fixed brains were cleared and imaged with light sheet fluorescent microscopy for probe tract tracing in the Allen Brain Atlas space and mapped onto the locations of isolated units to identify their placement in the brain (**Fig 6B** and see *Intact brain clearing* and *imaging*). After registering each brain to the Allen CCF^11^, probe tracts made by application of the lipophilic CM-DiI dye (Thermo Fisher, WA) to the probe shank before each recording were traced in Lasagna^10^ to yield a fitted line of XYZ coordinates, which were visualized in Brainrender^11^. A separate open-source tool^40^ was used to adjust the alignment of activity recorded along the shank to the regions identified along the tract.

##### Electrophysiological data analysis

Spike sorting was performed with Kilosort4^9^, isolated units were discarded if they had fewer than 500 spikes across the recording and filtered out further based on quality metrics assessment^41^. Quality metric thresholds were adjusted from standard defaults to account for neurons rendered silent during burst suppression epochs, as well as interrupted spike-train continuity leveraged for waveform drift correction, a potential cause for over-splitting^42,43^. Resulting units were then inspected for post-hoc merging during manual curation. Postprocessing and curation were performed using custom scripts integrating open-source Python packages^44^. Spike times were binned (250 ms) and per-sec firing rates were log10-transformed, then z-scored relative to the initial baseline period, with a regularization term added to the standard deviation to stabilize low-firing units. For classification of responses, each unit was classified as activated, suppressed, or neutral for ISO and CNO based on their change in z-scored activity (delta Z) during a post-onset period, relative to pre-onset (ISO: -3 min to ISO onset vs. 3 to 6 min post ISO onset;

##### CNO

5 min to CNO injection vs. 15 to 20 min post CNO injection). Within each comparison period, windows were re-binned into 5 s bins and compared per-unit using Wilcoxon signed-rank tests. FDR-corrections (Benjamini-Hochberg) were applied across all units, which were considered “responsive” at q < 0.05, and defined as “activated” vs. “suppressed” depending on the sign of their delta Z. To establish significant response rates above chance, we first obtained empirical nulls via circular-shift permutations for comparison (500 shuffles), shifting each unit’s combined pre-and post-period window bins by a random offset (between 10-90% of window), to disrupt the alignment of activity to experimental onset periods while maintaining intrinsic temporal dynamics. The classification pipeline was then re-applied to obtain the null distributions of response proportions for each analyzed brain region. Observed responsiveness rates were then determined to be significant if they exceeded these region-specific null rates (one-tailed exact binomial tests), and whether responsive populations exhibited a majority of activated vs. *s*uppressed units (two-tailed binomial tests; p=0.5). For each region, to determine whether ISO-and CNO-responsiveness co-occurred above chance, we tested overlaps based on Fisher’s exact tests on 2x2 conditional tables (ISO+ vs. ISO-vs. CNO+ vs. CNO-), across each region’s units. Overlaps are reported as conditional odds ratio with 95% confidence intervals, and confirmed using permutation tests shuffling CNO-responsiveness labels across units (10k iterations). Among units responsive to both ISO and CNO, whether activated (A) or suppressed (S), we applied Fisher’s exact tests on their direction contingency table (ISO-CNO: A-A vs. A-S vs. S-A vs. S-S) to determine whether response direction was maintained, with same-direction pairs (A-A, S-S) over-represented compared to opposite-direction (A-S, S-A). To determine whether responsiveness overlap was particularly enriched in PB relative to non-PB hindbrain, we fit a logistic model across all hindbrain units predicting ISO responsiveness from their CNO responsive label and whether they were in PB, to test whether the overlap in the latter was stronger than in neighboring hindbrain regions. Given small unit counts for the region, significance was defined using a penalized likelihood-ratio test. All analyses were conducted using custom Python scripts integrating open-source packages.

---

**Figure S1.**
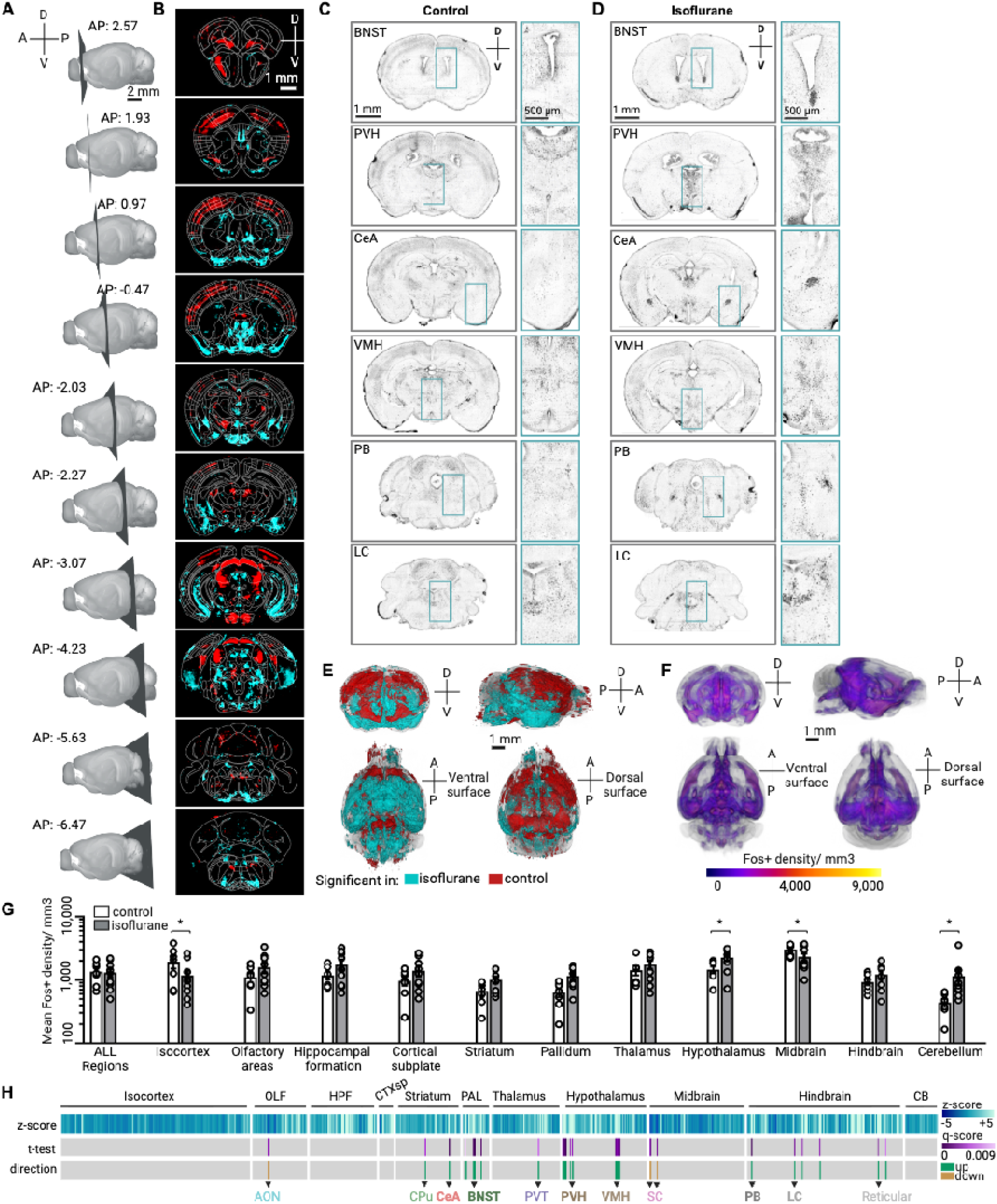
ClearMap analysis and Raw Fos images show subcortical isoflurane-activated hotspots. (**A**) Intact whole brains were imaged horizontally with light sheet microscopy to acquire cellular resolution Fos+ signal, then digitally resliced in the coronal plane at the indicated AP positions. (**B**) Voxel-wise comparison of control vs isoflurane conditions, with cyan indicating higher Fos+ cell densities in the isoflurane condition and red indicating higher Fos+ cell densities in the control condition. (**C**) Representative mean intensity projections of the raw Fos+ signal (3.6x objective) from the control condition and (**D**) isoflurane condition virtually re-sliced to the coronal plane at the indicated significant regions, from top to bottom: BNST = bed nucleus of stria terminalis; PVH = paraventricular hypothalamus; CeA = central amygdala; VMH = ventromedial hypothalamus; PB = parabrachial nucleus; LC = locus coeruleus. (**E**) Voxel-wise significance map of areas significantly regulated in the isoflurane (blue) or control (red) conditions. (**F**) 3D mean Fos+ density heatmaps from the isoflurane condition (also shown in Fig 1B-C). (**G**) Overall mean Fos+ density, calculated across major brain subdivisions after atlas registration by ClearMap in control vs. isoflurane conditions, identifies significant differences in the isocortex, hypothalamus, midbrain and cerebellum (unpaired t-test, *p<0.05). (**H**) Pairwise analysis of z-scored Fos density by each distinct anatomical subregion after FDR correction (q < 0.01) reveals the most significant subregions. Direction of effect in isoflurane relative to control is shown in the bottom bar with green indicating regions of enhanced Fos in isoflurane. Select regions are highlighted (AON: anterior olfactory nucleus, CPu: caudate-putamen, CeA: central amygdala; BNST: bed nucleus of stria terminalis; PVT: paraventricular thalamus; PVH: paraventricular hypothalamus; VMH: ventromedial hypothalamus; SC: superior colliculus; PB: parabrachial nucleus; LC: locus coeruleus). See Table S1 for a full list of regions from pairwise analysis, all raw and z-scored Fos data.

**Figure S2.**
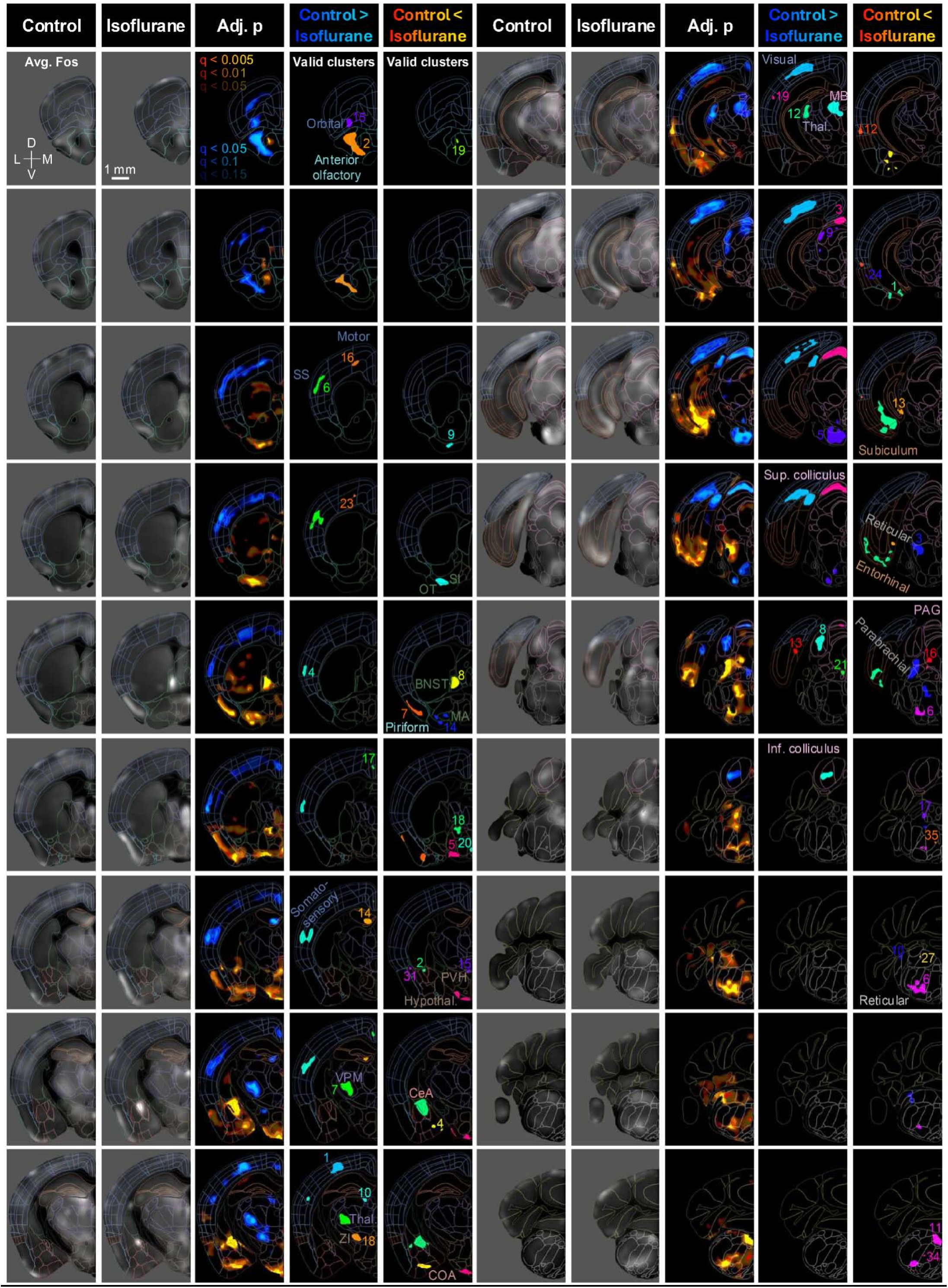
Identification of isoflurane-activated clusters from UNRAVEL analysis. Voxel-wise data from **Fig 1D-G** were warped to the Allen Mouse Brain Common Coordinate Framework (CCFv3) for display in coronal brain slices and overlaid with a colored wireframe using FSLeyes. Input images for voxel-wise comparisons were averaged for each group to show background-subtracted and z-scored Fos immunofluorescence. FDR-corrected p-value maps show increases (warm colors) and decreases (cool colors) in Fos expression resulting from isoflurane treatment. For each effect direction, the most stringent q-value threshold is fully opaque (100%), the next most stringent is semi-opaque (66%), and the least stringent is less opaque (33%). Valid clusters with significant differences in Fos+ cell density were randomly colored and labeled with their IDs. Abbreviations: Adj. = adjusted; Avg. = average; SS = somatosensory; OT = olfactory tubercle; SI = substantia innominata; BNST = bed nucleus of the stria terminalis; MA = magnocellular nucleus; PVH = paraventricular hypothalamic nucleus; Hypothal. = hypothalamus; CeA = central amygdala; Thal. = thalamus; ZI = zona incerta; COA = cortical amygdalar area; MB = midbrain; Sup. = superior; PAG = periaqueductal gray; Inf. = inferior. Please see **Table S1** for a full list of abbreviations and detailed cluster statistics.

**Figure S3.**
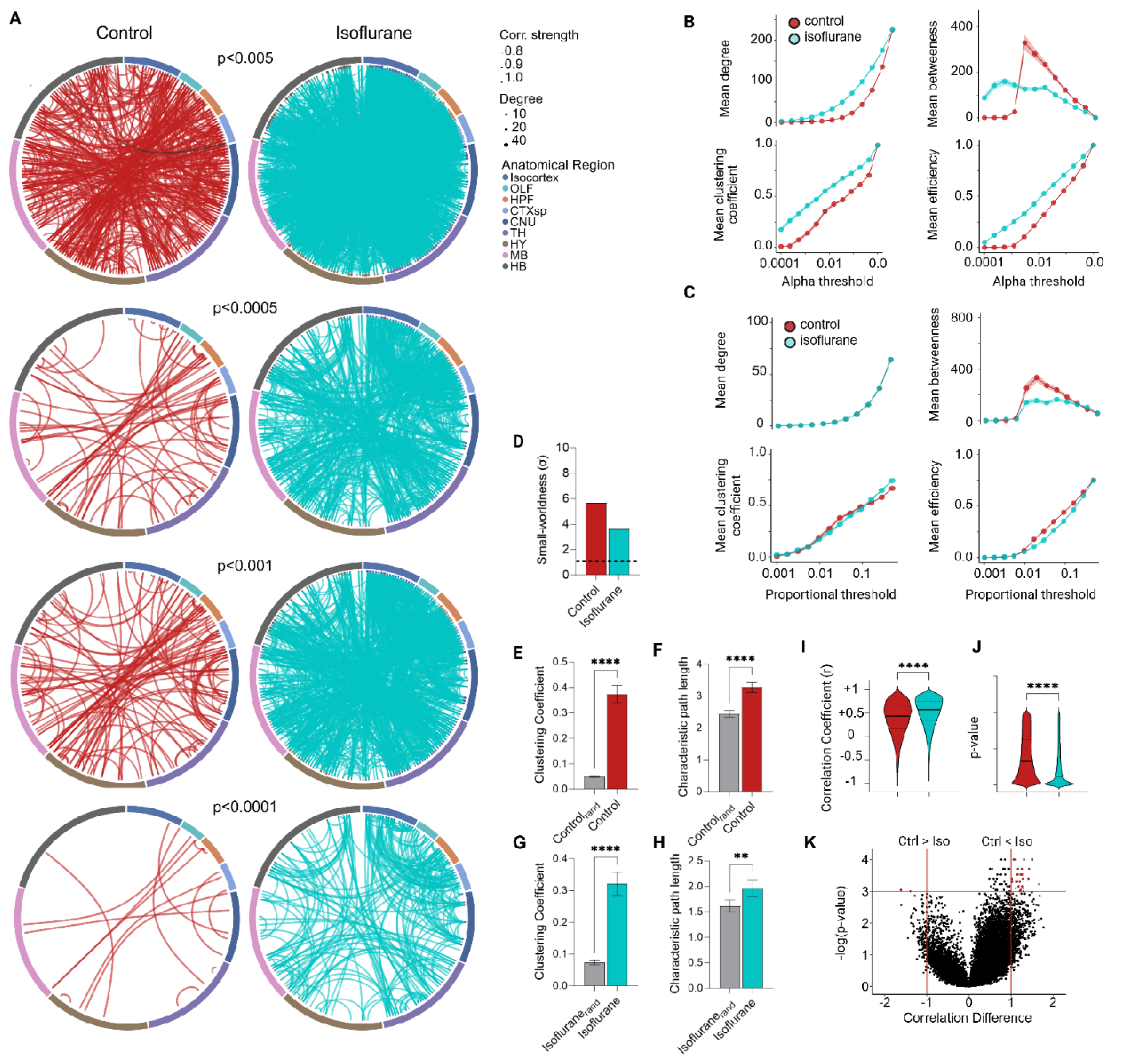
Isoflurane networks are denser across a variety of weight thresholds. (A) control (red) and isoflurane (blue) networks thresholded at various alpha values. (**B**) Mean degree, clustering, betweenness, and efficiency metrics calculated across a range of significance thresholds and (**C**) edge proportion thresholds. (**D**) Assessment of small-world properties of individual networks plotted in Fig 2. Small-world index (o) for both networks are above the threshold of 1 (black dotted horizontal line). The control network (**E**) clustering coefficient and (**F**) path length as well as the isoflurane (**G**) clustering coefficient and (**H**) path length are higher than randomly rewired networks with matching degree sequences. (**I**) Distributions of the Pearson correlation coefficients between all unique regional connections and their associated (**J**) p-value for the control and isoflurane networks. (**K**) Volcano plot illustrating the regional correlations which most differ between the networks. The horizontal red line denotes a significance threshold of p < 0.001. Red points in the upper right (control < isoflurane) and upper left quadrants (control > isoflurane), defined by vertical red line lines, indicate regional correlation differences with a magnitude shift greater than 1 between groups. Two-tailed Student’s t-test: **p<0.01, ****p<0.0001. Error bars represent mean ± SEM. See **Table S2** for full abbreviations and detailed statistics.

**Supplemental Figure S4:**
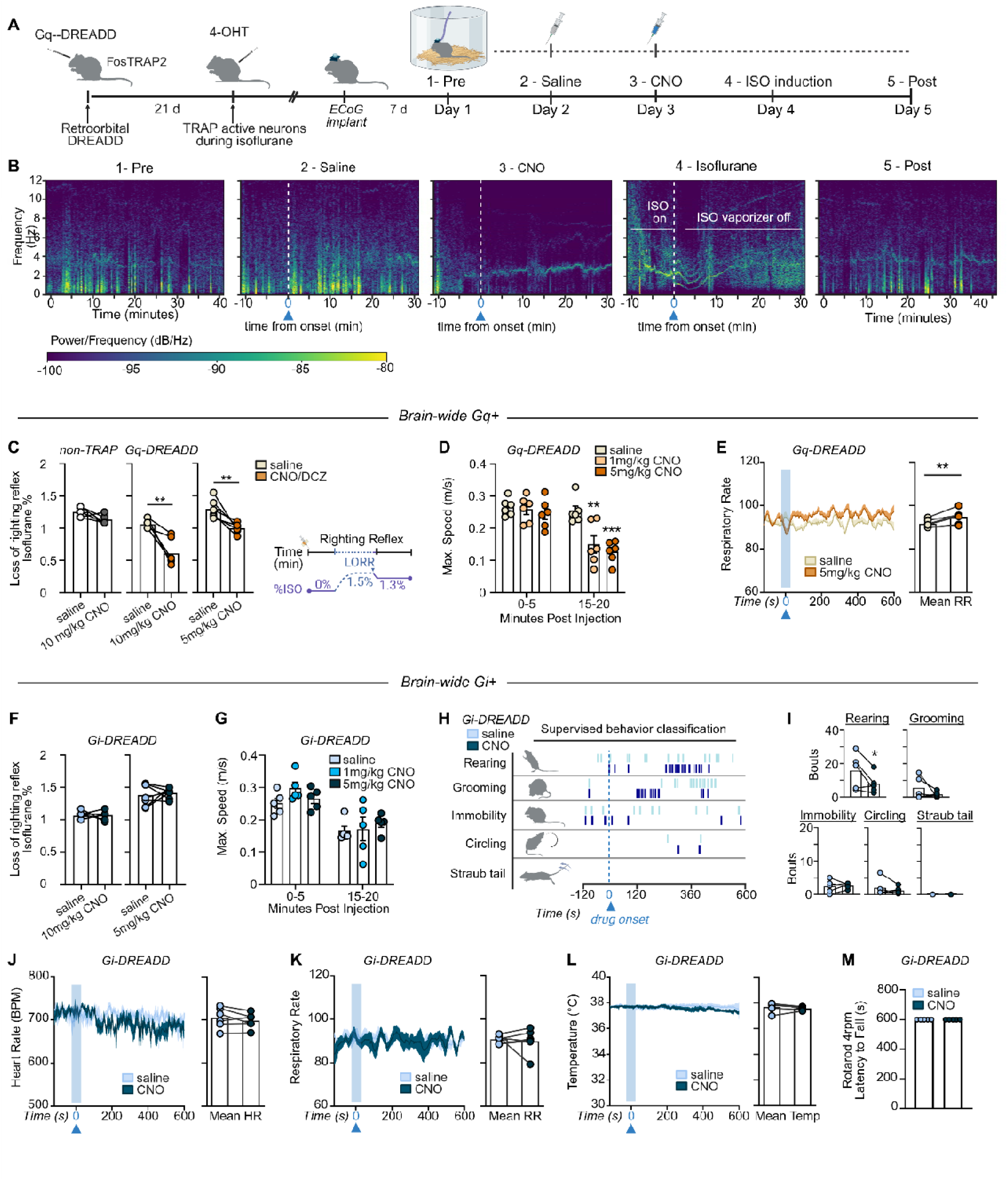
Electrocorticography and brain-wide chemogenetic behaviors. (**A**) Experimental timeline for electrocorticography recordings: FosTRAP2 transgenic mice received retroorbital AAV injection expressing Gq-DREADD and activity-dependent capture during isoflurane exposure, followed by neural electrocorticography (ECoG) recordings over five days (DREADD = designer receptor engineered to be activated by a designer drug, 4-OHT = 4-hydroxytamoxifen, ECoG = electrocorticography, CNO = clozapine n-oxide, ISO = isoflurane). (**B**) Day 1 (labeled “1 -Pre”) shows the average 40-minute baseline recording without manipulations. Recordings on Days 2-4 were time-aligned to the approximated onset of intraperitoneal CNO (10 min post-injection) indicated by the white dotted line. Day 5 (labeled “5 -Post”) was obtained without drug manipulation. The spectrograms shown are an average of 4 mice. (**C**) The concentration of isoflurane (%), as measured with a gas analyzer sampling from the chamber, at which mice display loss of righting reflex during saline or CNO (10 mg/kg and 5 mg/kg) is significantly reduced in Gq-DREADD but not non-TRAP controls that received oil injection instead of 4-OHT (n = 5-6 mice). (**D**) CNO induces dose-dependent reduction in maximum speed (m/s) in the open field in Gq-DREADD mice. (**E**) CNO increases mean respiratory rate in Gq-DREADD mice. (**F**) The concentration of isoflurane (%) at which Gi-DREADD mice display loss of righting reflex during saline or CNO (10 mg/kg, 5mg/kg) is unchanged. (**G**) Maximum speed (m/s) is unchanged in Gi-DREADD mice. (**H**) Representative raster plot from all 6 (3 saline, 3 CNO) of one Gi-DREADD animal’s trials after analysis of classified behavior. (**I**) Bouts of classified behavior for the average of saline and CNO (5 mg/kg) trials per animal expressing inhibiting Gi-DREADD virus (n = 5 mice). There is no change in (**J**) heart rate (BPM, beats per minute) (**K**) respiratory rate (breaths per minute) and (**L**) temperature (Celsius) as recorded from the wireless mechano-acoustic device during trials shown in H-I (n = 6 mice, 2-3 trials per drug condition). (**M**) Rotarod (4-rpm) performance in Gi-DREADD mice after 5 mg/kg CNO (n = 5 mice) is unchanged. Error bars show mean ± SEM. 2-way repeated measures ANOVA with Tukey’s post-hoc tests or paired t-tests, *p<0.05, **p<0.01, ***p<0.001. See **Figure 4** and **Methods** for experimental details. See **Table S3** for detailed statistics.

**Supplemental Figure S5:**
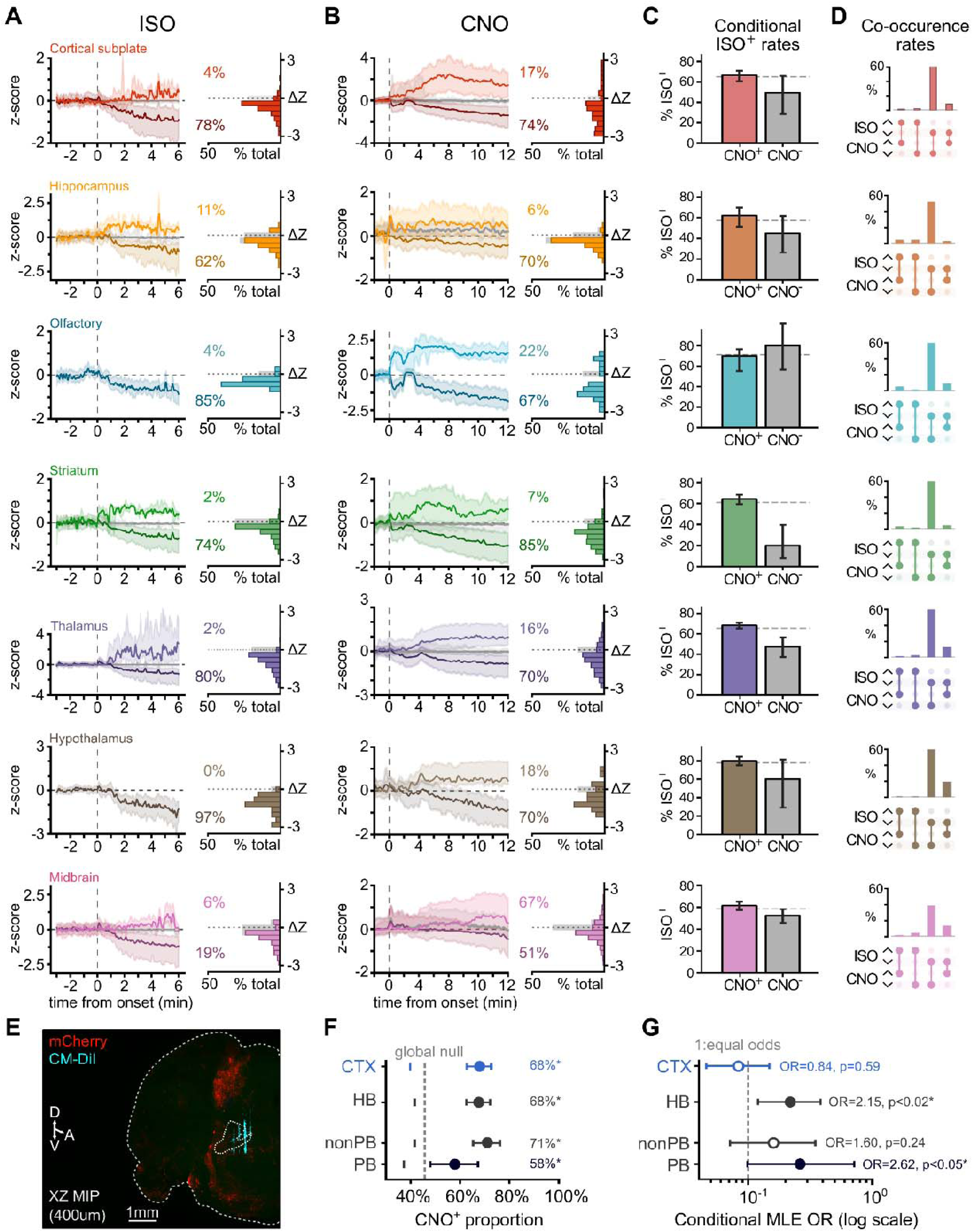
Brain-wide TRAP Neuropixels Supplemental Data. (**A**) ISO-response panels and (**B**) CNO-response panels for regions not included for analysis in Fig 6. Left: mean baseline-normalized logZ Z firing rate traces (“z-score”) for units significantly activated (top, lighter) or suppressed (bottom, darker) by isoflurane, shown pre vs. post onset. Right: distribution of single-unit response magnitudes as a change in their baseline-normalized firing rate (ΔZ, post -pre), as a % of total region population; gray bars denote neutral response distribution. (**C**) Percentage of units that were ISO-responsive, computed separately within CNO-responsive (CNOZ) and CNO-nonresponsive (CNOZ) subgroups (mean ± Wilson 95% CI). (**D**) Upset plots of joint ISO × CNO classification among double-responsive units, showing the percentage of each direction combination (activated / suppressed). (**E**) Representative maximum intensity projection (MIP) across coronal depth (400µm) showing mCherry-DREADD expression (red) and DiI probe tract (cyan) in parabrachial nucleus (inner dashed outline). (**F-G**) Forest plots illustrating CI and detailed statistics for (**F**) proportion tests against region and global empirical shuffle nulls and (**G**) conditional proportion tests for odds ratio of ISO+ given CNO+ / CNO-(log scale; dashed line at OR = 1, equal odds). See also Table S4 for detailed counts and statistics. FR: firing rate; OR: odds ratio.

**Supplemental Figure S6:**
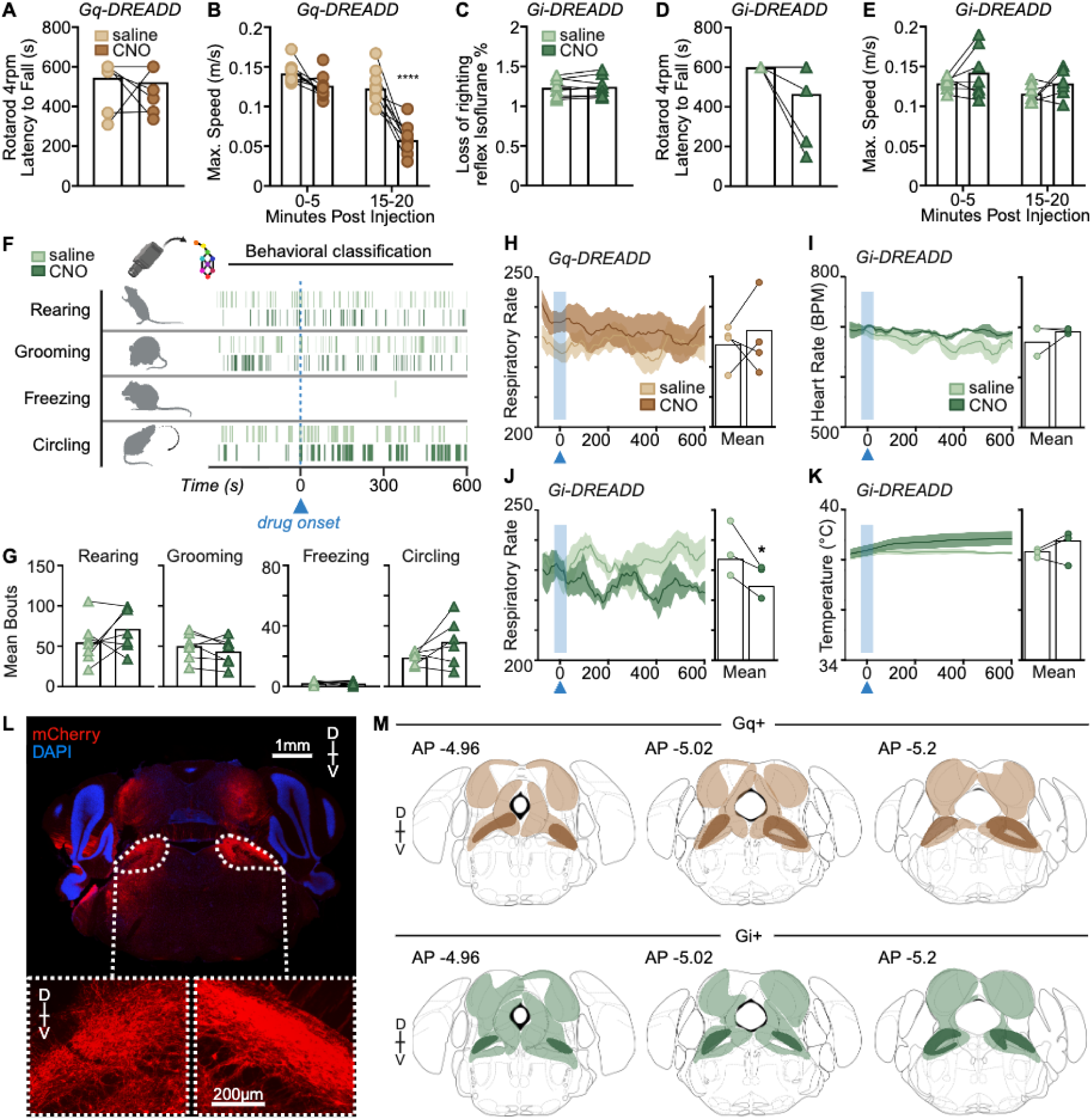
Supplemental Figure S6: (**A**) Latency to fall (s) in Gq-DREADD expressing mice is unchanged following CNO injection (5 mg/kg, n = 10). (**B**) Maximum speed (m/s) is significantly reduced in Gq-DREADD mice 15-20 min post-CNO (5 mg/kg, n = 10). There is no change in (**C**) loss of righting reflex (% isoflurane), (**D**) latency to fall (s) and (**E**) maximum speed (m/s) in Gi-DREADD mice following CNO injection (5 mg/kg, n = 7). (**F**) A representative raster plot from one animal after analysis of classified behavior (also see **Video S5**) after CNO injection as compared to saline in Gi-DREADD mice. (**G**) Bouts of classified behavior for the average of saline and CNO trials per animal (n = 7, total of 21 CNO and 31 saline trials) expressing inhibiting Gi-DREADD virus. CNO had no effect on rearing, grooming, freezing, or circling in Gi-DREADD mice. (**H**) Respiratory rate in Gq-DREADD mice is unchanged after CNO injection as compared to saline (n = 4). (**I**) Heart rate (BPM, beats per minute) is unchanged following CNO in Gi-DREADD mice (n = 3). (**J**) Respiratory rate is decreased following CNO in Gi-DREADD mice (n = 3). (**K**) Temperature (Celsius) is unchanged following CNO in Gi-DREADD mice (n = 3). (**L**) Representative image after mCherry-immunolabeling for DREADD expression and DAPI staining. Scale bar is 1 mm and 200 µm in inset. (M) Representative brain sections showing DREADD expression in Gq-(top) and Gi-(bottom) DREADD expressing mice. Highlighted regions indicate areas of viral expression. See **Figure 7** and **Methods** for experimental details. Error bars show mean ± SEM. Mixed effects analysis with Sidak’s multiple comparisons (panel b, e) and paired t-tests, *p<0.05, ****p<0.0001. See **Table S5** for statistics.

**Figure S7.**
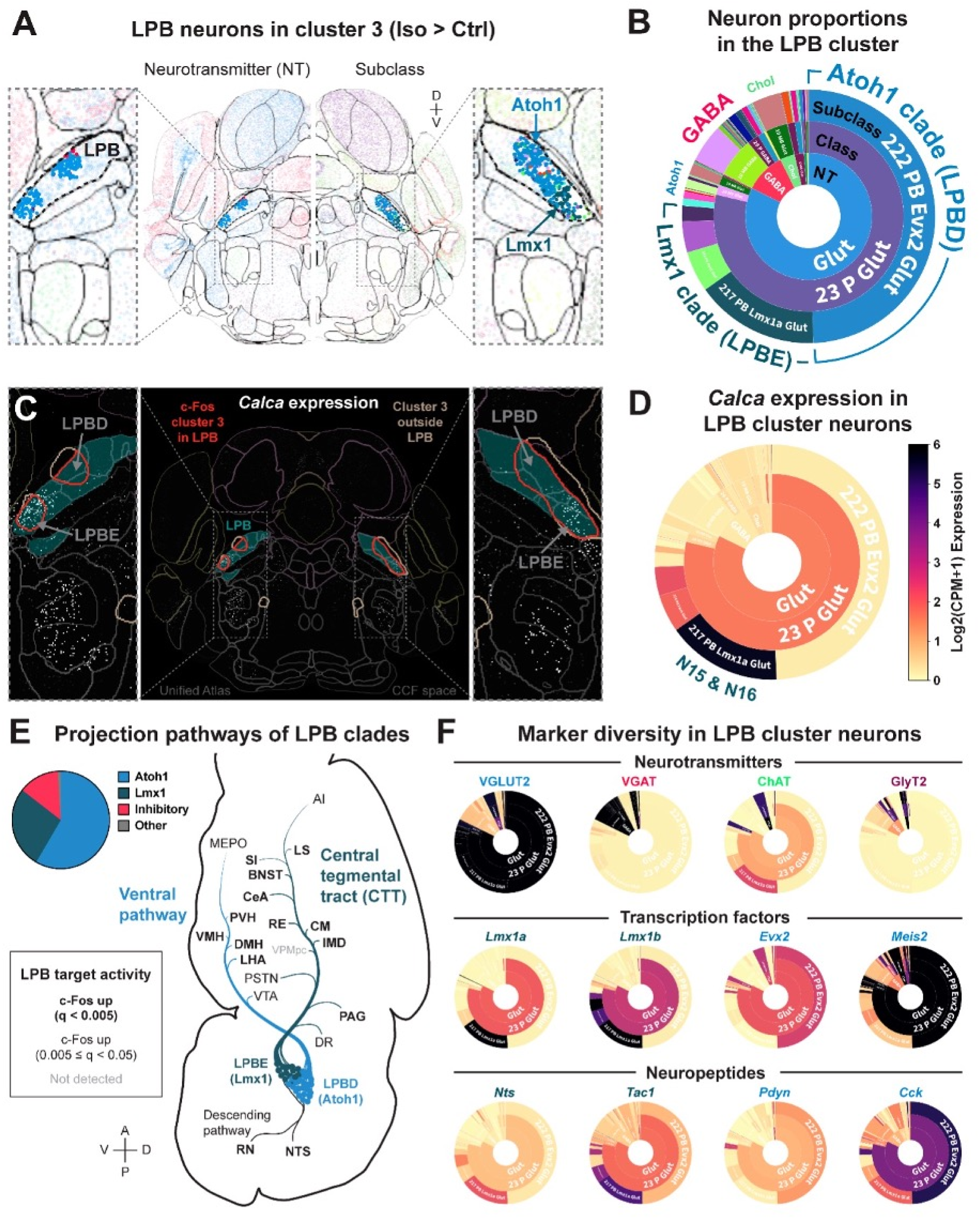
Diversity of neurons in LPB cluster 3. (**A**) Neuron types from the Allen Brain Cell Atlas (ABCA) within the LPB portion of Control < Isoflurane cluster 3 (from **Fig. 1D-G**) are highlighted in MERFISH-CCF space. For this, c-Fos clusters were warped to ABCA MERFISH-CCF space. The spatial distributions of *Atoh1-* and *Lmx1*-derived LPB clades, described by Pauli et al. (2022)51, are noted for reference. (**B**) ABCA neuron type proportions in the LPB portion of cluster 3. (**C**) To examine the alignment of cluster 3 with LPB subregions, the Unified Mouse Brain Atlas v2 labels were warped to MERFISH-CCF space. LPBE, external LPB; LPBD, dorsal LPB. Imputed Calca expression in ABCA neurons is shown in the context of LPB subregions and cluster 3; display range, 0-6 logD (CPM + 1). (**D**) Calca marks CGRP-associated LPB neurons, including N15 and N16 subclusters described by Pauli et al. (2022). (**E**) Major LPB projection pathways adapted from Pauli et al. (2022). Atoh1-related LPBD populations project predominantly through the ventral pathway, whereas Lmx1-related LPBE populations project predominantly through the central tegmental tract (CTT). Both have additional descending projections. Clade proportions were estimated using imputed marker expression, with marker-positive neurons defined as >3 logD (CPM + 1): *Atoh1 = Slc17a6+/Meis2+/Lmx1b-, Lmx1 = Slc17a6+/Meis2-/Lmx1b+*, inhibitory = GABAergic neurons, and other = remaining neurons. Target labels indicate c-Fos upregulation in the Control < Isoflurane UNRAVEL analysis: bold, FDR q < 0.005; regular black, 0.005 ≤ q < 0.05; gray, not detected at q < 0.05. Nearly all LPB target regions shown had increased c-Fos, suggesting that isoflurane-activated glutamatergic LPB neurons may contribute to widespread downstream activity. (**F**) Imputed marker expression across ABCA neurons within the LPB portion of cluster 3. *Slc17a6* encodes VGLUT2, *Slc32a1* encodes VGAT, *Chat* encodes ChAT, and *Slc6a5* encodes GlyT2. The transcription factors *Lmx1a* and *Lmx1b* are expressed in the *Lmx1* clade, whereas *Evx2* and *Meis2* are expressed in the *Atoh1* clade. Examples of neuropeptides preferentially associated with *Atoh1-* or *Lmx1*-related LPB populations are shown. Region abbreviations in (**E**): AI, agranular insula; BNST, bed nucleus of the stria terminalis; CeA, central amygdala; CM, central medial thalamus; DMH, dorsomedial hypothalamus; DR, dorsal raphe; IMD, intermediodorsal thalamus; LHA, lateral hypothalamic area; LS, lateral septum; MEPO, median preoptic nucleus; NTS, nucleus of the solitary tract; PAG, periaqueductal gray; PSTN, parasubthalamic nucleus; PVH, paraventricular hypothalamus; RE, nucleus of reuniens; RN, reticular nucleus; SI, substantia innominata; VMH, ventromedial hypothalamus; VPMpc, ventral posteromedial thalamus, parvicellular; VTA, ventral tegmental area.

## REFERENCES

1. Mashour, G.A. (2024). Anesthesia and the neurobiology of consciousness. Neuron 112, 1553–1567. 10.1016/j.neuron.2024.03.002.

2. Brown, E.N., Lydic, R., and Schiff, N.D. (2010). General Anesthesia, Sleep, and Coma. New England Journal of Medicine 363, 2638–2650. 10.1056/NEJMra0808281.

3. Kelz, M.B., and Mashour, G.A. (2019). The Biology of General Anesthesia from Paramecium to Primate. Curr Biol 29, R1199–R1210. 10.1016/j.cub.2019.09.071.

4. Moody, O.A., Zhang, E.R., Vincent, K.F., Kato, R., Melonakos, E.D., Nehs, C.J., and Solt, K. (2021). The Neural Circuits Underlying General Anesthesia and Sleep. Anesth Analg 132, 1254– 1264. 10.1213/ANE.0000000000005361.

5. Zhang, Z., Ferretti, V., Guntan, I., Moro, A., Steinberg, E.A., Ye, Z., Zecharia, A.Y., Yu, X., Vyssotski, A.L., Brickley, S.G., et al. (2015). Neuronal ensembles sufficient for recovery sleep and the sedative actions of α2 adrenergic agonists. Nat Neurosci 18, 553–561. 10.1038/nn.3957.

6. Jiang-Xie, L.-F., Yin, L., Zhao, S., Prevosto, V., Han, B.-X., Dzirasa, K., and Wang, F. (2019). A Common Neuroendocrine Substrate for Diverse General Anesthetics and Sleep. Neuron 102, 1053–1065.e4. 10.1016/j.neuron.2019.03.033.

7. Wasilczuk, A.Z., Takekawa, D., Cichon, J., Zhang, X., Blackwood, E.B., Parra-Munevar, J., Kim, M.-H., Spain, W., Krom, A.J., Kelz, M.B., et al. (2025). Convergent Control of NREM Sleep and Anesthesia by Prefrontal Layer 5 Extratelencephalic Neurons. Preprint at Research Square, 10.21203/rs.3.rs-7861434/v1 https://doi.org/10.21203/rs.3.rs-7861434/v1.

8. Nelson, L.E., Guo, T.Z., Lu, J., Saper, C.B., Franks, N.P., and Maze, M. (2002). The sedative component of anesthesia is mediated by GABA(A) receptors in an endogenous sleep pathway. Nat Neurosci 5, 979–984. 10.1038/nn913.

9. Taylor, N.E., Van Dort, C.J., Kenny, J.D., Pei, J., Guidera, J.A., Vlasov, K.Y., Lee, J.T., Boyden, E.S., Brown, E.N., and Solt, K. (2016). Optogenetic activation of dopamine neurons in the ventral tegmental area induces reanimation from general anesthesia. Proc Natl Acad Sci U S A 113, 12826– 12831. 10.1073/pnas.1614340113.

10. Moruzzi, G., and Magoun, H.W. (1949). Brain stem reticular formation and activation of the EEG. Electroencephalogr Clin Neurophysiol 1, 455–473.

11. Mashour, G.A., Pal, D., and Brown, E.N. (2022). Prefrontal cortex as a key node in arousal circuitry. Trends Neurosci. 10.1016/j.tins.2022.07.002.

12. Mashour, G.A., Palanca, B.J., Basner, M., Li, D., Wang, W., Blain-Moraes, S., Lin, N., Maier, K., Muench, M., Tarnal, V., et al. (2021). Recovery of consciousness and cognition after general anesthesia in humans. Elife 10, e59525. 10.7554/eLife.59525.

13. Palmiter, R.D. (2011). Dopamine signaling as a neural correlate of consciousness. Neuroscience 198, 213–220. 10.1016/j.neuroscience.2011.06.089.

14. Melonakos, E.D., Siegmann, M.J., Rey, C., O’Brien, C., Nikolaeva, K.K., Solt, K., and Nehs, C.J. (2021). Excitation of Putative Glutamatergic Neurons in the Rat Parabrachial Nucleus Region Reduces Delta Power during Dexmedetomidine but not Ketamine Anesthesia. Anesthesiology 135, 633–648. 10.1097/ALN.0000000000003883.

15. Kelz, M.B., Sun, Y., Chen, J., Cheng Meng, Q., Moore, J.T., Veasey, S.C., Dixon, S., Thornton, M., Funato, H., and Yanagisawa, M. (2008). An essential role for orexins in emergence from general anesthesia. Proc Natl Acad Sci U S A 105, 1309–1314. 10.1073/pnas.0707146105.

16. Zhou, W., Cheung, K., Kyu, S., Wang, L., Guan, Z., Kurien, P.A., Bickler, P.E., and Jan, L.Y. (2018). Activation of orexin system facilitates anesthesia emergence and pain control. Proc Natl Acad Sci U S A 115, E10740–E10747. 10.1073/pnas.1808622115.

17. Heshmati, M., and Bruchas, M.R. (2022). Historical and Modern Evidence for the Role of Reward Circuitry in Emergence. Anesthesiology 136, 997–1014. 10.1097/ALN.0000000000004148.

18. Hua, T., Chen, B., Lu, D., Sakurai, K., Zhao, S., Han, B.-X., Kim, J., Yin, L., Chen, Y., Lu, J., et al. (2020). General anesthetics activate a potent central pain-suppression circuit in the amygdala. Nat Neurosci 23, 854–868. 10.1038/s41593-020-0632-8.

19. Huang, Z., Mashour, G.A., and Hudetz, A.G. (2024). Propofol disrupts the functional core-matrix architecture of the thalamus in humans. Nat Commun 15, 7496. 10.1038/s41467-024-51837-1.

20. Bharioke, A., Munz, M., Brignall, A., Kosche, G., Eizinger, M.F., Ledergerber, N., Hillier, D., Gross-Scherf, B., Conzelmann, K.-K., Macé, E., et al. (2022). General anesthesia globally synchronizes activity selectively in layer 5 cortical pyramidal neurons. Neuron 110, 2024–2040.e10. 10.1016/j.neuron.2022.03.032.

21. Weinrich, J.A., Liu, C.D., Jewell, M.E., Andolina, C.R., Bernstein, M.X., Benitez, J., Rodriguez-Rosado, S., Braz, J.M., Maze, M., Nemenov, M.I., et al. (2023). Paradoxical increases in anterior cingulate cortex activity during nitrous oxide-induced analgesia reveal a signature of pain affect. Preprint at bioRxiv, 10.1101/2023.04.03.534475 https://doi.org/10.1101/2023.04.03.534475.

22. Zhang, X., Baer, A.G., Price, J.M., Jones, P.C., Garcia, B.J., Romero, J., Cliff, A.M., Mi, W., Brown, J.B., Jacobson, D.A., et al. (2020). Neurotransmitter networks in mouse prefrontal cortex are reconfigured by isoflurane anesthesia. J Neurophysiol 123, 2285–2296. 10.1152/jn.00092.2020.

23. Krom, A.J., Marmelshtein, A., Gelbard-Sagiv, H., Tankus, A., Hayat, H., Hayat, D., Matot, I., Strauss, I., Fahoum, F., Soehle, M., et al. (2020). Anesthesia-induced loss of consciousness disrupts auditory responses beyond primary cortex. Proc Natl Acad Sci USA 117, 11770–11780. 10.1073/pnas.1917251117.

24. Gao, S., Proekt, A., Renier, N., Calderon, D.P., and Pfaff, D.W. (2019). Activating an anterior nucleus gigantocellularis subpopulation triggers emergence from pharmacologically-induced coma in rodents. Nature Communications 10, 2897. 10.1038/s41467-019-10797-7.

25. Redinbaugh, M.J., Phillips, J.M., Kambi, N.A., Mohanta, S., Andryk, S., Dooley, G.L., Afrasiabi, M., Raz, A., and Saalmann, Y.B. (2020). Thalamus Modulates Consciousness via Layer-Specific Control of Cortex. Neuron 106, 66–75.e12. 10.1016/j.neuron.2020.01.005.

26. Hemmings, H.C., Riegelhaupt, P.M., Kelz, M.B., Solt, K., Eckenhoff, R.G., Orser, B.A., and Goldstein, P.A. (2019). Towards a Comprehensive Understanding of Anesthetic Mechanisms of Action: A Decade of Discovery. Trends in Pharmacological Sciences 40, 464–481. 10.1016/j.tips.2019.05.001.

27. Perouansky, M., Johnson-Schlitz, D., Sedensky, M.M., and Morgan, P.G. (2023). A primordial target: Mitochondria mediate both primary and collateral anesthetic effects of volatile anesthetics. Exp Biol Med (Maywood) 248, 545–552. 10.1177/15353702231165025.

28. Baron, M., and Devor, M. (2023). From molecule to oblivion: dedicated brain circuitry underlies anesthetic loss of consciousness permitting pain-free surgery. Front Mol Neurosci 16, 1197304. 10.3389/fnmol.2023.1197304.

29. Suzuki, M., and Larkum, M.E. (2020). General Anesthesia Decouples Cortical Pyramidal Neurons. Cell 180, 666–676.e13. 10.1016/j.cell.2020.01.024.

30. Reitz, S.L., and Kelz, M.B. (2021). Preoptic Area Modulation of Arousal in Natural and Drug Induced Unconscious States. Front. Neurosci. 15. 10.3389/fnins.2021.644330.

31. Ibraheem, A., Vaso, K., Minert, A., Yatziv, S.-L., Baron, M., and Devor, M. (2025). Loss-of-consciousness: sources of GABAergic input to the mesopontine tegmental anesthesia area. Front Neurosci 19, 1594984. 10.3389/fnins.2025.1594984.

32. Koch, C. (2018). What Is Consciousness? Nature 557, S8–S12. 10.1038/d41586-018-05097-x.

33. Mashour, G.A., Roelfsema, P., Changeux, J.-P., and Dehaene, S. (2020). Conscious Processing and the Global Neuronal Workspace Hypothesis. Neuron 105, 776–798. 10.1016/j.neuron.2020.01.026.

34. Dehaene, S., Kerszberg, M., and Changeux, J.-P. (1998). A neuronal model of a global workspace in effortful cognitive tasks. Proc Natl Acad Sci U S A 95, 14529–14534.

35. Madangopal, R., Szelenyi, E.R., Nguyen, J., Brenner, M.B., Drake, O.R., Pham, D.Q., Shekara, A., Jin, M., Choong, J.J., Heins, C., et al. (2022). Incubation of palatable food craving is associated with brain-wide neuronal activation in mice. Proceedings of the National Academy of Sciences 119, e2209382119. 10.1073/pnas.2209382119.

36. Rijsketic, D.R., Casey, A.B., Barbosa, D.A.N., Zhang, X., Hietamies, T.M., Ramirez-Ovalle, G., Pomrenze, M.B., Halpern, C.H., Williams, L.M., Malenka, R.C., et al. (2023). UNRAVELing the synergistic effects of psilocybin and environment on brain-wide immediate early gene expression in mice. Neuropsychopharmacol. 48, 1798–1807. 10.1038/s41386-023-01613-4.

37. Renier, N., Wu, Z., Simon, D.J., Yang, J., Ariel, P., and Tessier-Lavigne, M. (2014). iDISCO: A Simple, Rapid Method to Immunolabel Large Tissue Samples for Volume Imaging. Cell 159, 896– 910. 10.1016/j.cell.2014.10.010.

38. DeNardo, L.A., Liu, C.D., Allen, W.E., Adams, E.L., Friedmann, D., Fu, L., Guenthner, C.J., Tessier-Lavigne, M., and Luo, L. (2019). Temporal evolution of cortical ensembles promoting remote memory retrieval. Nat. Neurosci. 22, 460–469. 10.1038/s41593-018-0318-7.

39. Allen, W.E., DeNardo, L.A., Chen, M.Z., Liu, C.D., Loh, K.M., Fenno, L.E., Ramakrishnan, C., Deisseroth, K., and Luo, L. (2017). Thirst-associated preoptic neurons encode an aversive motivational drive. Science 357, 1149–1155. 10.1126/science.aan6747.

40. Armbruster, B.N., Li, X., Pausch, M.H., Herlitze, S., and Roth, B.L. (2007). Evolving the lock to fit the key to create a family of G protein-coupled receptors potently activated by an inert ligand. Proceedings of the National Academy of Sciences 104, 5163–5168. 10.1073/pnas.0700293104.

41. Jun, J.J., Steinmetz, N.A., Siegle, J.H., Denman, D.J., Bauza, M., Barbarits, B., Lee, A.K., Anastassiou, C.A., Andrei, A., Aydın, Ç., et al. (2017). Fully integrated silicon probes for high-density recording of neural activity. Nature 551, 232–236. 10.1038/nature24636.

42. Ouyang, W., Kilner, K.J., Xavier, R.M.P., Liu, Y., Lu, Y., Feller, S.M., Pitts, K.M., Wu, M., Ausra, J., Jones, I., et al. (2024). An implantable device for wireless monitoring of diverse physio-behavioral characteristics in freely behaving small animals and interacting groups. Neuron 112, 1764–1777.e5. 10.1016/j.neuron.2024.02.020.

43. Pereira, T.D., Tabris, N., Matsliah, A., Turner, D.M., Li, J., Ravindranath, S., Papadoyannis, E.S., Normand, E., Deutsch, D.S., Wang, Z.Y., et al. (2022). SLEAP: A deep learning system for multi-animal pose tracking. Nat Methods 19, 486–495. 10.1038/s41592-022-01426-1.

44. Goodwin, N.L., Choong, J.J., Hwang, S., Pitts, K., Bloom, L., Islam, A., Zhang, Y.Y., Szelenyi, E.R., Tong, X., Newman, E.L., et al. (2024). Simple Behavioral Analysis (SimBA) as a platform for explainable machine learning in behavioral neuroscience. Nat Neurosci 27, 1411–1424. 10.1038/s41593-024-01649-9.

45. Muindi, F., Kenny, J.D., Taylor, N.E., Solt, K., Wilson, M.A., Brown, E.N., and Van Dort, C.J. (2016). Electrical stimulation of the parabrachial nucleus induces reanimation from isoflurane general anesthesia. Behavioural Brain Research 306, 20–25. 10.1016/j.bbr.2016.03.021.

46. Wang, T.-X., Xiong, B., Xu, W., Wei, H.-H., Qu, W.-M., Hong, Z.-Y., and Huang, Z.-L. (2019). Activation of Parabrachial Nucleus Glutamatergic Neurons Accelerates Reanimation from Sevoflurane Anesthesia in Mice. Anesthesiology 130, 106–118. 10.1097/ALN.0000000000002475.

47. Luo, T., Yu, S., Cai, S., Zhang, Y., Jiao, Y., Yu, T., and Yu, W. (2018). Parabrachial Neurons Promote Behavior and Electroencephalographic Arousal From General Anesthesia. Frontiers in molecular neuroscience 11, 420. 10.3389/fnmol.2018.00420.

48. Xu, Q., Wang, D.-R., Dong, H., Chen, L., Lu, J., Lazarus, M., Cherasse, Y., Chen, G.-H., Qu, W.-M., and Huang, Z.-L. (2021). Medial Parabrachial Nucleus Is Essential in Controlling Wakefulness in Rats. Front Neurosci 15, 645877. 10.3389/fnins.2021.645877.

49. Renier, N., Adams, E.L., Kirst, C., Wu, Z., Azevedo, R., Kohl, J., Autry, A.E., Kadiri, L., Venkataraju, K.U., Zhou, Y., et al. (2016). Mapping of Brain Activity by Automated Volume Analysis of Immediate Early Genes. Cell 165, 1789–1802. 10.1016/j.cell.2016.05.007.

50. Jin, M., Ogundare, S.O., Lanio, M., Sorid, S., Whye, A.R., Leal Santos, S., Franceschini, A., and Denny, C.A. (2025). A SMARTTR workflow for multi-ensemble atlas mapping and brain-wide network analysis. eLife 13, RP101327. 10.7554/eLife.101327.3.

51. Vetere, G., Kenney, J.W., Tran, L.M., Xia, F., Steadman, P.E., Parkinson, J., Josselyn, S.A., and Frankland, P.W. (2017). Chemogenetic Interrogation of a Brain-wide Fear Memory Network in Mice. Neuron 94, 363–374.e4. 10.1016/j.neuron.2017.03.037.

52. Wheeler, A.L., Teixeira, C.M., Wang, A.H., Xiong, X., Kovacevic, N., Lerch, J.P., McIntosh, A.R., Parkinson, J., and Frankland, P.W. (2013). Identification of a Functional Connectome for Long-Term Fear Memory in Mice. PLoS Comput Biol 9, e1002853. 10.1371/journal.pcbi.1002853.

53. Wheeler, A.L., Creed, M.C., Voineskos, A.N., and Nobrega, J.N. (2014). Changes in brain functional connectivity after chronic haloperidol in rats: a network analysis. Int J Neuropsychopharmacol 17, 1129–1138. 10.1017/S1461145714000042.

54. Watts, D.J., and Strogatz, S.H. (1998). Collective dynamics of ‘small-world’ networks. Nature 393, 440–442. 10.1038/30918.

55. Clauset, A., Newman, M.E.J., and Moore, C. (2004). Finding community structure in very large networks. Phys. Rev. E 70, 066111. 10.1103/PhysRevE.70.066111.

56. Chan, K.Y., Jang, M.J., Yoo, B.B., Greenbaum, A., Ravi, N., Wu, W.-L., Sánchez-Guardado, L., Lois, C., Mazmanian, S.K., Deverman, B.E., et al. (2017). Engineered AAVs for efficient noninvasive gene delivery to the central and peripheral nervous systems. Nat Neurosci 20, 1172– 1179. 10.1038/nn.4593.

57. Dickinson, R., White, I., Lieb, W.R., and Franks, N.P. (2000). Stereoselective Loss of Righting Reflex in Rats by Isoflurane. Anesthesiology 93, 837–843. 10.1097/00000542-200009000-00035.

58. Manvich, D.F., Webster, K.A., Foster, S.L., Farrell, M.S., Ritchie, J.C., Porter, J.H., and Weinshenker, D. (2018). The DREADD agonist clozapine N-oxide (CNO) is reverse-metabolized to clozapine and produces clozapine-like interoceptive stimulus effects in rats and mice. Sci Rep 8, 3840. 10.1038/s41598-018-22116-z.

59. Jendryka, M., Palchaudhuri, M., Ursu, D., van der Veen, B., Liss, B., Kätzel, D., Nissen, W., and Pekcec, A. (2019). Pharmacokinetic and pharmacodynamic actions of clozapine-N-oxide, clozapine, and compound 21 in DREADD-based chemogenetics in mice. Sci Rep 9, 4522. 10.1038/s41598-019-41088-2.

60. Hsueh, B., Chen, R., Jo, Y., Tang, D., Raffiee, M., Kim, Y.S., Inoue, M., Randles, S., Ramakrishnan, C., Patel, S., et al. (2023). Cardiogenic control of affective behavioural state. Nature 615, 292–299. 10.1038/s41586-023-05748-8.

61. Amzica, F., and Steriade, M. (1998). Electrophysiological correlates of sleep delta waves. Electroencephalogr Clin Neurophysiol 107, 69–83. 10.1016/s0013-4694(98)00051-0.

62. Hrvatin, S., Sun, S., Wilcox, O.F., Yao, H., Lavin-Peter, A.J., Cicconet, M., Assad, E.G., Palmer, M.E., Aronson, S., Banks, A.S., et al. (2020). Neurons that regulate mouse torpor. Nature 583, 115– 121. 10.1038/s41586-020-2387-5.

63. Huang, Y.-G., Flaherty, S.J., Pothecary, C.A., Foster, R.G., Peirson, S.N., and Vyazovskiy, V.V. (2021). The relationship between fasting-induced torpor, sleep, and wakefulness in laboratory mice. Sleep 44, zsab093. 10.1093/sleep/zsab093.

64. Akeju, O., Hamilos, A.E., Song, A.H., Pavone, K.J., Purdon, P.L., and Brown, E.N. (2016). GABAA circuit mechanisms are associated with ether anesthesia-induced unconsciousness. Clinical Neurophysiology 127, 2472–2481. 10.1016/j.clinph.2016.02.012.

65. Pauli, J.L., Chen, J.Y., Basiri, M.L., Park, S., Carter, M.E., Sanz, E., McKnight, G.S., Stuber, G.D., and Palmiter, R.D. (2022). Molecular and anatomical characterization of parabrachial neurons and their axonal projections. eLife 11, e81868. 10.7554/eLife.81868.

66. Vanini, G., Bassana, M., Bower, M., Mondino, A., Cerda, I., Phyle, M., Chen, V., Colmenero, A.V., Hambrecht-Wiedbusch, V.S., and Mashour, G.A. (2020). Activation of Preoptic GABAergic or Glutamatergic Neurons Modulates Sleep-Wake Architecture But Not Anesthetic State Transitions. Current biologyD: CB 30, 779. 10.1016/j.cub.2019.12.063.

67. Jung, S., Kondaurova, A., Lee, M., Kwon, J.J., Shin, H., and Kim, S.-Y. (2026). Parabrachial neurons orchestrate cold-induced homeostatic responses. Nat Metab 8, 1679–1693. 10.1038/s42255-026-01565-1.

68. Yahiro, T., Kataoka, N., and Nakamura, K. (2023). Two Ascending Thermosensory Pathways from the Lateral Parabrachial Nucleus That Mediate Behavioral and Autonomous Thermoregulation. J. Neurosci. 43, 5221–5240. 10.1523/JNEUROSCI.0643-23.2023.

69. Yang, W.Z., Du, X., Zhang, W., Gao, C., Xie, H., Xiao, Y., Jia, X., Liu, J., Xu, J., Fu, X., et al. (2020). Parabrachial neuron types categorically encode thermoregulation variables during heat defense. Sci Adv 6, eabb9414. 10.1126/sciadv.abb9414.

70. Machado, N.L.S., Raffin, F., Kaur, S., Banks, A.S., Lynch, N., Fanari, O., Plascencia, O.R., Aten, S., Lima, J.D., Bandaru, S.S., et al. (2023). Prolonged activation of EP3 receptor-expressing preoptic neurons underlies torpor responses. Res Sq, rs.3.rs-2861253. 10.21203/rs.3.rs-2861253/v1

71. Wasilczuk, A.Z., Rinehart, C., Aggarwal, A., Stone, M.E., Mashour, G.A., Avidan, M.S., Kelz, M.B., Proekt, A., and ReCCognition Study Group (2024). Hormonal basis of sex differences in anesthetic sensitivity. Proceedings of the National Academy of Sciences 121, e2312913120. 10.1073/pnas.2312913120.

72. Suntsova, N., Guzman-Marin, R., Kumar, S., Alam, M.N., Szymusiak, R., and McGinty, D. (2007). The Median Preoptic Nucleus Reciprocally Modulates Activity of Arousal-Related and Sleep-Related Neurons in the Perifornical Lateral Hypothalamus. J. Neurosci. 27, 1616–1630. 10.1523/JNEUROSCI.3498-06.2007.

73. Zhang, J.-S., Yao, W., Zhang, L., Li, Z.-S., Gong, X.-T., Duan, L.-L., Huang, Z.-X., Chen, T., Huang, J.-C., Yang, S.-X., et al. (2025). GABAergic Neurons in the Central Amygdala Promote Emergence from Isoflurane Anesthesia in Mice. Anesthesiology 142, 278–297. 10.1097/ALN.0000000000005279.

74. Wang, D., Li, J., Li, H., Zhao, G., Wu, H., Zhang, X., Wang, Y., Wang, S., Li, B., Li, H., et al. (2026). A direct central amygdala-ventral tegmental area GABAergic circuit modulates anaesthesia through dopaminergic disinhibition in mice. Br J Anaesth 136, 145–157. 10.1016/j.bja.2025.07.097.

75. Campos, C.A., Bowen, A.J., Roman, C.W., and Palmiter, R.D. (2018). Encoding of danger by parabrachial CGRP neurons. Nature 555, 617–622. 10.1038/nature25511.

76. Condon, L.F., Yu, Y., Park, S., Cao, F., Pauli, J.L., Nelson, T.S., and Palmiter, R.D. (2024). Parabrachial Calca neurons drive nociplasticity. Cell Rep 43, 114057. 10.1016/j.celrep.2024.114057.

77. Sun, L., Liu, R., Guo, F., Wen, M., Ma, X., Li, K., Sun, H., Xu, C., Li, Y., Wu, M., et al. (2020). Parabrachial nucleus circuit governs neuropathic pain-like behavior. Nat Commun 11, 5974. 10.1038/s41467-020-19767-w.

78. Pal, D., and Mashour, G.A. (2022). General anesthesia and the cortical stranglehold on consciousness. Neuron 110, 1891–1893. 10.1016/j.neuron.2022.05.014.

79. Proekt, A., and Kelz, M.B. (2020). Explaining anaesthetic hysteresis with effect-site equilibration. Br J Anaesth. 10.1016/j.bja.2020.09.022.

80. Wang, Q., Zheng, Y., Lu, J., Chen, L., Wang, G.-N., and Zhou, J.-X. (2009). Isoflurane Potency in Mice from the First and Second Parity. J Am Assoc Lab Anim Sci 48, 714–717.

81. Teng, M.Z., Merenick, D., Jessel, A., Ganshorn, H., and Pang, D.S.J. (2024). Consistency in Reporting of Loss of Righting Reflex for Assessment of General Anesthesia in Rats and Mice: A Systematic Review. Comp Med 74, 12–18. 10.30802/AALAS-CM-23-000063.

82. Ramadasan-Nair, R., Hui, J., Itsara, L.S., Morgan, P.G., and Sedensky, M.M. (2019). Mitochondrial Function in Astrocytes Is Essential for Normal Emergence from Anesthesia in Mice. Anesthesiology 130, 423–434. 10.1097/ALN.0000000000002528.

83. Jiang, Q., Shao, L.-X., Yao, S., Savalia, N.K., Gilbert, A.D., Davoudian, P.A., Nothnagel, J.D., Tian, G., Hung, T.S., Lai, H.M., et al. (2026). Psilocybin triggers an activity-dependent rewiring of large-scale cortical networks. Cell 189, 659–675.e22. 10.1016/j.cell.2025.11.009.

84. Heifets, B.D., Salgado, J.S., Taylor, M.D., Hoerbelt, P., Cardozo Pinto, D.F., Steinberg, E.E., Walsh, J.J., Sze, J.Y., and Malenka, R.C. (2019). Distinct neural mechanisms for the prosocial and rewarding properties of MDMA. Science Translational Medicine 11, eaaw6435. 10.1126/scitranslmed.aaw6435.

85. Nota, M.H.C., MacMillen, L.K., Lazaro, H., Gaur, J.O., Campuzano, I., Neiswanger, C., Golden, S.A., and Heshmati, M. (2026). Activity-dependent capture reveals brain-wide signatures of isoflurane anesthesia-induced unconsciousness. Preprint at bioRxiv, 10.64898/2025.12.02.691631 https://doi.org/10.64898/2025.12.02.691631.

86. Jang, H., Fotiadis, P., Mashour, G.A., Hudetz, A.G., and Huang, Z. (2025). Role of thalamus in human conscious perception revealed by low-intensity focused ultrasound neuromodulation. Nat Commun 16, 11700. 10.1038/s41467-025-66832-3.

87. Zelmann, R., Paulk, A.C., Tian, F., Balanza Villegas, G.A., Dezha Peralta, J., Crocker, B., Cosgrove, G.R., Richardson, R.M., Williams, Z.M., Dougherty, D.D., et al. (2023). Differential cortical network engagement during states of un/consciousness in humans. Neuron 111, 3479–3495.e6. 10.1016/j.neuron.2023.08.007.

## Methods References

88. Chon, U., Vanselow, D.J., Cheng, K.C., and Kim, Y. (2019). Enhanced and unified anatomical labeling for a common mouse brain atlas. Nat Commun 10, 5067. 10.1038/s41467-019-13057-w.

89. Schneider, C.A., Rasband, W.S., and Eliceiri, K.W. (2012). NIH Image to ImageJ: 25 years of image analysis. Nat Methods 9, 671–675. 10.1038/nmeth.2089.

90. Pereira, T.D., Tabris, N., Matsliah, A., Turner, D.M., Li, J., Ravindranath, S., Papadoyannis, E.S., Normand, E., Deutsch, D.S., Wang, Z.Y., et al. (2022). SLEAP: A deep learning system for multi-animal pose tracking. Nat Methods 19, 486–495. 10.1038/s41592-022-01426-1.

91. Goodwin, N.L., Choong, J.J., Hwang, S., Pitts, K., Bloom, L., Islam, A., Zhang, Y.Y., Szelenyi, E.R., Tong, X., Newman, E.L., et al. (2024). Simple Behavioral Analysis (SimBA) as a platform for explainable machine learning in behavioral neuroscience. Nat Neurosci 27, 1411–1424. 10.1038/s41593-024-01649-9.

92. Jin, M., Ogundare, S.O., Lanio, M., Sorid, S., Whye, A.R., Leal Santos, S., Franceschini, A., and Denny, C.A. (2025). A SMARTTR workflow for multi-ensemble atlas mapping and brain-wide network analysis. eLife 13, RP101327. 10.7554/eLife.101327.3.

93. Rijsketic, D.R., Casey, A.B., Barbosa, D.A.N., Zhang, X., Hietamies, T.M., Ramirez-Ovalle, G., Pomrenze, M.B., Halpern, C.H., Williams, L.M., Malenka, R.C., et al. (2023). UNRAVELing the synergistic effects of psilocybin and environment on brain-wide immediate early gene expression in mice. Neuropsychopharmacol. 48, 1798–1807. 10.1038/s41386-023-01613-4.

94. Renier, N., Adams, E.L., Kirst, C., Wu, Z., Azevedo, R., Kohl, J., Autry, A.E., Kadiri, L., Venkataraju, K.U., Zhou, Y., et al. (2016). Mapping of Brain Activity by Automated Volume Analysis of Immediate Early Genes. Cell 165, 1789–1802. 10.1016/j.cell.2016.05.007.

95. Berg, S., Kutra, D., Kroeger, T., Straehle, C.N., Kausler, B.X., Haubold, C., Schiegg, M., Ales, J., Beier, T., Rudy, M., et al. (2019). ilastik: interactive machine learning for (bio)image analysis. Nat Methods 16, 1226–1232. 10.1038/s41592-019-0582-9.

96. Pachitariu, M., Sridhar, S., Pennington, J., and Stringer, C. (2024). Spike sorting with Kilosort4. Nat Methods 21, 914–921. 10.1038/s41592-024-02232-7.

97. Campbell, R., Blot, A., Crousseau, and Winter, O. (2020). SainsburyWellcomeCentre/lasagna: Stable IBL. Version 0.2 (Zenodo). 10.5281/ZENODO.3941894.

98. Tyson, A.L., Vélez-Fort, M., Rousseau, C.V., Cossell, L., Tsitoura, C., Lenzi, S.C., Obenhaus, H.A., Claudi, F., Branco, T., and Margrie, T.W. (2022). Accurate determination of marker location within whole-brain microscopy images. Sci Rep 12, 867. 10.1038/s41598-021-04676-9.

99. Madangopal, R., Szelenyi, E.R., Nguyen, J., Brenner, M.B., Drake, O.R., Pham, D.Q., Shekara, A., Jin, M., Choong, J.J., Heins, C., et al. (2022). Incubation of palatable food craving is associated with brain-wide neuronal activation in mice. Proceedings of the National Academy of Sciences 119, e2209382119. 10.1073/pnas.2209382119.

100. Chan, K.Y., Jang, M.J., Yoo, B.B., Greenbaum, A., Ravi, N., Wu, W.-L., Sánchez-Guardado, L., Lois, C., Mazmanian, S.K., Deverman, B.E., et al. (2017). Engineered AAVs for efficient noninvasive gene delivery to the central and peripheral nervous systems. Nat Neurosci 20, 1172–1179. 10.1038/nn.4593.

101. DeNardo, L.A., Liu, C.D., Allen, W.E., Adams, E.L., Friedmann, D., Fu, L., Guenthner, C.J., Tessier-Lavigne, M., and Luo, L. (2019). Temporal evolution of cortical ensembles promoting remote memory retrieval. Nat. Neurosci. 22, 460–469. 10.1038/s41593-018-0318-7.

102. Szelenyi, E.R., Navarrete, J.S., Murry, A.D., Zhang, Y., Girven, K.S., Kuo, L., Cline, M.M., Bernstein, M.X., Burdyniuk, M., Bowler, B., et al. (2024). An arginine-rich nuclear localization signal (ArgiNLS) strategy for streamlined image segmentation of single cells. Proceedings of the National Academy of Sciences 121, e2320250121. 10.1073/pnas.2320250121.

103. Perens, J., Salinas, C.G., Skytte, J.L., Roostalu, U., Dahl, A.B., Dyrby, T.B., Wichern, F., Barkholt, P., Vrang, N., Jelsing, J., et al. (2021). An Optimized Mouse Brain Atlas for Automated Mapping and Quantification of Neuronal Activity Using iDISCO+ and Light Sheet Fluorescence Microscopy. Neuroinformatics 19, 433–446. 10.1007/s12021-020-09490-8.

104. Yao, Z., van Velthoven, C.T.J., Kunst, M., Zhang, M., McMillen, D., Lee, C., Jung, W., Goldy, J., Abdelhak, A., Aitken, M., et al. (2023). A high-resolution transcriptomic and spatial atlas of cell types in the whole mouse brain. Nature 624, 317–332. 10.1038/s41586-023-06812-z.

105. Pauli, J.L., Chen, J.Y., Basiri, M.L., Park, S., Carter, M.E., Sanz, E., McKnight, G.S., Stuber, G.D., and Palmiter, R.D. (2022). Molecular and anatomical characterization of parabrachial neurons and their axonal projections. eLife 11, e81868. 10.7554/eLife.81868.

106. Clauset, A., Newman, M.E.J., and Moore, C. (2004). Finding community structure in very large networks. Phys. Rev. E 70, 066111. 10.1103/PhysRevE.70.066111.

107. Garrison, K.A., Scheinost, D., Finn, E.S., Shen, X., and Constable, R.T. (2015). The (in)stability of functional brain network measures across thresholds. Neuroimage 118, 651– 661. 10.1016/j.neuroimage.2015.05.046.

108. Bullmore, E., and Sporns, O. (2009). Complex brain networks: graph theoretical analysis of structural and functional systems. Nat Rev Neurosci 10, 186–198. 10.1038/nrn2575.

109. Watts, D.J., and Strogatz, S.H. (1998). Collective dynamics of ‘small-world’ networks. Nature 393, 440–442. 10.1038/30918.

110. Maslov, S., and Sneppen, K. (2002). Specificity and stability in topology of protein networks. Science 296, 910–913. 10.1126/science.1065103.

111. English, A., Uittenbogaard, F., Torrens, A., Sarroza, D., Slaven, A.V.E., Piomelli, D., Bruchas, M.R., Stella, N., and Land, B.B. (2024). A preclinical model of THC edibles that produces high-dose cannabimimetic responses. eLife 12, RP89867. 10.7554/eLife.89867.

112. Abraham, A.D., Schattauer, S.S., Reichard, K.L., Cohen, J.H., Fontaine, H.M., Song, A.J., Johnson, S.D., Land, B.B., and Chavkin, C. (2018). Estrogen Regulation of GRK2 Inactivates Kappa Opioid Receptor Signaling Mediating Analgesia, But Not Aversion. J. Neurosci. 38, 8031–8043. 10.1523/JNEUROSCI.0653-18.2018.

113. Chavkin, C., Cohen, J.H., and Land, B.B. (2019). Repeated Administration of Norbinaltorphimine Produces Cumulative Kappa Opioid Receptor Inactivation. Front. Pharmacol. 10. 10.3389/fphar.2019.00088.

114. Schattauer, S.S., Land, B.B., Reichard, K.L., Abraham, A.D., Burgeno, L.M., Kuhar, J.R., Phillips, P.E.M., Ong, S.E., and Chavkin, C. (2017). Peroxiredoxin 6 mediates Gai protein-coupled receptor inactivation by cJun kinase. Nat Commun 8, 743. 10.1038/s41467-017-00791-2.

115. Abraham, A.D., Leung, E.J.Y., Wong, B.A., Rivera, Z.M.G., Kruse, L.C., Clark, J.J., and Land, B.B. (2020). Orally consumed cannabinoids provide long-lasting relief of allodynia in a mouse model of chronic neuropathic pain. Neuropsychopharmacol. 45, 1105–1114. 10.1038/s41386-019-0585-3.

116. Hsu, Y.-W.A., Gile, J.J., Perez, J.G., Morton, G., Ben-Hamo, M., Turner, E.E., and de la Iglesia, H.O. (2017). The dorsal medial habenula minimally impacts circadian regulation of locomotor activity and sleep. J Biol Rhythms 32, 444–455. 10.1177/0748730417730169.

117. Lam, S.K., Pitrou, A., and Seibert, S. (2015). Numba: a LLVM-based Python JIT compiler. In Proceedings of the Second Workshop on the LLVM Compiler Infrastructure in HPC LLVM ’15. (Association for Computing Machinery), pp. 1–6. 10.1145/2833157.2833162.

118. Gillies, S., van der Wel, C., Van den Bossche, J., Taves, M.W., Arnott, J., Ward, B.C., and others (2024). Shapely. Version 2.0.6 (Zenodo). 10.5281/zenodo.13345370.

119. Sabnis, G., Hession, L., Mahoney, J.M., Mobley, A., Santos, M., and Kumar, V. (2024). Visual detection of seizures in mice using supervised machine learning. Preprint at bioRxiv, 10.1101/2024.05.29.596520.

120. Lopez, G.C., Camp, L.D.V., Kovaleski, R.F., Schaid, M.D., Moreno-Ramos, O.A., Awatramani, R., Cox, J.M., and Lerner, T.N. (2024). Region-specific Nucleus Accumbens Dopamine Signals Encode Distinct Aspects of Avoidance Learning. Preprint at bioRxiv, 10.1101/2024.08.28.610149.

121. Ouyang, W., Kilner, K.J., Xavier, R.M.P., Liu, Y., Lu, Y., Feller, S.M., Pitts, K.M., Wu, M., Ausra, J., Jones, I., et al. (2024). An implantable device for wireless monitoring of diverse physio-behavioral characteristics in freely behaving small animals and interacting groups. Neuron 112, 1764–1777.e5. 10.1016/j.neuron.2024.02.020.

122. Amzica, F., and Steriade, M. (1998). Electrophysiological correlates of sleep delta waves. Electroencephalogr Clin Neurophysiol 107, 69–83. 10.1016/s0013-4694(98)00051-0.

123. International Brain Laboratory (2020). Behavior: Appendix 1: IBL protocol for headbar implant surgery in mice. 10437384 Bytes. 10.6084/M9.FIGSHARE.11634726.V4.

124. International Brain Laboratory (2020). Behavior: Appendix 2: IBL protocol for mice training. 2735505 Bytes. 10.6084/M9.FIGSHARE.11634729.V3.

125. International Brain Laboratory (2021). Behavior: Appendix 3: IBL protocol for setting up the behavioral training rig. 33751033 Bytes. 10.6084/M9.FIGSHARE.11634732.V6.

126. Birman, D., Yang, K.J., West, S.J., Karsh, B., Browning, Y., Laboratory, the I.B., Siegle, J.H., and Steinmetz, N.A. (2023). Pinpoint: trajectory planning for multi-probe electrophysiology and injections in an interactive web-based 3D environment. Preprint at bioRxiv, 10.1101/2023.07.14.548952.

127. Faulkner, M. (2020). Ephys Atlas GUI. https://github.com/int-brain-lab/iblapps/tree/master/atlaselectrophysiology.

128. Hill, D.N., Mehta, S.B., and Kleinfeld, D. (2011). Quality Metrics to Accompany Spike Sorting of Extracellular Signals. Journal of Neuroscience 31, 8699–8705. 10.1523/JNEUROSCI.0971-11.2011.

129. Gosselin, E., Bagur, S., and Bathellier, B. (2024). Massive perturbation of sound representations by anesthesia in the auditory brainstem. Science Advances 10, eado2291. 10.1126/sciadv.ado2291.

130. Hildebrandt, K.J., Sahani, M., and Linden, J.F. (2017). The Impact of Anesthetic State on Spike-Sorting Success in the Cortex: A Comparison of Ketamine and Urethane Anesthesia. Front. Neural Circuits 11. 10.3389/fncir.2017.00095.

131. Buccino, A.P., Hurwitz, C.L., Garcia, S., Magland, J., Siegle, J.H., Hurwitz, R., and Hennig, M.H. (2020). SpikeInterface, a unified framework for spike sorting. eLife 9, e61834. 10.7554/eLife.61834.

